# *biocentral*: embedding-based protein predictions

**DOI:** 10.1101/2025.11.21.689449

**Authors:** Sebastian Franz, Tobias Olenyi, Paula Schloetermann, Amine Smaoui, Luisa F. Jimenez-Soto, Burkhard Rost

## Abstract

The rise of protein Language Models (pLMs) is reshaping the landscape of protein prediction. Embeddings are powerful protein representations provided by pLMs, but they come at a cost: their generation requires expensive hardware, and leveraging models often requires expert knowledge. To some extent, these hurdles limit the ease of use and benefits of those methods both for experimental and computational biologists. Biocentral aims at providing a free and open embedding-based service which addresses these challenges. We support standardized access to most pLMs currently in use, enabling researchers to generate embeddings, get embedding-based protein feature predictions, and train embedding-based models. Here, we showcase biocentral in a large-scale analysis of the BFVD virus database through *biocentral’s* predict module. We also show how readily *biocentral’s* training module reproduces an existing embedding-based prediction method. The server is accessible through a graphical user interface and a programmatic Application Programming Interface (API) at: https://biocentral.rostlab.org

## Introduction

Protein language models (*pLMs*) can generate high-dimensional vectors representing proteins. These vectors, known as *embeddings*, typically hold about 1,000 values for each residue. Recent methods effectively leverage embeddings for a variety of feature predictions, ranging from phenotype predictions such as disorder and membrane interactions, and virus-host interaction to 3D structure prediction [1-4]. The effective development and use of such methods often require expert knowledge along with access to non-standard hardware, most notably powerful GPUs with large cache and fast disk storage. This creates disparity: only well-funded labs can use the latest methods.

Over the past three decades, efforts to democratize protein prediction have led to a diverse ecosystem of computational tools. Integrated platforms such as the first online server for protein structure prediction, *PredictProtein* [5, 6] and *Phyre* [7] pioneered freely available one-stop prediction but depended on slow, MSA-based workflows. Later, workflow engines such as *Galaxy* [8] and *Nextflow* [9, 10] emphasized reproducibility and modularity, albeit at the cost of technically demanding deployments. More recently, programmatic frameworks such as *bio_embeddings* [11] have combined excellent flexibility and performance but rely on programming expertise and local computational resources. *UniProtKB* [12] has begun providing precomputed *ProtT5* [13] embeddings for its core protein database, making these large-scale, standardized protein representations broadly accessible.

*biocentral* leverages modern pLMs to provide fast, embedding-based predictions through a web interface and an Application Programming Interface (API), building upon the foundational vision of *PredictProtein*. By combining the accessibility of integrated servers with the scalability of programmatic frameworks, it aims to enable expert-centric pipelines and open, reproducible access to state-of-the-art protein embeddings and predictions. The goal of *biocentral* is to make computational tools more accessible to biomedical researchers by reducing technical barriers to the latest models and methods, thus allowing these domain experts to really focus on results interpretation and scientific discovery.

We demonstrate the application of *biocentral* through two examples. Firstly, we use the *biocentral* prediction module to predict transmembrane regions through *TMbed* [2] based on *ProtT5* for all sequences in the *BFVD* virus database [14] with the goal to update our existing *TMvisDB* resource [15]. Secondly, we leverage the embedding and training modules of *biocentral* to reproduce a recently published method using embeddings to identify bacterial exotoxins from sequence, namely *ExoTox* [16]. The *ExoTox* model is available as part of the prediction module of *biocentral*.

*biocentral* is available at https://biocentral.rostlab.org. For ease-of-use in coding environments, we provide an API package (for *Python* and *Dart*) at https://github.com/biocentral/biocentral_api. The service repository can be found at https://github.com/biocentral/biocentral_server. A frontend is constantly being developed and available at https://app.biocentral.cloud.

## Methods

### Service architecture

Our lab has a long-standing history of developing publicly accessible protein prediction services [5, 17]. The transition toward embedding-based methods required rethinking these systems in light of new computational and maintenance challenges [11]. Two core issues emerged: (1) fragile software dependencies that hindered long-term maintainability and (2) complex coordination of costly resources such as GPUs, databases, and API gateways.

To address the first issue, *biocentral* introduced a modular, distributed architecture built on containerized services. Each functional component runs independently via *docker-compose*, enabling deployment across machines and independent scaling. Embedding caching reduces redundant GPU computation, while an *Open Neural Network Exchange* (*ONNX*)-based [18] workflow standardizes model integration and reduces dependency conflicts. This pipeline allows leveraging the *Nvidia Triton* inference server for optimized and distributed model inference [19]. Our architecture simplifies sustainable operation, maintenance, and rapid incorporation of new resources like pLMs and prediction methods.

### Core modules

#### Embedding module

The embedding module forms the computational core of *biocentral*, providing per-residue (per-token) and per-protein (pooled) pLM embeddings for any compatible transformer model hosted on *HuggingFace* [20]. Embedding generation is implemented via the *biotrainer* framework [21], with additional support for *ONNX*-optimized versions of commonly used pLMs such as *ProtT5* [13] and *ESM-2* (8M, 3B) [3]. These optimized models are executed through the *NVIDIA Triton* inference server for optimal resource usage [22].

To minimize redundant computation, generated embeddings are cached in a *PostgreSQL* database indexed by a sequence hash and the model identifier. Caching embeddings uses up a lot of disk space (roughly 1 GB for 1,000 sequences for per-residue *ProtT5* embeddings). That is why the database is cleaned up regularly following a first-in-first-out principle to have the latest used embeddings available.

*ProtSpace* [23] visualizes embedding analysis. Beyond embedding generation, the *biocentral* module can also inter-operate with the *ProtSpace* library.

#### Prediction module

Building on the embedding module, *biocentral’s* prediction module integrates several established models for different protein prediction tasks (Table 1). Each model is included as a lightweight interface that defines input and output transformations, records metadata such as the underlying pLM, and links to the corresponding optimized *ONNX/Triton* model. The wrapper itself contains no machine-learning logic; the latter is entirely handled by the *ONNX* runtime. The exported *ONNX* model, derived from the original architecture and pre-trained weights, can be deployed independently via *Triton*.

**Table 1:**
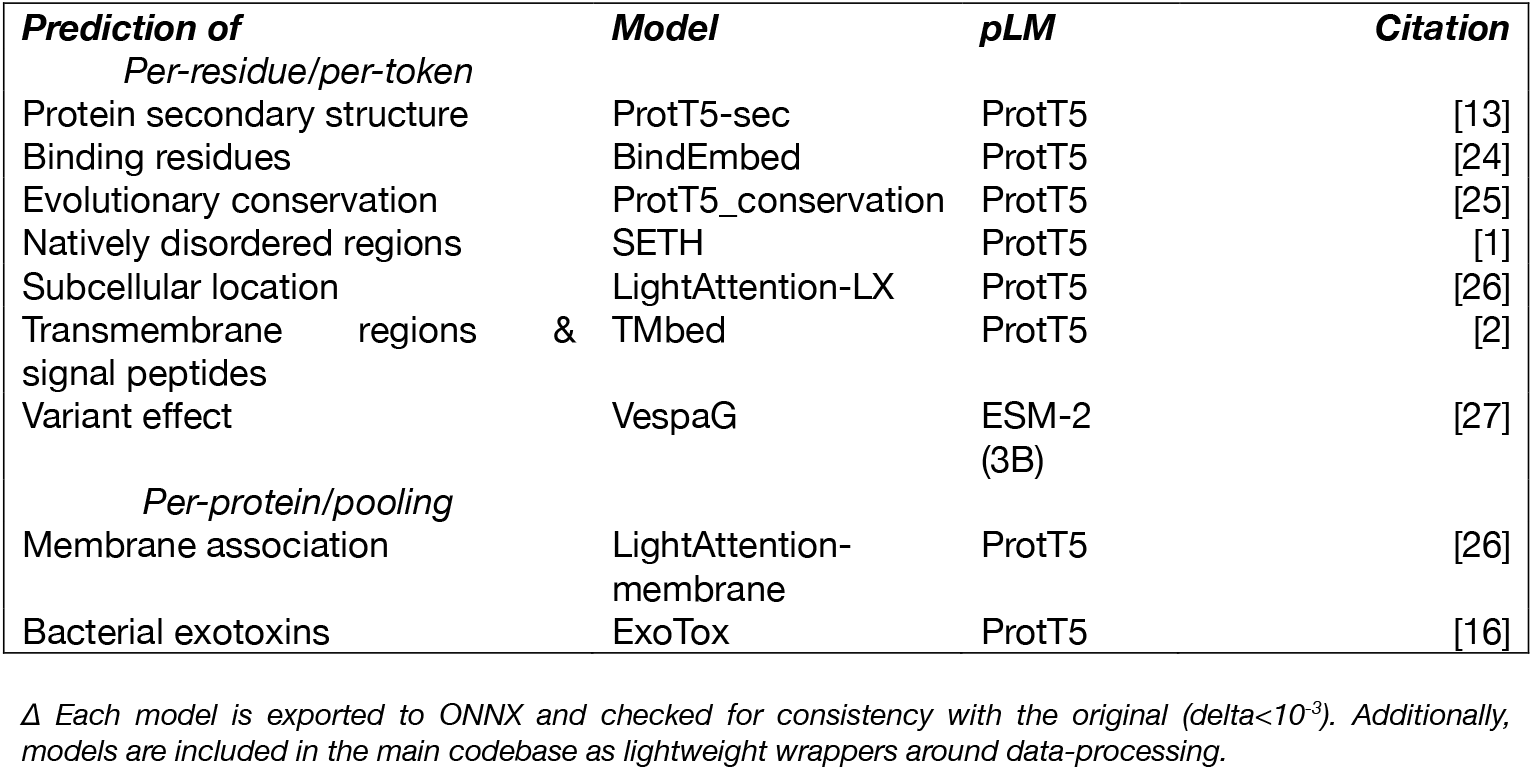
Prediction models in *biocentral* ^Δ^.

| *Prediction of* | *Model* | *pLM* | *Citation* |
| --- | --- | --- | --- |
| *Per-residue/per-token* |  |  |  |
| Protein secondary structure | ProtT5-sec | ProtT5 | [13] |
| Binding residues | BindEmbed | ProtT5 | [24] |
| Evolutionary conservation | ProtT5_conservation | ProtT5 | [25] |
| Natively disordered regions | SETH | ProtT5 | [1] |
| Subcellular location | LightAttention-LX | ProtT5 | [26] |
| Transmembrane regions & signal peptides | TMbed | ProtT5 | [2] |
| Variant effect | VespaG | ESM-2 (3B) | [27] |
| *Per-protein/pooling* |  |  |  |
| Membrane association | LightAttention-membrane | ProtT5 | [26] |
| Bacterial exotoxins | ExoTox | ProtT5 | [16] |

<sup>Δ</sup> Each model is exported to ONNX and checked for consistency with the original ( $\Delta < 10^{-3}$ ). Additionally, models are included in the main codebase as lightweight wrappers around data-processing.

This modular workflow enables rapid integration of new methods into *biocentral* without adding additional dependencies or computational overhead. The current release includes a broad range of models (Table 1), which we continuously expand both internally and in collaboration with model developers (SOM; Algorithm 1).

#### Model training

The training module allows to develop new supervised prediction models based on *pLM* embeddings. The module is implemented using *biotrainer* [21]. Input parameters are provided as a *biotrainer* configuration file. Before training, the config options are validated for internal consistency. If available, embeddings are loaded from the internal *PostgreSQL* database; otherwise, they are computed on the fly. After successful completion, model checkpoint(s) and a summary file are returned. This allows for full reproducibility and control over the training process and results. *Biotrainer* provides several sanity checks and test set baselines out-of-the-box, that allow for a quick initial evaluation of the training process and model quality. The module enables rapid prototyping of biomedical prediction models based on embeddings and will be extended in the future.

### Soft- and hardware

#### Software stack

The server is implemented in Python 3.12 using the *FastAPI* web framework [28]. Deployment is handled via *docker-compose*, allowing for the orchestration of multiple containers that may even run on different machines. This setup enables local deployment for advanced users with sufficient hardware. The programmatic API was generated using *openapi-generator* with the *OpenAPI* specifications [29] from the *FastAPI* framework. To handle the underlying complexity, we provide a stable high-level interface exposing core functions such as *train, embed* and *predict* for straightforward programmatic use. An early version graphical user interface (GUI) is available and being improved. It is built using *Dart/Flutter* supporting all common desktop platforms and web browsers.

#### Deployment hardware

The *biocentral* service is currently deployed on a DELL PowerEdge R7525 with two AMD 7313 (32 thread) processors, 256GB RAM, and an NVIDIA A10 (24GB) GPU.

### Data sets for case studies

#### Big Fantastic Virus Database (BFVD)

The *Big Fantastic Virus Database (BFVD)* is a comprehensive repository holding 351,242 predicted viral protein structures derived from UniRef30 clusters of virus proteins [14]. BFVD structures are predicted by *ColabFold* [30] and subsequently refined by integrating homologous sequences from a large assembled-sequencing data set. *BFVD* represents a highly diverse set of viral structures: over 60% of have little or no structural similarity to AFDB (the database with *AlphaFold*2 predictions of protein structure [31]) or to the experimental 3D structures from the *PDB* [32].

#### ExoTox

*ExoTox* is a machine-learning model predicting bacterial exotoxins from sequences of secreted non-toxic bacterial proteins [16]. The method has been trained on a curated dataset of 1,069 bacterial exotoxins and 1,308 secreted non-toxins derived from *UniProtKB/Swiss-Prot* [12] and *PSORTb 3*.*0b* [33]. Redundancy was reduced to ≤30% pairwise sequence identity (PIDE) using *MMseqs2* [34] to ensure non-overlapping training and test sets, i.e., no pair in the data set had >30% PIDE. *ExoTox* inputs were the first twenty principal components of pooled (per-protein averaged) *ProtT5* embeddings. All models were evaluated using the Matthew’s Correlation Coefficient (MCC), calculated using Equation 1.

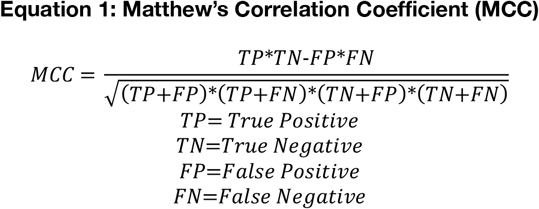

A Support Vector Machine (SVM) [35] trained on these embeddings performs well in distinguishing exotoxins from secreted controls (MCC≈0.94 [16]). It outperforms both sequence-based (*BLAST* [36]) and generalized toxin prediction methods.

## Results and Discussion

### Predicting for large datasets: membrane proteins in BVFD

To explore the applicability of *biocentral* for large-scale dataset analysis, we used *Tmbed* [2] to predict transmembrane topology across all predicted viral proteins from *BFVD* [14]. After a basic filtering step, we submitted the dataset of 345,141 sequences in seven batches of around 50,000 proteins. In total, the computation took around two days.

*TMbed* predicted transmembrane regions in approximately 11.6% of all investigated *BFVD* entries. To examine whether these predictions reflect biologically meaningful trends, we compared the results between enveloped and non-enveloped viruses. Aligning with biological expectation, proteins from enveloped viruses were predicted to more frequently contain transmembrane regions [37] than those from non-enveloped viruses (Table 2). While these observations remain illustrative, they support *TMbed*’s applicability to viruses.

**Table 2:**
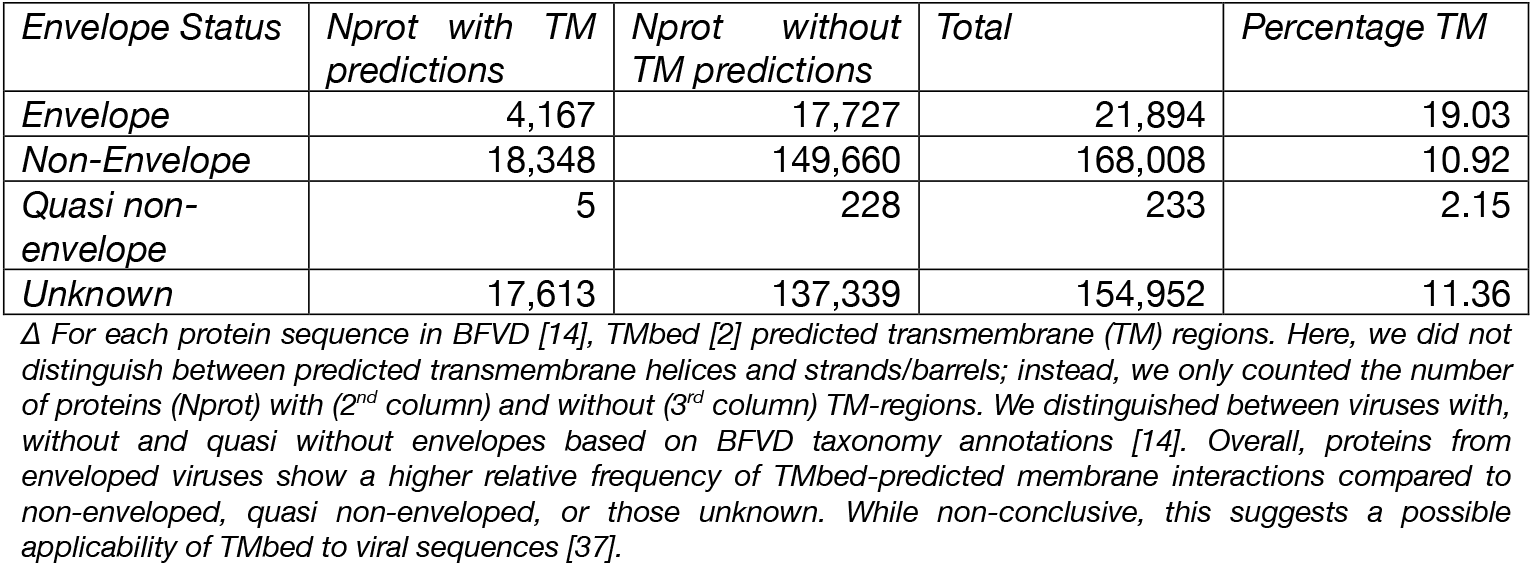
TMbed predictions on BFVD ^Δ^.

| *Envelope Status* | *Nprot with TM predictions* | *Nprot without TM predictions* | *Total* | *Percentage TM* |
| --- | --- | --- | --- | --- |
| *Envelope* | 4,167 | 17,727 | 21,894 | 19.03 |
| *Non-Envelope* | 18,348 | 149,660 | 168,008 | 10.92 |
| *Quasi non-envelope* | 5 | 228 | 233 | 2.15 |
| *Unknown* | 17,613 | 137,339 | 154,952 | 11.36 |

<sup>Δ</sup> For each protein sequence in BFVD [14], TMbed [2] predicted transmembrane (TM) regions. Here, we did not distinguish between predicted transmembrane helices and strands/barrels; instead, we only counted the number of proteins (Nprot) with (2<sup>nd</sup> column) and without (3<sup>rd</sup> column) TM-regions. We distinguished between viruses with, without and quasi without envelopes based on BFVD taxonomy annotations [14]. Overall, proteins from enveloped viruses show a higher relative frequency of TMbed-predicted membrane interactions compared to non-enveloped, quasi non-enveloped, or those unknown. While non-conclusive, this suggests a possible applicability of TMbed to viral sequences [37].

We further examined this by analyzing sequences predicted to contain both alpha-helical (TMH) and beta-barrel (TMB) regions—a combination unlikely to exist in nature and previously used as a lower bound for incorrect predictions [2]. Out of all 345,141 viral proteins, 43 proteins (<0.2‰) were predicted with such mistakes. Only 19 of all proteins with predicted TM regions had high-confidence 3D structure predictions (pLDDT>70 provided by BFVD). We examined the shortest protein (A0A516M0S4; 54 residues; average pLDDT ≈ 88) and the longest protein (A0A6J5RKD4; 339 residues; average pLDDT ≈ 85) (SOM; Figure 1). While the irregular pattern of predicted beta-barrel residues suggests a mistake, the structure resembles similar pore-forming structures [39], hinting at the predictor’s confusion.

**Figure 1:**
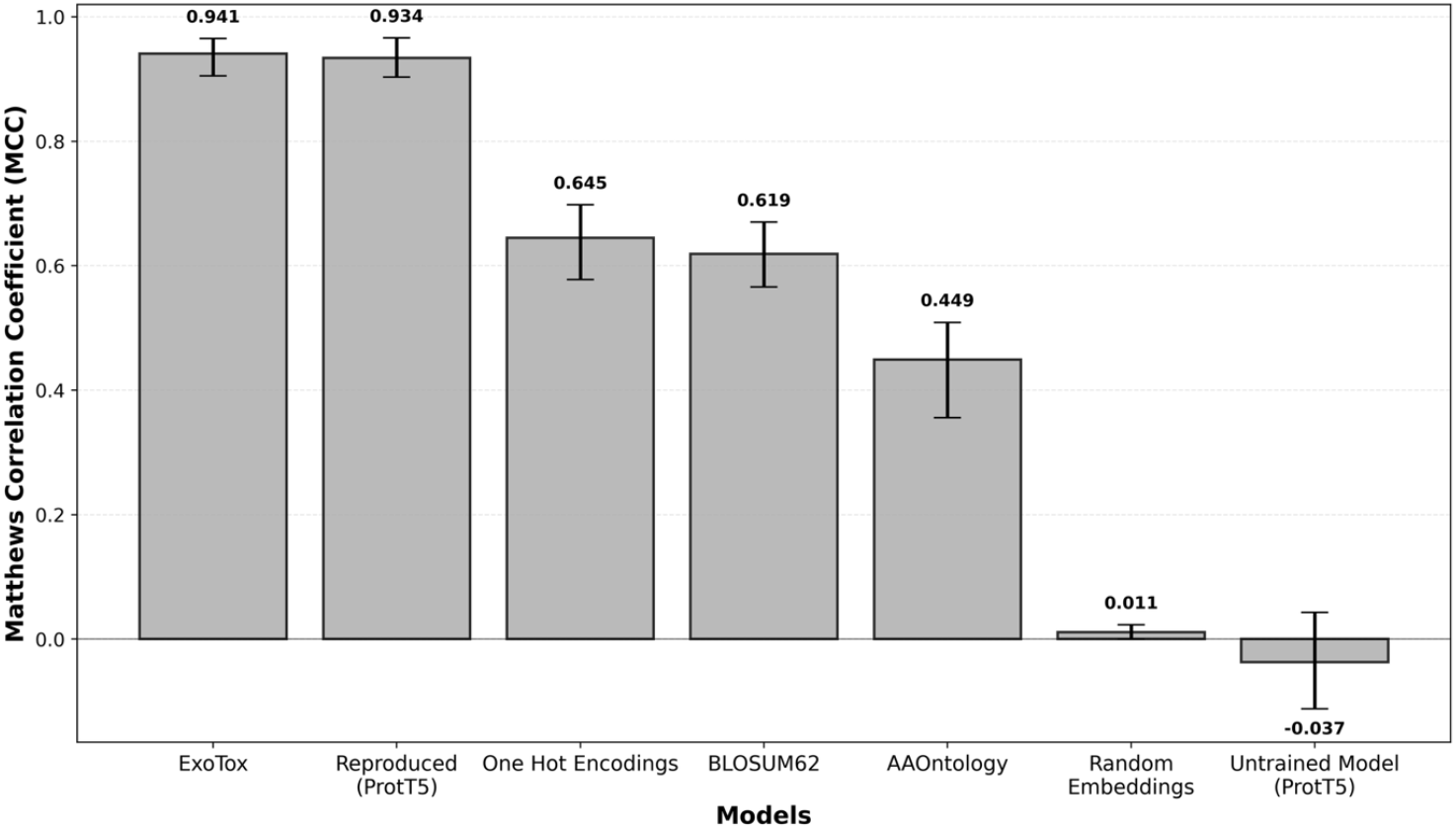
Original ExoTox compared to baseline and *biocentral* reproduced FNN ^Δ^. Δ Comparing MCC of the original ExoTox predictor [16] (SVM) against the reproduced FNN and baseline encodings. The reproduced model matches the original’s performance closely, indicating that the biocentral training module can reproduce a published biological predictor. Both ProtT5-based models outperform the common baselines of one-hot encoded sequences and BLOSUM62 [40] (encoding an amino acid by its respective substitution vector) as well as AAOntology, where an amino acid is encoded by its respective values from 568 physicochemical scales [41]. All these models and baselines outperform random performance, empirically evaluated using randomly generated embedding vectors (dimension: 128) as input, and a randomly initialized, untrained model using ProtT5 embeddings.

All membrane predictions on the *BVFD* facilitated through *biocentral* are now integrated into *TMVisDB* [15], extending its coverage to viral proteins. While TMbed predictions on BFVD follow biological intuitions, results should be treated with caution due to the absence of viral sequences in the training of TMbed [2]. We hope that the availability of viral trans-membrane predictions can lead to new insights for future model development and unravel crucial differences in viral membrane interactions.

### Training embedding-based models from scratch: reproducing ExoTox

Exotoxins produced by bacteria are proteins released into the extracellular space and can affect target cells. Precise detection of these toxins can improve pharmaceutical development and guarantee the security of medical treatments that utilize bacteria. The recently released *ExoTox* prediction method [16] inputs *ProtT5* embeddings (reduced to the largest 20 principal components) and is trained on a curated subset of bacterial exotoxins and secreted non-toxic proteins. Using *biocentral*, we reproduce this method as evidenced by reaching similar performance on the same test set (MCC≈0.93, Equation 1) as a feed-forward neural network (FNN) (compared to the original SVM), while outperforming all evaluated baselines (Figure 1). Creating this model only requires providing annotated sequences (with labels and sets) before calling the biocentral training module. This reduces initial model development time without prior programming knowledge from weeks or months to days or even hours. The results also show that models trained via *biocentral* can serve as a control group or sanity check for more sophisticated, specialized architectures.

We compiled a complete guide of the steps required for reproduction in the supplementary online material (SOM; Algorithm 2). The original ExoTox SVM was converted to *ONNX* and is available as a predictor in the *biocentral* prediction module.

### Limitations and Future Work

The *biocentral* project will continue to evolve. Maintainability and longevity remain central goals. To strengthen portability and sustainability, we plan to explore advanced deployment methods, such as Kubernetes-based orchestration. This would simplify deployment across diverse cloud environments and raise resilience to changes in available computational resources and funding.

Beyond maintaining and expanding the current ecosystem, we aim to extend *biocentral* towards the emerging field of protein design and engineering, broadening access to state-of-the-art generative and optimization tools. We also plan to further emphasize training and fine-tuning of existing models to make machine learning more accessible. We hope to expand the catalogue of models through community engagement, encouraging model developers to contribute and collaborate on shared standards of accessibility and interpretability.

Our case study of *TMbed* demonstrated that large-scale predictions through *biocentral* can help to explore model behavior for sequences that deviate strongly from the training distribution. To further aid interpretation, we plan to develop model-specific applicability domain layers that document strengths and weaknesses, and guide users toward reasonable use of predictive models. Thus, a collaboration between domain experts applying prediction models to their field of research and model developers can speed up model development in the future.

The reproduction of the *ExoTox* method exemplified rapid prototyping of prediction models on biological data through *biocentral*. This can significantly lower the barriers for biomedical researchers to create custom deep learning solutions. However, reducing the required technological expertise to create such models has the potential to decrease the ability to interpret the model’s behavior and results. We plan to address this by extending the provided frontend with utilities for model comparison and explainability, supporting the interpretation process for domain experts.

## Conclusion

We introduced *biocentral*, a freely available service that makes state-of-the-art, embedding-based protein predictions accessible to a broad community. Through its modular architecture, standardized *ONNX* model format, and *Triton*-based inference, the server offers a reproducible and sustainable framework for integrating and deploying protein prediction models. The two case studies demonstrate both its scalability for large datasets and its capacity to reproduce existing embedding-based methods with minimal effort.

By abstracting the complexity of machine-learning pipelines into a stable programmatic interface, *biocentral* establishes a pattern that others can adopt to publish and maintain their models in a unified, interoperable format. In this way, it extends the long-standing vision of democratizing protein prediction, removing barriers between method developers and end-users, and creating a common platform that evolves with the advancing field of protein language models.

## Abbreviations

3D: three-dimensional
API: Application Programming Interface
BFVD: Big Fantastic Virus Database
BLAST: Basic Local Alignment Search Tool
FNN: Feedforward Neural Network
GPU: Graphics Processing Unit
MCC: Matthews Correlation Coefficient
MSA: Multiple Sequence Alignment
NCBI: National Center for Biotechnology Information
ONNX: Open Neural Network Exchange
pLDDT: predicted local distance difference test
pLM: protein Language Model
SVM: Support Vector Machine
UniProtKB: Universal Protein Resource Knowledgebase (database).

## Availability and requirements

**Project name:** biocentral

**Project home page:** https://github.com/biocentral/Operatingsystem(s): Platform independent

**Programming language: Python 3.12**

**Other requirements:**

○ Redis 7 (alpine)
○ PostgreSQL 15
○ SeaweedFS (chrislusf/seaweedfs)
○ NVIDIA Triton Inference Server 24.12
○ Prometheus (prom/prometheus)
○ taxoniq (≥1.0.1, <2.0.0)
○ blosc2 (≥2.7.1, <3.0.0)
○ psycopg[binary] (≥3.2.3, <4.0.0)
○ rq (≥2.1.0, <3.0.0)
○ python-dotenv (≥1.2.1, <2.0.0)
○ gpytorch (≥1.14, <2.0)
○ onnxruntime-gpu (≥1.16.0, <2.0.0; Linux only)
○ onnxruntime (≥1.16.0, <2.0.0; non-Linux platforms)
○ tritonclient[grpc] (≥2.61.0)
○ circuitbreaker (≥2.0.0)
○ httpx (≥0.28.1)
○ pydantic (≥2.12.4)
○ fastapi (≥0.121.0)
○ uvicorn (≥0.38.0)
○ prometheus-client (≥0.20.0)
○ prometheus-fastapi-instrumentator (≥6.1.0)
○ fastapi-limiter (≥0.1.6)
○ biotrainer (github.com/biocentral/biotrainer.git@fix/api-improvements)
○ protspace (github.com/tsenoner/protspace.git@v3.1.1)
○ hvi_toolkit (github.com/SebieF/hvi_toolkit.git@maintenance/update-biotrainer)
○ vespag (github.com/biocentral/VespaG.git@onnx-deployment)
○ tmbed (github.com/biocentral/TMbed.git@onnx-deployment)

**Hardware requirements:** NVIDIA GPU with CUDA support (recommended: ≥16GB VRAM for full model deployment)

**License:** GPL-3.0

**Any restrictions on use by non-academics:** None

## Declarations

### Availability of data and materials

The code for the biocentral service is available at https://github.com/biocentral/biocentral_server, the API client’s code at https://github.com/biocentral/biocentral_api, both open source under GPL-3.0. The API is available at https://biocentral.rostlab.org/docs. The frontend is available at https://app.biocentral.cloud. The Big Fantastic Virus Database (BFVD) is available under https://bfvd.steineggerlab.workers.dev/. The ExoTox dataset is available here: https://data.ub.uni-muenchen.de/576/. The scripts for creating the analysis of the BFVD TMbed predictions and the reproduction of the ExoTox predictor is available in the supplementary materials. The latter was also published as an example for the usage of the biocentral API at https://github.com/biocentral/biocentral_api/tree/main/python/examples/exotox_reproduction.

### Authors’ contributions

S.F. conceptualized the project; S.F. and T.O developed software and methodology, with P.S. contributing to software and model conversion. S.F. and T.O performed analyses with A.S. contributing data and L. JS. contributing data and expertise. S.F. and T.O wrote the initial manuscript with all authors contributing. B.R. provided supervision and acquired funding. All authors read and approved the final manuscript.

## Acknowledgments

We thank Nikita Kugut (TUM) for his support with many aspects of this work. We thank the Technical University of Munich (TUM) for providing facilities and resources. Also, thanks to the anonymous reviewers and the editor for improving our work. Finally, we thank those who deposit experimental data in public databases, maintain these databases, and develop methods to enrich experimental data.

*Declaration of generative AI and AI-assisted technologies in the writing process:*

While preparing this work, the authors used Claude.ai and ChatGPT.com to check the language and grammar of this manuscript. After using these services, the authors reviewed and edited the content as needed and take full responsibility for the content of the published article.

## Ethics approval and consent to participate

Not applicable

### Consent for publication

Not applicable

