## Supplementary material for "*biocentral*: embedding-based protein predictions": enrich_bfvd_from_local_taxonomy.py

```
#!/usr/bin/env python3
"""
enrich_bfvd_from_local_taxonomy.py

Enrich a BFVD/TMbed-style protein CSV with taxonomy using ONLY the bfvd taxonomy metadata file:

    bfvd_taxid_rank_scientificname_lineage.tsv

Expected TSV format (tab-separated, NO header):

    col0: PDB filename
          e.g. A0A514CX81_unrelaxed_rank_001_alphafold2_ptm_model_3_seed_000.pdb
    col1: TaxID
          e.g. 2588489
    col2: Rank
          e.g. "no rank"
    col3: Scientific name
          e.g. "Gordonia phage Dardanus"
    col4: Lineage
          e.g. "d_Viruses;-_Duplodnaviria;k_Heunggongvirae;..."

UniProt accession is taken as the part of the PDB filename BEFORE the first "_":
    "A0A514CX81_unrelaxed_..." -> "A0A514CX81"

We then enrich the input CSV (which must contain "uniprot_accession") with:

    organism_name  (from TSV scientific_name)
    domain         (Bacteria / Archaea / Eukaryota / unclassified sequence)
    kingdom        (from k_... in lineage, else "Unknown")
    taxid          (from TSV)
    lineage        (full lineage string from TSV)

Output CSV columns:

    uniprot_accession
    length
    tm_class
    has_signal_peptide
    num_TMH
    num_TMB
    organism_name
    domain
    kingdom
    taxid
    lineage

    Usage:
  python enrich_bfvd_from_local_taxonomy.py INPUT.csv bfvd_taxid_rank_scientificname_lineage.tsv -o OUTPUT.csv

INPUT.csv  = your BFVD/TMbed predictions (must have 'uniprot_accession' etc.)
TSV        = BFVD taxonomy file with PDB filename, taxid, rank, scientific_name, lineage
OUTPUT.csv = same proteins + added columns: organism_name, domain, kingdom, taxid, lineage

"""

import argparse
import csv
import logging
import os
from typing import Dict, Optional


def setup_logging(verbose: bool = False) -> None:
    level = logging.DEBUG if verbose else logging.INFO
    logging.basicConfig(
        level=level,
        format="%(asctime)s [%(levelname)s] %(message)s",
        datefmt="%Y-%m-%d %H:%M:%S",
    )


def extract_base_accession(value: str) -> str:
    """
    Take the part before the first underscore, e.g.:

        "A0A514CX81_unrelaxed_rank_001..." -> "A0A514CX81"
        "A0A023MIL2_A"                    -> "A0A023MIL2"
    """
    if not value:
        return ""
    return value.split("_", 1)[0].strip()


def detect_csv_dialect(path: str, sample_size: int = 4096) -> csv.Dialect:
    with open(path, "r", encoding="utf-8", newline="") as fh:
        sample = fh.read(sample_size)
        fh.seek(0)
        sniffer = csv.Sniffer()
        try:
            dialect = sniffer.sniff(sample)
        except csv.Error:
            logging.warning("Could not detect CSV dialect, defaulting to comma delimiter.")

            class SimpleDialect(csv.excel):
                delimiter = ","

            dialect = SimpleDialect()
    logging.info("Detected delimiter: %r", dialect.delimiter)
    return dialect


def classify_domain_from_lineage(lineage: str) -> str:
    """
    Classify domain into one of:
        - Bacteria
        - Archaea
        - Eukaryota
        - unclassified sequence  (for viruses or anything else)

    BFVD lineage looks like:
        d_Viruses;-_Duplodnaviria;k_Heunggongvirae;...

    We grab the token starting with "d_" (if any) and inspect it.
    """
    if not lineage:
        return "unclassified sequence"

    tokens = [t.strip() for t in lineage.split(";") if t.strip()]
    for token in tokens:
        if token.startswith("d_"):
            dom_value = token[2:].strip()  # remove "d_"
            dom_cf = dom_value.casefold()
            if dom_cf.startswith("bacteria"):
                return "Bacteria"
            if dom_cf.startswith("archaea"):
                return "Archaea"
            if dom_cf.startswith("eukaryota"):
                return "Eukaryota"
            # Viruses or anything else: fall through to unclassified sequence
            return "unclassified sequence"

    return "unclassified sequence"


def classify_kingdom_from_lineage(lineage: str) -> str:
    """
    Extract a "kingdom-like" string from BFVD lineage.

    BFVD lineage looks like:
        d_Viruses;-_Duplodnaviria;k_Heunggongvirae;...

    We take the token starting with "k_" (if present) and strip the prefix.
    Otherwise, return "Unknown".
    """
    if not lineage:
        return "Unknown"

    tokens = [t.strip() for t in lineage.split(";") if t.strip()]
    for token in tokens:
        if token.startswith("k_"):
            return token[2:].strip() or "Unknown"

    return "Unknown"


def load_bfvd_taxonomy_tsv(tsv_path: str) -> Dict[str, dict]:
    """
    Load bfvd_taxid_rank_scientificname_lineage.tsv and build a dictionary:

        base_accession -> {
            "organism_name": scientific_name,
            "taxid": taxid,
            "lineage": lineage,
            "domain": domain,
            "kingdom": kingdom,
        }

    If multiple PDB entries map to the same accession, the first one wins.
    """
    mapping: Dict[str, dict] = {}
    logging.info("Loading BFVD taxonomy from %s ...", tsv_path)

    with open(tsv_path, "r", encoding="utf-8") as fh:
        for line_no, line in enumerate(fh, start=1):
            line = line.rstrip("\n")
            if not line:
                continue
            parts = line.split("\t")
            if len(parts) < 5:
                logging.warning("Skipping malformed TSV line %d: %r", line_no, line)
                continue

            pdb_filename, taxid, rank, scientific_name, lineage = parts[:5]

            base_acc = extract_base_accession(pdb_filename)
            if not base_acc:
                logging.warning(
                    "Could not extract accession from PDB filename %r at line %d; skipping.",
                    pdb_filename,
                    line_no,
                )
                continue

            if base_acc in mapping:
                # Already have an entry for this accession; keep the first one.
                continue

            lineage = lineage.strip() if lineage else ""
            domain = classify_domain_from_lineage(lineage)
            kingdom = classify_kingdom_from_lineage(lineage)

            mapping[base_acc] = {
                "organism_name": scientific_name.strip() if scientific_name else "",
                "taxid": taxid.strip() if taxid else "",
                "lineage": lineage,
                "domain": domain,
                "kingdom": kingdom,
            }

    logging.info("Loaded taxonomy for %d unique UniProt accessions from TSV.", len(mapping))
    return mapping


def enrich_csv_with_local_taxonomy(
    input_csv: str,
    taxonomy_tsv: str,
    output_csv: str,
) -> None:
    # Load local mapping
    taxonomy_map = load_bfvd_taxonomy_tsv(taxonomy_tsv)

    # Read input CSV, detect dialect
    dialect = detect_csv_dialect(input_csv)

    logging.info("Reading input CSV %s and writing enriched CSV %s ...", input_csv, output_csv)

    with open(input_csv, "r", encoding="utf-8", newline="") as in_fh, open(
        output_csv,
        "w",
        encoding="utf-8",
        newline="",
    ) as out_fh:
        reader = csv.DictReader(in_fh, dialect=dialect)

        required_cols = [
            "uniprot_accession",
            "length",
            "tm_class",
            "has_signal_peptide",
            "num_TMH",
            "num_TMB",
        ]
        missing = [c for c in required_cols if c not in (reader.fieldnames or [])]
        if missing:
            logging.error(
                "Input CSV is missing required columns %s. Available columns: %s",
                missing,
                reader.fieldnames,
            )
            raise SystemExit(1)

        # Output columns: includes taxid & lineage, no reviewed / PE
        fieldnames = [
            "uniprot_accession",
            "length",
            "tm_class",
            "has_signal_peptide",
            "num_TMH",
            "num_TMB",
            "organism_name",
            "domain",
            "kingdom",
            "taxid",
            "lineage",
        ]
        writer = csv.DictWriter(out_fh, fieldnames=fieldnames)
        writer.writeheader()

        total_rows = 0
        enriched_rows = 0
        missing_tax_rows = 0

        for row in reader:
            total_rows += 1

            acc_value = row.get("uniprot_accession", "").strip()
            base_acc = extract_base_accession(acc_value)
            info: Optional[dict] = taxonomy_map.get(base_acc)

            if info is None:
                missing_tax_rows += 1
                organism_name = ""
                domain = "unclassified sequence"
                kingdom = "Unknown"
                taxid = ""
                lineage = ""
            else:
                enriched_rows += 1
                organism_name = info["organism_name"]
                domain = info["domain"]
                kingdom = info["kingdom"]
                taxid = info["taxid"]
                lineage = info["lineage"]

            out_row = {
                "uniprot_accession": row.get("uniprot_accession", ""),
                "length": row.get("length", ""),
                "tm_class": row.get("tm_class", ""),
                "has_signal_peptide": row.get("has_signal_peptide", ""),
                "num_TMH": row.get("num_TMH", ""),
                "num_TMB": row.get("num_TMB", ""),
                "organism_name": organism_name,
                "domain": domain,
                "kingdom": kingdom,
                "taxid": taxid,
                "lineage": lineage,
            }
            writer.writerow(out_row)

    logging.info("Finished. Total rows: %d", total_rows)
    logging.info("Rows enriched from TSV: %d", enriched_rows)
    if missing_tax_rows:
        logging.warning(
            "No local taxonomy found for %d rows "
            "(organism_name empty, domain=unclassified sequence, kingdom=Unknown, taxid/lineage empty).",
            missing_tax_rows,
        )


def parse_args():
    parser = argparse.ArgumentParser(
        description="Enrich BFVD/TMbed CSV with taxonomy using bfvd_taxid_rank_scientificname_lineage.tsv only.",
        formatter_class=argparse.ArgumentDefaultsHelpFormatter,
    )
    parser.add_argument("input_csv", help="Input CSV file with BFVD/TMbed predictions.")
    parser.add_argument(
        "taxonomy_tsv",
        help="bfvd_taxid_rank_scientificname_lineage.tsv file "
             "(PDB filename, taxid, rank, scientific_name, lineage).",
    )
    parser.add_argument(
        "-o",
        "--output",
        help="Output CSV file. If not provided, will append '.local_taxonomy.csv' to the input filename.",
    )
    parser.add_argument(
        "-v",
        "--verbose",
        action="store_true",
        help="Enable verbose (DEBUG-level) logging.",
    )
    return parser.parse_args()


def main():
    args = parse_args()
    setup_logging(args.verbose)

    input_csv = args.input_csv
    taxonomy_tsv = args.taxonomy_tsv
    output_csv = args.output or (os.path.splitext(input_csv)[0] + ".local_taxonomy.csv")

    logging.info("Input CSV:    %s", input_csv)
    logging.info("Taxonomy TSV: %s", taxonomy_tsv)
    logging.info("Output CSV:   %s", output_csv)

    enrich_csv_with_local_taxonomy(
        input_csv=input_csv,
        taxonomy_tsv=taxonomy_tsv,
        output_csv=output_csv,
    )


if __name__ == "__main__":
    main()
```
