## Supplementary material for "*biocentral*: embedding-based protein predictions": enrich_bfvd_with_taxonomy.py

```
#!/usr/bin/env python
"""
DICLAIMER: This script might take a while to run, the api requests to uniprot are rate-limited.

Join a BFVD TMbed CSV with bfvd_taxid.tsv and enrich with UniProt taxonomy metadata
(organism_name, reviewed, protein_existence).

Usage example
-------------
python enrich_bfvd_with_taxonomy.py \
    --input shards_merged.csv \
    --bfvd-taxid bfvd_taxid.tsv \
    --output bfvd_with_taxonomy.csv
"""

import argparse
import os
import sys
import time
from typing import Dict, Iterable, List

import pandas as pd
import requests

UNIPROT_SEARCH_URL = "https://rest.uniprot.org/uniprotkb/search"

# Map UniProt protein existence descriptions to numeric PE levels
PE_LEVELS = {
    "evidence at protein level": 1,
    "evidence at transcript level": 2,
    "inferred from homology": 3,
    "predicted": 4,
    "uncertain": 5,
}


def chunked(iterable: Iterable[str], size: int) -> Iterable[List[str]]:
    """Yield lists of at most `size` elements from `iterable`."""
    chunk: List[str] = []
    for item in iterable:
        chunk.append(item)
        if len(chunk) >= size:
            yield chunk
            chunk = []
    if chunk:
        yield chunk


def normalise_header_name(name: str) -> str:
    """
    Normalise TSV header names from UniProt to a simpler, lowercase form.

    E.g. 'Entry' -> 'accession', 'Organism' -> 'organism_name', etc.
    """
    n = name.strip().lower()
    n = n.replace(" ", "_")
    if n in ("entry", "accession"):
        return "accession"
    if n in ("organism", "organism_name"):
        return "organism_name"
    if n in ("status", "reviewed"):
        return "reviewed"
    if n in ("protein_existence", "existence", "protein_existence_pe"):
        return "protein_existence"
    return n


def format_protein_existence(pe_value: str) -> str:
    """
    Turn a UniProt 'protein_existence' string into 'N: Description' form
    if possible, otherwise return as-is.

    Example:
        'Inferred from homology' -> '3: Inferred from homology'
        'Predicted'             -> '4: Predicted'
    """
    if not isinstance(pe_value, str):
        return pe_value

    clean = pe_value.strip()
    if not clean:
        return clean

    key = clean.lower()
    if key in PE_LEVELS:
        level = PE_LEVELS[key]
        return f"{level}: {clean}"
    # Already numeric? e.g. "3: Inferred from homology"
    if clean[0].isdigit():
        return clean
    return clean


def query_uniprot_chunk(accessions: List[str],
                        sleep_between: float = 0.5,
                        max_retries: int = 3) -> Dict[str, Dict[str, str]]:
    """
    Query UniProt REST API for a chunk of base accessions.

    Returns:
        dict[accession] -> {
            'organism_name': str or None,
            'reviewed': 'True'/'False' or None,
            'protein_existence': formatted str or None
        }
    """
    if not accessions:
        return {}

    # Build query: (accession:A0A023GZ41) OR (accession:A0A023H4N3) ...
    query = " OR ".join(f"(accession:{acc})" for acc in accessions)

    params = {
        "query": query,
        "fields": "accession,organism_name,reviewed,protein_existence",
        "format": "tsv",
        "size": len(accessions),
    }

    for attempt in range(1, max_retries + 1):
        try:
            resp = requests.get(UNIPROT_SEARCH_URL, params=params, timeout=60)
            if resp.status_code == 200:
                break
            else:
                sys.stderr.write(
                    f"[WARN] UniProt request failed (status {resp.status_code}), "
                    f"attempt {attempt}/{max_retries}\n"
                )
        except requests.RequestException as e:
            sys.stderr.write(
                f"[WARN] UniProt request exception on attempt {attempt}/{max_retries}: {e}\n"
            )
        time.sleep(sleep_between)
    else:
        sys.stderr.write(
            "[ERROR] UniProt request failed after maximum retries; "
            "this chunk will be skipped.\n"
        )
        return {}

    text = resp.text.strip()
    if not text:
        return {}

    lines = text.splitlines()
    if len(lines) <= 1:
        return {}

    header = lines[0].split("\t")
    norm_header = [normalise_header_name(h) for h in header]
    col_idx = {norm_header[i]: i for i in range(len(norm_header))}

    out: Dict[str, Dict[str, str]] = {}

    for line in lines[1:]:
        if not line.strip():
            continue
        cols = line.split("\t")

        # Build a row dict that tolerates missing trailing columns
        row: Dict[str, str] = {}
        for i, h in enumerate(norm_header):
            v = cols[i].strip() if i < len(cols) else ""
            row[h] = v

        acc = row.get("accession", "").strip()
        if not acc:
            continue

        organism_name = row.get("organism_name") or None

        # Reviewed: UniProt usually returns 'reviewed'/'unreviewed' or boolean-like values
        raw_rev = (row.get("reviewed") or "").strip().lower()
        if raw_rev in ("reviewed", "true"):
            reviewed_flag = "True"
        elif raw_rev in ("unreviewed", "false"):
            reviewed_flag = "False"
        elif raw_rev:
            reviewed_flag = raw_rev
        else:
            reviewed_flag = None

        raw_pe = (row.get("protein_existence") or "").strip()
        protein_existence = format_protein_existence(raw_pe) if raw_pe else None

        out[acc] = {
            "organism_name": organism_name,
            "reviewed": reviewed_flag,
            "protein_existence": protein_existence,
        }

    return out


def fetch_uniprot_metadata(base_accessions: Iterable[str]) -> Dict[str, Dict[str, str]]:
    """
    Fetch UniProt metadata for all base accessions, in chunks.

    Returns:
        dict[base_accession] -> {
            'organism_name': ...,
            'reviewed': ...,
            'protein_existence': ...
        }
    """
    unique_accs = sorted(set(a for a in base_accessions if isinstance(a, str) and a))
    sys.stderr.write(
        f"[INFO] Fetching UniProt metadata for {len(unique_accs)} base accessions...\n"
    )

    meta: Dict[str, Dict[str, str]] = {}
    for i, chunk in enumerate(chunked(unique_accs, 100), start=1):
        sys.stderr.write(
            f"[INFO]  Querying UniProt for chunk {i} ({len(chunk)} accessions)...\n"
        )
        chunk_meta = query_uniprot_chunk(chunk)
        meta.update(chunk_meta)
        # be gentle to the API
        time.sleep(0.2)

    sys.stderr.write(
        f"[INFO] Received metadata for {len(meta)} / {len(unique_accs)} accessions.\n"
    )
    missing = len(unique_accs) - len(meta)
    if missing > 0:
        sys.stderr.write(
            f"[WARN] No UniProt metadata for {missing} accessions "
            "(accession not found or request failure).\n"
        )
    return meta


def chain_index_to_letter(idx: int) -> str:
    """
    Convert an integer chain index (1,2,3,...) to a chain letter like A,B,C,...

    1 -> A
    2 -> B
    ...
    26 -> Z
    27 -> AA
    etc. (simple base-26 A–Z scheme)
    """
    if idx < 1:
        return "A"
    letters = []
    while idx > 0:
        idx, rem = divmod(idx - 1, 26)
        letters.append(chr(ord("A") + rem))
    return "".join(reversed(letters))


def pdb_filename_to_uniprot_accession(pdb_name: str) -> str:
    """
    Convert a BFVD PDB filename like:
        A0A2P1GMZ4_1_unrelaxed_rank_001_alphafold2_ptm_model_2_seed_000.pdb
    into your shard accession like:
        A0A2P1GMZ4_A
    """
    base = os.path.basename(pdb_name)
    base = base.split(".")[0]  # remove .pdb
    parts = base.split("_")
    if not parts:
        return pdb_name.strip()

    base_acc = parts[0]
    chain_idx = 1
    if len(parts) > 1:
        try:
            chain_idx = int(parts[1])
        except ValueError:
            chain_idx = 1

    chain_letter = chain_index_to_letter(chain_idx)
    return f"{base_acc}_{chain_letter}"


def load_bfvd_taxid(path: str) -> pd.DataFrame:
    """
    Load bfvd_taxid.tsv in a robust way.

    Supported formats:
    1) With headers:
         uniprot_accession <tab> taxid
    2) Without headers, but first col is PDB filename, second is taxid:
         A0A2P1GMZ4_1_unrelaxed_rank_001_alphafold2_ptm_model_2_seed_000.pdb  2116385
    3) Without headers, but first col is already uniprot_accession, second is taxid.

    Returns DataFrame with columns:
        uniprot_accession, taxid
    """
    df = pd.read_csv(path, sep="\t", dtype=str)
    df_cols_lower = [c.strip().lower() for c in df.columns]

    if "uniprot_accession" in df_cols_lower and "taxid" in df_cols_lower:
        # Already in the right form
        col_map = {}
        for c in df.columns:
            lc = c.strip().lower()
            if lc == "uniprot_accession":
                col_map[c] = "uniprot_accession"
            elif lc == "taxid":
                col_map[c] = "taxid"
        df = df.rename(columns=col_map)
        return df[["uniprot_accession", "taxid"]]

    # If there is no proper header, re-read without header
    if df.shape[1] == 1:
        df = pd.read_csv(path, sep="\t", header=None, dtype=str)

    if df.shape[1] < 2:
        raise SystemExit(
            f"[ERROR] bfvd_taxid file {path} must have at least two columns "
            "(pdb_id/uniprot_accession and taxid)."
        )

    # Interpret first column as PDB filename or uniprot_accession, second as taxid
    col0 = df.iloc[:, 0].astype(str)
    col1 = df.iloc[:, 1].astype(str)

    first_val = col0.iloc[0]
    if isinstance(first_val, str) and first_val.strip().endswith(".pdb"):
        sys.stderr.write("[INFO] bfvd_taxid: detected PDB filenames in first column, converting to uniprot_accession...\n")
        uniprot_acc = col0.apply(pdb_filename_to_uniprot_accession)
    else:
        sys.stderr.write("[INFO] bfvd_taxid: assuming first column is already uniprot_accession.\n")
        uniprot_acc = col0

    taxid = col1
    out_df = pd.DataFrame({"uniprot_accession": uniprot_acc, "taxid": taxid})
    return out_df


def parse_args() -> argparse.Namespace:
    p = argparse.ArgumentParser(
        description=(
            "Join a BFVD TMbed CSV with bfvd_taxid.tsv and enrich with UniProt "
            "taxonomy metadata (organism_name, reviewed, protein_existence)."
        )
    )
    p.add_argument(
        "--input",
        required=True,
        help="Input CSV with TMbed predictions (uniprot_accession,...,sequence,labels).",
    )
    p.add_argument(
        "--bfvd-taxid",
        required=True,
        help="TSV file with mapping: either 'uniprot_accession taxid' or 'pdb_filename taxid'.",
    )
    p.add_argument(
        "--output",
        required=True,
        help="Output CSV with taxonomy-enriched data.",
    )
    return p.parse_args()


def main() -> None:
    args = parse_args()

    # -----------------------------
    # 1) Load TMbed prediction CSV
    # -----------------------------
    sys.stderr.write(f"[INFO] Loading TMbed CSV: {args.input}\n")
    df = pd.read_csv(args.input)

    required_cols = [
        "uniprot_accession",
        "length",
        "tm_class",
        "has_signal_peptide",
        "num_TMH",
        "num_TMB",
        "segments",
        "sequence",
        "labels",
    ]
    missing_cols = [c for c in required_cols if c not in df.columns]
    if missing_cols:
        raise SystemExit(
            f"[ERROR] Input CSV is missing required columns: {missing_cols}"
        )

    # -----------------------------
    # 2) Load bfvd_taxid TSV (robust)
    # -----------------------------
    sys.stderr.write(f"[INFO] Loading bfvd_taxid TSV: {args.bfvd_taxid}\n")
    tax_df = load_bfvd_taxid(args.bfvd_taxid)

    # -----------------------------
    # 3) Join TMbed CSV with taxid
    # -----------------------------
    sys.stderr.write("[INFO] Joining TMbed CSV with bfvd_taxid on 'uniprot_accession'...\n")
    df = df.merge(
        tax_df[["uniprot_accession", "taxid"]],
        on="uniprot_accession",
        how="left",
        validate="m:1",
    )

    missing_taxid = df["taxid"].isna().sum()
    if missing_taxid > 0:
        sys.stderr.write(
            f"[WARN] {missing_taxid} rows have no 'taxid' after joining bfvd_taxid.\n"
        )

    # -----------------------------
    # 4) Compute base accession for UniProt queries
    #    (strip AFDB chain suffix: A0A023GZ41_A -> A0A023GZ41)
    # -----------------------------
    df["base_accession"] = df["uniprot_accession"].astype(str).str.split("_").str[0]

    # -----------------------------
    # 5) Fetch UniProt metadata
    #    -> only for rows that actually have a taxid (to limit queries)
    # -----------------------------
    mask_has_taxid = df["taxid"].notna()
    accs_for_query = df.loc[mask_has_taxid, "base_accession"]
    sys.stderr.write(
        f"[INFO] Restricting UniProt queries to {accs_for_query.nunique()} base accessions with taxid.\n"
    )
    meta = fetch_uniprot_metadata(accs_for_query)

    # Map metadata back to each row
    org_names: List[str] = []
    reviewed_flags: List[str] = []
    pe_values: List[str] = []

    for base_acc, has_tax in zip(df["base_accession"], df["taxid"].notna()):
        if has_tax:
            info = meta.get(base_acc, {})
        else:
            info = {}
        org_names.append(info.get("organism_name"))
        reviewed_flags.append(info.get("reviewed"))
        pe_values.append(info.get("protein_existence"))

    df["organism_name"] = org_names
    df["reviewed"] = reviewed_flags
    df["protein_existence"] = pe_values

    # -----------------------------
    # 6) Set domain column
    # -----------------------------
    df["domain"] = "Other"

    # -----------------------------
    # 7) Select and write output columns
    # -----------------------------
    out_cols = [
        "uniprot_accession",
        "length",
        "tm_class",
        "has_signal_peptide",
        "num_TMH",
        "num_TMB",
        "domain",
        "taxid",
        "organism_name",
        "reviewed",
        "protein_existence",
    ]

    missing_out = [c for c in out_cols if c not in df.columns]
    if missing_out:
        raise SystemExit(
            f"[ERROR] Internal error: missing expected output columns {missing_out}"
        )

    sys.stderr.write(f"[INFO] Writing enriched CSV to: {args.output}\n")
    df[out_cols].to_csv(args.output, index=False)
    sys.stderr.write("[INFO] Done.\n")


if __name__ == "__main__":
    main()
```
