## Supplementary material for "*biocentral*: embedding-based protein predictions": enrich_envelope_status.py

```
#!/usr/bin/env python3
# -*- coding: utf-8 -*-

"""
Hardcoded enrichment: add envelope_status using (priority):
  1) NCBI Taxonomy via ete3.NCBITaxa (family/order from taxid)
  2) Order-level fallback rules (uniform envelope behavior)
  3) Name-based heuristics for common cases (phage, HIV-1, Hantaviridae,
     Bromoviridae, etc.)
Also merges a curated family->envelope map if present.

CLI usage (example):

  py "enrich_envelope_status.py" ^
     --input  "bfvd_with_taxonomy.csv" ^
     --output "bfvd_with_envelope.csv" ^
     --family-map "family_envelope.csv"

Plots and statistics are generated by default next to the output file.
Use --no-plots to skip them or --plots-prefix to change filenames.
"""

from pathlib import Path
import sys
import argparse

import numpy as np
import pandas as pd
import matplotlib.pyplot as plt

# ------------------ DEFAULT PATHS (used if CLI args omitted) ------------------
BASE = Path(r"C:\Users\smaou\OneDrive\Bureau\BA\BFVD_fasta_shards\merged")
DEFAULT_INPUT_CSV = BASE / "bfvd_with_taxonomy.csv"
DEFAULT_OUTPUT_CSV = BASE / "bfvd_with_envelope.csv"
DEFAULT_FAMILY_MAP = BASE / "family_envelope.csv"  # optional curated family map
# -----------------------------------------------------------------------------


# Try to import ete3 for robust TaxID->lineage resolution
HAS_ETE = False
try:
    from ete3 import NCBITaxa  # type: ignore
    HAS_ETE = True
except Exception as e:
    print("[WARN] ete3/NCBITaxa import failed; will rely on heuristics only.")
    print("       Detailed error:", repr(e))
    HAS_ETE = False


def norm(x):
    if pd.isna(x):
        return None
    return " ".join(str(x).strip().lower().split())


# ---- Order-level defaults (override with family if available) ----
ORDER_ENVELOPE_DEFAULT = {
    # Mostly enveloped negative-sense RNA
    "mononegavirales": "Enveloped",
    "bunyavirales": "Enveloped",
    "articulavirales": "Enveloped",
    "nidovirales": "Enveloped",
    "herpesvirales": "Enveloped",
    "orfavirales": "Enveloped",
    "mulpavirales": "Enveloped",

    # Mostly non-enveloped positive/other
    "picornavirales": "Non-enveloped",
    "dividovirales": "Non-enveloped",
    "martellivirales": "Non-enveloped",
    "sobelivirales": "Non-enveloped",
    "timlovirales": "Non-enveloped",
    "nodamuvirales": "Non-enveloped",
    "stellavirales": "Non-enveloped",
    "patatavirales": "Non-enveloped",

    # dsRNA
    "reovirales": "Non-enveloped",
}

# ---- Family-level patches / expansions ----
FAMILY_ENVELOPE_FALLBACK = {
    "retroviridae": "Enveloped",
    "rhabdoviridae": "Enveloped",
    "paramyxoviridae": "Enveloped",
    "filoviridae": "Enveloped",
    "orthomyxoviridae": "Enveloped",
    "coronaviridae": "Enveloped",
    "arteriviridae": "Enveloped",
    "herpesviridae": "Enveloped",
    "poxviridae": "Enveloped",
    "hepadnaviridae": "Enveloped",
    "arenaviridae": "Enveloped",
    "hantaviridae": "Enveloped",
    "peribunyaviridae": "Enveloped",
    "phenuiviridae": "Enveloped",
    "nairoviridae": "Enveloped",

    "adenoviridae": "Non-enveloped",
    "papillomaviridae": "Non-enveloped",
    "polyomaviridae": "Non-enveloped",
    "picornaviridae": "Non-enveloped",
    "astroviridae": "Non-enveloped",
    "caliciviridae": "Non-enveloped",
    "parvoviridae": "Non-enveloped",
    "reoviridae": "Non-enveloped",
    "birnaviridae": "Non-enveloped",

    # Plant examples:
    "bromoviridae": "Non-enveloped",
    "virgaviridae": "Non-enveloped",
    "secoiviridae": "Non-enveloped",
    "tombusviridae": "Non-enveloped",

    # Archaeal spindle/lemon
    "fuselloviridae": "Enveloped",
    "lipothrixviridae": "Enveloped",
    "rudiviridae": "Non-enveloped",
}


# ---- Name heuristics ----
def heuristic_family_from_name(name_norm: str):
    if not name_norm:
        return None, None
    # HIV-1
    if "human immunodeficiency virus" in name_norm or name_norm.startswith("hiv"):
        return "Retroviridae", "name_heuristic"
    # orthohantavirus
    if "orthohantavirus" in name_norm or "hantavirus" in name_norm:
        return "Hantaviridae", "name_heuristic"
    # alfalfa mosaic virus, tomato aspermy virus -> Bromoviridae
    if "alfalfa mosaic virus" in name_norm or "tomato aspermy virus" in name_norm:
        return "Bromoviridae", "name_heuristic"
    # granulosis virus
    if "granulosis virus" in name_norm:
        return "Baculoviridae", "name_heuristic"
    # limnipivirus
    if "limnipivirus" in name_norm:
        return "Rhabdoviridae", "name_heuristic"
    # generic picornavirus
    if "picornavirus" in name_norm:
        return "Picornaviridae", "name_heuristic"
    # generic phage
    if " phage" in name_norm or name_norm.startswith("phage ") or name_norm.endswith(" phage"):
        return None, "phage_heuristic"
    return None, None


# -----------------------------------------------------------------------------


def infer_tm_columns(df: pd.DataFrame) -> pd.DataFrame:
    """
    Add:
      - has_tm (bool)
      - tm_class_simple in {TMH-only, TMB-only, Both, None}
    if possible. If necessary columns are missing, this is a no-op.
    """
    num_tmh = None
    num_tmb = None

    if "num_TMH" in df.columns:
        num_tmh = pd.to_numeric(df["num_TMH"], errors="coerce").fillna(0).astype(int)
    if "num_TMB" in df.columns:
        num_tmb = pd.to_numeric(df["num_TMB"], errors="coerce").fillna(0).astype(int)

    # Option 1: explicit counts
    if num_tmh is not None or num_tmb is not None:
        if num_tmh is None:
            num_tmh = pd.Series(0, index=df.index)
        if num_tmb is None:
            num_tmb = pd.Series(0, index=df.index)

        df["has_tm"] = (num_tmh > 0) | (num_tmb > 0)

        def _class_from_counts(tmh, tmb):
            if tmh > 0 and tmb == 0:
                return "TMH-only"
            if tmb > 0 and tmh == 0:
                return "TMB-only"
            if tmb > 0 and tmh > 0:
                return "Both"
            return "None"

        df["tm_class_simple"] = [
            _class_from_counts(tmh, tmb) for tmh, tmb in zip(num_tmh, num_tmb)
        ]
        return df

    # Option 2: tm_class string
    if "tm_class" in df.columns:
        raw = df["tm_class"].astype(str).str.strip().str.lower()

        def _class_from_string(s: str) -> str:
            if s in {"tmh", "tmh-only", "alpha", "alpha_only"}:
                return "TMH-only"
            if s in {"tmb", "tmb-only", "beta", "beta_only"}:
                return "TMB-only"
            if s in {"both", "tmh+tmb", "alpha+beta"}:
                return "Both"
            if s in {"none", "other", "no_tm", "nan", ""}:
                return "None"
            return "None"

        df["tm_class_simple"] = raw.map(_class_from_string)
        df["has_tm"] = df["tm_class_simple"].isin(["TMH-only", "TMB-only", "Both"])
        return df

    print("[INFO] Could not infer TM columns (no num_TMH/num_TMB/tm_class).")
    return df


def make_plots(df: pd.DataFrame, plot_prefix: Path):
    """
    Create plots:

    1) Envelope labels (overall)  -> <prefix>_counts_envelope_overall.png
    2) Envelope status composition pie -> <prefix>_envelope_pie.png
    3) Fraction of proteins with ≥1 TM by envelope group ->
         <prefix>_TM_fraction_envelope.png
    4) TM fraction by envelope group (all) ->
         <prefix>_tm_fraction_all.png
    5) TM class composition by envelope group ->
         <prefix>_TMclass_by_envelope.png
    """
    if "envelope_status" not in df.columns:
        print("[WARN] 'envelope_status' column missing; skipping all plots.")
        return

    # Normalize envelope labels into a clean column
    env = df["envelope_status"].fillna("unknown").astype(str)
    env_norm = env.str.strip()
    df = df.copy()
    df["envelope_norm"] = env_norm

    # -------- 1) Barplot: counts per envelope_norm  --------
    counts = df["envelope_norm"].value_counts().sort_index()
    if counts.empty:
        print("[WARN] No envelope labels for plotting.")
        return

    plt.figure(figsize=(7, 5))
    counts.plot(kind="bar")
    plt.ylabel("Count")
    plt.xlabel("envelope_norm")
    plt.title("Envelope labels (overall)")
    plt.tight_layout()
    plt.savefig(
        plot_prefix.with_name(plot_prefix.name + "_counts_envelope_overall.png"),
        dpi=150,
    )
    plt.close()

    # -------- 2) Pie chart: composition --------
    plt.figure(figsize=(8, 6))
    counts.plot(kind="pie", autopct="%0.1f%%")
    plt.ylabel("")
    plt.title("Envelope status composition")
    plt.tight_layout()
    plt.savefig(
        plot_prefix.with_name(plot_prefix.name + "_envelope_pie.png"),
        dpi=150,
    )
    plt.close()

    # -------- TM-based plots need has_tm & tm_class_simple --------
    if "has_tm" not in df.columns or "tm_class_simple" not in df.columns:
        print("[INFO] No TM information (has_tm/tm_class_simple); skipping TM plots.")
        return

    # 3) Fraction of proteins with ≥1 TM by envelope group
    frac = df.groupby("envelope_norm")["has_tm"].mean().sort_index()
    plt.figure(figsize=(7, 5))
    bars = plt.bar(frac.index.astype(str), frac.values)
    plt.ylim(0, 1.0)
    plt.ylabel("Fraction")
    plt.xlabel("envelope_status")
    plt.title("Fraction of proteins with ≥1 TM region")
    for b, v in zip(bars, frac.values):
        plt.text(
            b.get_x() + b.get_width() / 2,
            v + 0.01,
            f"{v * 100:.1f}%",
            ha="center",
            va="bottom",
        )
    plt.tight_layout()
    plt.savefig(
        plot_prefix.with_name(plot_prefix.name + "_TM_fraction_envelope.png"),
        dpi=150,
    )
    plt.close()

    # 4) Same info, separate figure
    plt.figure(figsize=(7, 5))
    bars = plt.bar(frac.index.astype(str), frac.values)
    plt.ylim(0, 1.0)
    plt.ylabel("Fraction with ≥1 TM")
    plt.xlabel("envelope_status")
    plt.title("TM fraction by envelope group (all)")
    for b, v in zip(bars, frac.values):
        plt.text(
            b.get_x() + b.get_width() / 2,
            v + 0.01,
            f"{v:.2f}",
            ha="center",
            va="bottom",
        )
    plt.tight_layout()
    plt.savefig(
        plot_prefix.with_name(plot_prefix.name + "_tm_fraction_all.png"),
        dpi=150,
    )
    plt.close()

    # 5) Stacked bar: TM class composition by envelope group
    cats = ["TMH-only", "TMB-only", "Both", "None"]

    tmp = df.copy()
    tmp["tm_class_simple"] = tmp["tm_class_simple"].fillna("None")

    # Pivot table: counts per (envelope_norm, tm_class_simple)
    comp = tmp.pivot_table(
        index="envelope_norm",
        columns="tm_class_simple",
        values="__name__",   # any column, we just count
        aggfunc="size",
        fill_value=0,
    )

    # Normalize rows to fractions
    comp = comp.div(comp.sum(axis=1), axis=0)

    # Ensure all categories exist as columns
    for c in cats:
        if c not in comp.columns:
            comp[c] = 0.0
    comp = comp[cats].sort_index()

    x_labels = comp.index.astype(str)
    x = np.arange(len(x_labels))

    plt.figure(figsize=(7, 5))
    bottom = np.zeros(len(comp))
    for cls in cats:
        vals = comp[cls].values
        plt.bar(x, vals, bottom=bottom, label=cls)
        bottom = bottom + vals

    plt.xticks(x, x_labels, rotation=0)
    plt.ylabel("Fraction")
    plt.xlabel("envelope_status")
    plt.title("TM class composition by envelope status")
    plt.legend()
    plt.tight_layout()
    plt.savefig(
        plot_prefix.with_name(plot_prefix.name + "_TMclass_by_envelope.png"),
        dpi=150,
    )
    plt.close()

    # Console summary
    print("[stats] Envelope label counts:")
    print(counts)
    print()
    print("[stats] Fraction with ≥1 TM by envelope_norm:")
    print(frac.round(3))


# -----------------------------------------------------------------------------


def make_tm_summary_table(df: pd.DataFrame, out_csv: Path):
    """
    Build a summary table like the paper's Table 2:

        Envelope Status | Nprot with TM predictions | Nprot without TM predictions
                        | Total | Percentage TM

    Uses:
      - df['envelope_status']  (values 'Enveloped', 'Non-enveloped', 'unknown', ...)
      - df['has_tm']           (True/False), provided by infer_tm_columns().
    """
    if "envelope_status" not in df.columns:
        print("[WARN] Cannot build TM summary table: 'envelope_status' missing.")
        return
    if "has_tm" not in df.columns:
        print("[WARN] Cannot build TM summary table: 'has_tm' missing.")
        return

    env = df["envelope_status"].fillna("unknown").astype(str).str.strip().str.lower()
    has_tm = df["has_tm"].fillna(False).astype(bool)

    # Map internal values to table row labels
    label_map = {
        "enveloped": "Envelope",
        "non-enveloped": "Non-Envelope",
        "non envelope": "Non-Envelope",
        "unknown": "Unknown",
    }

    # Ensure we only use the three main groups, in this order
    groups = ["enveloped", "non-enveloped", "unknown"]

    rows = []
    for g in groups:
        mask = env == g
        total = int(mask.sum())
        if total == 0:
            n_with = 0
            n_without = 0
            pct = 0.0
        else:
            n_with = int(has_tm[mask].sum())
            n_without = int(total - n_with)
            pct = 100.0 * n_with / total

        display_label = label_map.get(g, g.title())
        rows.append(
            {
                "Envelope Status": display_label,
                "Nprot with TM predictions": n_with,
                "Nprot without TM predictions": n_without,
                "Total": total,
                "Percentage TM": f"{pct:.1f}%",
            }
        )

    table_df = pd.DataFrame(rows)

    print("\n[TM summary table]")
    print(table_df.to_string(index=False))

    table_df.to_csv(out_csv, index=False)
    print(f"[TM summary] Wrote table to: {out_csv}")


# -----------------------------------------------------------------------------


def run_enrichment(
    input_csv: Path,
    output_csv: Path,
    family_map_path: Path | None = None,
) -> pd.DataFrame:
    # --- load input ---
    if not input_csv.exists():
        sys.exit(f"Input not found: {input_csv}")
    print(f"[INFO] Loading input CSV: {input_csv}")
    df = pd.read_csv(input_csv, dtype=str, low_memory=False)

    org_col = (
        "organism_name"
        if "organism_name" in df.columns
        else ("name" if "name" in df.columns else None)
    )
    if not org_col:
        sys.exit("Input needs 'organism_name' (or 'name').")

    df["__name__"] = df[org_col].apply(norm)
    has_taxid = "taxid" in df.columns

    # --- load curated family map if present ---
    fam_map: dict[str, str] = {}
    if family_map_path and family_map_path.exists():
        print(f"[INFO] Loading curated family map: {family_map_path}")
        env = pd.read_csv(family_map_path, dtype=str, low_memory=False)
        if "family" in env.columns:
            famc = "family"
            envc = (
                "envelope_status"
                if "envelope_status" in env.columns
                else ("enveloped" if "enveloped" in env.columns else None)
            )
            if envc:
                env["family_norm"] = env[famc].apply(norm)
                env = env.dropna(subset=["family_norm"]).drop_duplicates("family_norm")
                fam_map = dict(zip(env["family_norm"], env[envc]))

    # Add built-in fallback families:
    for fam, status in FAMILY_ENVELOPE_FALLBACK.items():
        fam_map.setdefault(fam.lower(), status)

    # --- NCBI taxonomy resolver (if available) ---
    ncbi = None
    if HAS_ETE:
        try:
            ncbi = NCBITaxa()
        except Exception as e:
            print("[WARN] NCBITaxa() initialization failed; falling back to heuristics.")
            print("       Detailed error:", repr(e))
            ncbi = None

    if ncbi is None:
        print("[WARN] ete3/NCBITaxa not available; relying on heuristics only.")

    family_resolved = []
    order_resolved = []
    envelope_status = []
    source_used = []

    print("[INFO] Starting envelope annotation ...")
    for _, row in df.iterrows():
        fam = None
        ordv = None
        envs = None
        src = None

        # 1) Try taxid -> family/order via NCBI
        if ncbi is not None and has_taxid and pd.notna(row.get("taxid")):
            try:
                tid = int(str(row["taxid"]))
                lineage = ncbi.get_lineage(tid)
                ranks = ncbi.get_rank(lineage)
                names = ncbi.get_taxid_translator(lineage)
                for t in lineage:
                    if ranks.get(t) == "family" and not fam:
                        fam = names.get(t)
                    if ranks.get(t) == "order" and not ordv:
                        ordv = names.get(t)
                src = "ncbi_taxid"
            except Exception:
                pass

        # 2) If still no family, try name heuristics
        if not fam:
            guess, hsrc = heuristic_family_from_name(row["__name__"])
            if guess:
                fam = guess
            if hsrc:
                src = (src + "+" + hsrc) if src else hsrc

        # 3) Decide envelope:
        fam_norm = norm(fam) if fam else None
        ord_norm = norm(ordv) if ordv else None

        # 3a) family-level map wins if present
        if fam_norm and fam_norm in fam_map:
            envs = fam_map[fam_norm]
            src = (src + "+family_map") if src else "family_map"
        # 3b) else order-level default
        if not envs and ord_norm and ord_norm in ORDER_ENVELOPE_DEFAULT:
            envs = ORDER_ENVELOPE_DEFAULT[ord_norm]
            src = (src + "+order_default") if src else "order_default"
        # 3c) else phage heuristic
        if not envs and row["__name__"]:
            nm = row["__name__"]
            if " phage" in nm or nm.startswith("phage ") or nm.endswith(" phage"):
                envs = "Non-enveloped"
                src = (src + "+phage_default") if src else "phage_default"

        # 3d) final fallback
        if not envs:
            envs = "unknown"
            src = src or "unknown"

        family_resolved.append(fam)
        order_resolved.append(ordv)
        envelope_status.append(envs)
        source_used.append(src)

    df["family_resolved"] = family_resolved
    df["order_resolved"] = order_resolved
    df["envelope_status"] = envelope_status
    df["envelope_source"] = source_used

    # Reorder columns so new columns are at the end
    cols = list(df.columns)
    order_hint = ["family_resolved", "order_resolved", "envelope_status", "envelope_source"]
    cols = [c for c in cols if c not in order_hint] + order_hint

    print(f"[INFO] Writing enriched CSV to: {output_csv}")
    df[cols].to_csv(output_csv, index=False)

    # summary
    total = len(df)
    known = (df["envelope_status"] != "unknown").sum()
    print(f"[done] total rows: {total:,}")
    print(f"[done] with envelope label: {known:,} ({known / total:.1%})")
    print(df["envelope_status"].value_counts(dropna=False))

    return df[cols]


# -----------------------------------------------------------------------------


def parse_args(argv=None):
    parser = argparse.ArgumentParser(
        description="Enrich BFVD/TMbed CSV with envelope_status labels "
        "using NCBI taxonomy + heuristics, and generate plots."
    )
    parser.add_argument(
        "-i",
        "--input",
        type=str,
        default=str(DEFAULT_INPUT_CSV),
        help=f"Input CSV (default: {DEFAULT_INPUT_CSV})",
    )
    parser.add_argument(
        "-o",
        "--output",
        type=str,
        default=str(DEFAULT_OUTPUT_CSV),
        help=f"Output CSV with envelope annotations (default: {DEFAULT_OUTPUT_CSV})",
    )
    parser.add_argument(
        "-f",
        "--family-map",
        type=str,
        default=str(DEFAULT_FAMILY_MAP),
        help=f"Optional curated family->envelope CSV (default: {DEFAULT_FAMILY_MAP})",
    )
    parser.add_argument(
        "--no-plots",
        action="store_true",
        help="Disable statistics and plot generation.",
    )
    parser.add_argument(
        "--plots-prefix",
        type=str,
        default=None,
        help=(
            "Prefix (path + stem) for plot filenames. "
            "Default: output CSV path without extension."
        ),
    )
    return parser.parse_args(argv)


def main(argv=None):
    args = parse_args(argv)

    input_csv = Path(args.input)
    output_csv = Path(args.output)
    family_map_path = Path(args.family_map) if args.family_map else None

    df = run_enrichment(input_csv, output_csv, family_map_path)

    # Add TM helper columns for plotting & summary (if possible)
    df = infer_tm_columns(df)

    # TM summary table, like the paper's Table 2
    summary_csv = output_csv.with_name(output_csv.stem + "_tm_summary.csv")
    make_tm_summary_table(df, summary_csv)

    if not args.no_plots:
        if args.plots_prefix is not None:
            plot_prefix = Path(args.plots_prefix)
        else:
            plot_prefix = output_csv.with_suffix("")  # same directory, stem of output
        print(f"[INFO] Generating plots with prefix: {plot_prefix}")
        make_plots(df, plot_prefix)
    else:
        print("[INFO] Plot generation disabled (--no-plots).")


if __name__ == "__main__":
    main()
```
