## Supplementary material for "*biocentral*: embedding-based protein predictions": enrich_envelope_status_ncbi_eutils_cli.py

```
# enrich_envelope_status_ncbi_eutils_cli.py
"""
CSV-only CLI to label envelope status using NCBI Taxonomy (E-utilities)

Usage:
  py enrich_envelope_status_ncbi_eutils_cli.py --input INPUT.csv --output OUTPUT.csv \
     [--family-map FAMILY_MAP.csv] [--cache CACHE.csv] [--sample-n 2000] [--]

Notes:
- If --cache is omitted, a temporary cache is used and removed after the run.
- Envelope logic is unchanged: family_map -> order defaults -> 'phage' name heuristic -> unknown.
- Prints end-of-run stats: group % and TM ratios (overall, per group, and binary-only).
"""

from __future__ import annotations
from pathlib import Path
import argparse
import csv
import logging
import math
import sys
import tempfile
import time
import xml.etree.ElementTree as ET

import pandas as pd
import requests

# --------- E-utilities defaults (unchanged) ---------
EUTILS_URL   = "https://eutils.ncbi.nlm.nih.gov/entrez/eutils/efetch.fcgi"
CHUNK        = 100       # taxids per request
REQ_TIMEOUT  = 60        # seconds per request
PAUSE        = 0.35      # pause between requests
MAX_RETRIES  = 3         # per batch

# --------- Envelope rules (unchanged) ---------
ORDER_ENVELOPE_DEFAULT = {
    "mononegavirales": "Enveloped",
    "bunyavirales":    "Enveloped",
    "articulavirales": "Enveloped",
    "nidovirales":     "Enveloped",
    "herpesvirales":   "Enveloped",
    "orfavirales":     "Enveloped",
    "mulpavirales":    "Enveloped",
    "picornavirales":  "Non-enveloped",
    "dividovirales":   "Non-enveloped",
    "martellivirales": "Non-enveloped",
    "sobelivirales":   "Non-enveloped",
    "timlovirales":    "Non-enveloped",
    "nodamuvirales":   "Non-enveloped",
    "stellavirales":   "Non-enveloped",
    "patatavirales":   "Non-enveloped",
    "reovirales":      "Non-enveloped",
}

FAMILY_ENVELOPE_FALLBACK = {
    "retroviridae":"Enveloped","rhabdoviridae":"Enveloped","paramyxoviridae":"Enveloped","filoviridae":"Enveloped",
    "orthomyxoviridae":"Enveloped","coronaviridae":"Enveloped","arteriviridae":"Enveloped","herpesviridae":"Enveloped",
    "poxviridae":"Enveloped","hepadnaviridae":"Enveloped","arenaviridae":"Enveloped","hantaviridae":"Enveloped",
    "peribunyaviridae":"Enveloped","phenuiviridae":"Enveloped","nairoviridae":"Enveloped",
    "adenoviridae":"Non-enveloped","papillomaviridae":"Non-enveloped","polyomaviridae":"Non-enveloped",
    "picornaviridae":"Non-enveloped","astroviridae":"Non-enveloped","caliciviridae":"Non-enveloped",
    "parvoviridae":"Non-enveloped","reoviridae":"Non-enveloped","birnaviridae":"Non-enveloped",
    "bromoviridae":"Non-enveloped","virgaviridae":"Non-enveloped","secoiviridae":"Non-enveloped","tombusviridae":"Non-enveloped",
    "fuselloviridae":"Enveloped","lipothrixviridae":"Enveloped","rudiviridae":"Non-enveloped",
}

# ----------------- Helpers -----------------
def norm(x):
    if pd.isna(x): return None
    return " ".join(str(x).strip().lower().split())

def read_csv_any(path: Path) -> pd.DataFrame:
    if not path.exists():
        sys.exit(f"[ERR] Not found: {path}")
    try:
        return pd.read_csv(path, dtype=str, sep=None, engine="python", low_memory=False)
    except Exception:
        return pd.read_csv(path, dtype=str, low_memory=False)

def write_csv(df: pd.DataFrame, path: Path):
    path.parent.mkdir(parents=True, exist_ok=True
    )
    df.to_csv(path, index=False)

def load_family_map_csv(path: Path | None) -> dict[str, str]:
    fam_map = {}
    if not path:
        return fam_map
    if not path.exists():
        logging.info(f"[info] FAMILY_MAP not found → skip: {path}")
        return fam_map
    df = read_csv_any(path)
    cols = {c.lower(): c for c in df.columns}
    fam_col = cols.get("family") or cols.get("family_resolved") or cols.get("family_norm")
    env_col = (cols.get("envelope_status") or cols.get("enveloped") or
               cols.get("envelope") or cols.get("label"))
    if not fam_col or not env_col:
        logging.warning(f"[warn] FAMILY_MAP lacks expected columns; found: {list(df.columns)}")
        return fam_map
    df["family_norm"] = df[fam_col].apply(norm)
    df = df.dropna(subset=["family_norm"]).drop_duplicates("family_norm")
    fam_map = dict(zip(df["family_norm"], df[env_col]))
    return fam_map

# ---- Cache (CSV: taxid,family,order) ----
def load_cache(path: Path) -> dict[str, dict[str, str | None]]:
    cache: dict[str, dict[str, str | None]] = {}
    if not path.exists(): return cache
    with path.open("r", newline="", encoding="utf-8") as f:
        reader = csv.DictReader(f)
        for r in reader:
            tid = str(r.get("taxid","")).strip()
            if not tid: continue
            cache[tid] = {"family": r.get("family") or None, "order": r.get("order") or None}
    return cache

def save_cache(path: Path, cache: dict[str, dict[str, str | None]]):
    path.parent.mkdir(parents=True, exist_ok=True)
    with path.open("w", newline="", encoding="utf-8") as f:
        w = csv.writer(f)
        w.writerow(["taxid","family","order"])
        for tid, v in cache.items():
            w.writerow([tid, v.get("family") or "", v.get("order") or ""])

# ---- E-utilities fetch ----
def fetch_taxonomy_batch(session: requests.Session, taxids: list[str], email: str | None, tool: str | None) -> dict[str, dict[str, str | None]]:
    params = {"db":"taxonomy", "id": ",".join(taxids), "retmode":"xml"}
    if email: params["email"] = email
    if tool:  params["tool"] = tool
    r = session.get(EUTILS_URL, params=params, timeout=REQ_TIMEOUT)
    r.raise_for_status()
    out: dict[str, dict[str, str | None]] = {}
    root = ET.fromstring(r.text)
    for tx in root.findall("./Taxon"):
        tid_el = tx.find("TaxId")
        if tid_el is None: continue
        tid = tid_el.text.strip()
        fam_name, ord_name = None, None
        lin = tx.find("LineageEx")
        if lin is not None:
            for t in lin.findall("Taxon"):
                rank = (t.findtext("Rank") or "").strip().lower()
                name = (t.findtext("ScientificName") or "").strip()
                if rank == "family" and fam_name is None: fam_name = name
                elif rank == "order" and ord_name is None: ord_name = name
        out[tid] = {"family": fam_name, "order": ord_name}
    return out

def resolve_taxonomy_via_ncbi(taxids: list[str], cache_path: Path,
                              email: str | None, tool: str | None) -> dict[str, dict[str, str | None]]:
    cache = load_cache(cache_path)
    need = [tid for tid in taxids if tid and tid not in cache]
    logging.info(f"[ncbi] unique taxids: {len(taxids):,} | in cache: {len(taxids)-len(need):,} | to fetch: {len(need):,}")
    if not need:
        return cache

    sess = requests.Session()
    sess.headers.update({"User-Agent": f"{tool or 'bfvd-envelope-labeller'} (pandas requests)", "Accept-Encoding": "gzip, deflate"})

    total = len(need)
    batches = math.ceil(total/CHUNK)
    start = time.time()

    for bi in range(batches):
        chunk = need[bi*CHUNK:(bi+1)*CHUNK]
        last_err = None
        for attempt in range(1, MAX_RETRIES+1):
            try:
                got = fetch_taxonomy_batch(sess, chunk, email, tool)
                cache.update(got)
                break
            except Exception as e:
                last_err = e
                backoff = 2 ** (attempt - 1)
                logging.warning(f"[warn] batch {bi+1}/{batches} failed (try {attempt}/{MAX_RETRIES}): {e}. Backoff {backoff}s.")
                time.sleep(backoff)
        else:
            logging.error(f"[err] batch {bi+1}/{batches} final failure: {last_err}. Skipping this batch.")

        # persist incrementally
        save_cache(cache_path, cache)

        # progress
        done = min((bi+1)*CHUNK, total)
        elapsed = time.time() - start
        pct = 100.0 * done / total
        rate = done / elapsed if elapsed > 0 else 0.0
        eta = (total - done) / rate if rate > 0 else float("nan")
        logging.info(f"[ncbi] {done:,}/{total:,} ({pct:5.1f}%) | elapsed {elapsed:5.1f}s | ~{rate:6.1f} ids/s | ETA {eta:6.1f}s")

        time.sleep(PAUSE)

    return cache

# ---- TM helpers (added stats, no change to labeling logic) ----
def ensure_has_tm(df: pd.DataFrame) -> tuple[pd.DataFrame, bool]:
    """
    Create/overwrite has_TM_segment as 0/1 using num_TMH/num_TMB if available; fallback to tm_class.
    Returns (df, ok) where ok indicates whether TM info could be computed.
    """
    if "num_TMH" in df.columns and "num_TMB" in df.columns:
        df["num_TMH"] = pd.to_numeric(df["num_TMH"], errors="coerce").fillna(0).astype(int)
        df["num_TMB"] = pd.to_numeric(df["num_TMB"], errors="coerce").fillna(0).astype(int)
        df["has_TM_segment"] = ((df["num_TMH"] + df["num_TMB"]) > 0).astype(int)
        return df, True
    if "tm_class" in df.columns:
        t = df["tm_class"].astype(str).str.upper()
        df["has_TM_segment"] = t.isin(["TMH","TMB","TMH+TMB","TM"]).astype(int)
        return df, True
    logging.warning("[warn] No TM columns found. Expected ('num_TMH' & 'num_TMB') or 'tm_class'. Skipping TM stats.")
    return df, False

def normalize_envelope_values(s: pd.Series) -> pd.Series:
    def map_env(v):
        v = norm(v)
        if v in (None, "", "nan"): return "Unknown"
        if v in {"enveloped","yes"}: return "Enveloped"
        if v in {"non-enveloped","non enveloped","no","nonenveloped"}: return "Non-enveloped"
        if v in {"variable","yes or no","predicted yes"}: return "Variable"
        return v.capitalize() if v else "Unknown"
    return s.map(map_env)

# ----------------- Main -----------------
def main():
    ap = argparse.ArgumentParser(description="Label envelope status by resolving family/order from NCBI Taxonomy (CSV only).")
    ap.add_argument("--input",  required=True, help="Input CSV with columns including 'taxid' and 'organism_name' (or 'name').")
    ap.add_argument("--output", required=True, help="Output CSV path.")
    ap.add_argument("--family-map", default=None, help="Optional CSV mapping family->envelope label.")
    ap.add_argument("--cache", default=None, help="Optional cache CSV (taxid,family,order). If omitted, a temp cache is used and removed.")
    ap.add_argument("--sample-n", type=int, default=None, help="Optional cap on unique taxids to fetch (for quick test).")
    ap.add_argument("--email", default=None, help="Optional contact email for NCBI E-utilities.")
    ap.add_argument("--tool-name", default="bfvd-envelope-labeller", help="Tool name for NCBI requests header.")
    ap.add_argument("--log-level", default="INFO", choices=["DEBUG","INFO","WARNING","ERROR","CRITICAL"], help="Logging verbosity.")
    args = ap.parse_args()

    logging.basicConfig(level=getattr(logging, args.log_level), format="%(asctime)s %(levelname)s %(message)s")

    input_path   = Path(args.input)
    output_path  = Path(args.output)
    fam_map_path = Path(args.family_map) if args.family_map else None

    # Read input CSV
    df = read_csv_any(input_path)
    logging.info(f"[info] Input columns: {list(df.columns)}")

    # Sanity checks
    if "taxid" not in df.columns:
        sys.exit("[ERR] Input needs a 'taxid' column.")
    org_col = "organism_name" if "organism_name" in df.columns else ("name" if "name" in df.columns else None)
    if not org_col:
        sys.exit("[ERR] Input needs 'organism_name' (or 'name').")

    df["__name__"] = df[org_col].apply(norm)
    df["taxid"] = df["taxid"].astype(str).str.strip()

    # Unique taxids
    uniq_taxids = sorted(set(df["taxid"].dropna().tolist()))
    if args.sample_n:
        uniq_taxids = uniq_taxids[:int(args.sample_n)]
        logging.info(f"[debug] SAMPLE_N active → limiting to {len(uniq_taxids)} unique taxids for a quick run.")

    # Cache handling (temp if not provided)
    if args.cache:
        cache_path = Path(args.cache)
        cache_path.parent.mkdir(parents=True, exist_ok=True)
        temp_ctx = None
        logging.info(f"[cache] Using persistent cache: {cache_path}")
    else:
        temp_ctx = tempfile.TemporaryDirectory(prefix="ncbi_tax_cache_")
        cache_path = Path(temp_ctx.name) / "taxid_family_order_cache.csv"
        logging.info(f"[cache] Using temp cache: {cache_path} (will be removed after run)")

    try:
        # --- Resolve taxonomy ---
        taxmap = resolve_taxonomy_via_ncbi(uniq_taxids, cache_path, args.email, args.tool_name)

        fams, ords = [], []
        for _, row in df.iterrows():
            tid = str(row["taxid"]).strip()
            rec = taxmap.get(tid, {})
            fams.append(rec.get("family"))
            ords.append(rec.get("order"))
        df["family_resolved"] = fams
        df["order_resolved"]  = ords

        # --- Label envelope (unchanged logic) ---
        fam_map = load_family_map_csv(fam_map_path)
        for fam, status in FAMILY_ENVELOPE_FALLBACK.items():
            fam_map.setdefault(fam.lower(), status)

        env, src = [], []
        for _, row in df.iterrows():
            fam = norm(row.get("family_resolved"))
            ordv = norm(row.get("order_resolved"))
            e = None
            s = None
            if fam and fam in fam_map:
                e, s = fam_map[fam], "family_map"
            if not e and ordv and ordv in ORDER_ENVELOPE_DEFAULT:
                e, s = ORDER_ENVELOPE_DEFAULT[ordv], (s + "+order_default") if s else "order_default"
            nm = row["__name__"] or ""
            if not e and (nm.startswith("phage ") or nm.endswith(" phage") or " phage" in nm):
                e, s = "Non-enveloped", (s + "+phage_default") if s else "phage_default"
            if not e:
                e, s = "unknown", s or "unknown"
            env.append(e); src.append(s)

        df["envelope_status"] = env
        df["envelope_source"] = src

        # --- Summary (unchanged style) ---
        total = len(df)
        known = (df["envelope_status"] != "unknown").sum()
        logging.info(f"[done] total rows: {total:,}")
        logging.info(f"[done] with envelope label: {known:,} ({known/total:.1%})")
        logging.info("envelope_status counts:\n%s", df["envelope_status"].value_counts(dropna=False).to_string())
        logging.info("\n[source breakdown]:\n%s", df["envelope_source"].value_counts(dropna=False).head(10).to_string())

        # --- Extra end-of-run stats (TM ratios & group percentages) ---
        # Compute has_TM_segment if columns allow it
        df, tm_ok = ensure_has_tm(df)
        # Normalize envelope groups (handles 'Variable' if present in family map)
        env_group = normalize_envelope_values(df["envelope_status"])
        df["__env_group__"] = env_group

        # Group distribution (% of dataset)
        dist = (
            df["__env_group__"]
              .value_counts(dropna=False)
              .rename_axis("group")
              .reset_index(name="count")
        )
        dist["pct_of_dataset"] = (dist["count"] / total * 100).round(2)

        logging.info("\n=== Envelope group distribution ===")
        for _, r in dist.iterrows():
            logging.info(f"{r['group']:>13}:  {r['count']:>10}  ({r['pct_of_dataset']:>6.2f}%)")

        # Binary known share
        known_binary = df["__env_group__"].isin(["Enveloped","Non-enveloped"]).sum()
        logging.info(f"\n[known binary] {known_binary:,}/{total:,} ({known_binary/total:.1%})")

        if tm_ok:
            # Per-group TM percentages
            agg = (
                df.groupby("__env_group__", dropna=False)["has_TM_segment"]
                  .agg(total="count", tm_count="sum")
                  .reset_index()
            )
            agg["pct_with_TM"] = (agg["tm_count"] / agg["total"] * 100).round(2)

            logging.info("\n=== TM %% by Envelope Group ===")
            for _, r in agg.iterrows():
                logging.info(f"{r['__env_group__']:>13}:  TM {int(r['tm_count']):>8}/{int(r['total']):<8}  = {r['pct_with_TM']:>6.2f}%")

            # Binary-only quick comparison
            bin_df = df[df["__env_group__"].isin(["Enveloped","Non-enveloped"])].copy()
            if len(bin_df) >= 2:
                pe = bin_df.loc[bin_df["__env_group__"]=="Enveloped","has_TM_segment"].mean()
                pn = bin_df.loc[bin_df["__env_group__"]=="Non-enveloped","has_TM_segment"].mean()
                logging.info(f"\n[binary] Frac with TM | Enveloped: {pe:.3f}   Non-enveloped: {pn:.3f}   Δ={pe-pn:+.3f}")

        # --- Write output CSV ---
        hint = ["family_resolved","order_resolved","envelope_status","envelope_source"]
        cols = [c for c in df.columns if c not in hint] + hint
        write_csv(df[cols], output_path)
        logging.info(f"[write] {output_path}")

    finally:
        # Clean up temp cache if used
        if 'temp_ctx' in locals() and temp_ctx is not None:
            try:
                temp_ctx.cleanup()
                logging.info("[cache] Temp cache removed.")
            except Exception as e:
                logging.warning(f"[cache] Failed to remove temp cache: {e}")

if __name__ == "__main__":
    main()
```
