## Supplementary material for "*biocentral*: embedding-based protein predictions": filter_tmb_tmh.py

```
#!/usr/bin/env python3
"""
Filter proteins that have both TMH and TMB predictions (biologically unlikely),
enrich them with pLDDT from a local BFVD metadata TSV, and create plots.

Assumptions about BFVD metadata TSV (no header):
    col 0: UniProt ID      (e.g. A0A2P1GMZ4)
    col 1: model ID        (e.g. A0A2P1GMZ4_2)
    col 2: mean pLDDT      (float, 0–100)
    col 3: some extra metric (ignored)
    col 4: integer flag    (ignored)
    col 5: tag string      (ignored)

We aggregate by col 0 (UniProt ID) taking the mean of col 2 as pLDDT.

Main input CSV must have at least:
    - uniprot_accession
    - length
    - tm_class
    - num_TMH
    - num_TMB

Usage example
-------------

python filter_tmb_tmh.py \
    --input shards_merged.csv \
    --bfvd-meta "bfvd_metadata.tsv" \
    --output-prefix tm_conflicts_enriched \
    --out-format csv

Dependencies
------------
    pip install pandas numpy matplotlib
"""

import argparse
import os

import numpy as np
import pandas as pd
import matplotlib.pyplot as plt


# -------------------------------------------------------------------
# Helpers to load data
# -------------------------------------------------------------------

def load_main_table(path: str) -> pd.DataFrame:
    """Load main CSV with TM annotations."""
    df = pd.read_csv(path)

    required = ["uniprot_accession", "length", "tm_class", "num_TMH", "num_TMB"]
    missing = [c for c in required if c not in df.columns]
    if missing:
        raise ValueError(f"Input file '{path}' missing columns: {missing}")

    df["num_TMH"] = pd.to_numeric(df["num_TMH"], errors="coerce").fillna(0).astype(int)
    df["num_TMB"] = pd.to_numeric(df["num_TMB"], errors="coerce").fillna(0).astype(int)
    df["length"] = pd.to_numeric(df["length"], errors="coerce")

    return df


def filter_conflicts(df: pd.DataFrame) -> pd.DataFrame:
    """
    Select proteins that have BOTH TMHs and TMBs.

    Conditions:
        - num_TMH > 0 AND num_TMB > 0
      OR
        - tm_class (case-insensitive) == "BOTH"
    """
    tm_class = df["tm_class"].astype(str).str.upper()
    mask = (df["num_TMH"] > 0) & (df["num_TMB"] > 0)
    mask |= tm_class.eq("BOTH")
    return df.loc[mask].copy()


def strip_chain(acc: str) -> str:
    """
    Turn 'A0A2P1GMZ4_A' or 'A0A2P1GMZ4_3_A' into 'A0A2P1GMZ4'.

    We just take the first token before an underscore.
    Adjust if you know your pattern is different.
    """
    return acc.split("_")[0]


def load_bfvd_metadata(path: str) -> pd.DataFrame:
    """
    Load local BFVD metadata TSV (no header) with columns:
        0: UniProt ID
        1: model ID
        2: mean pLDDT
        3: extra
        4: extra
        5: extra/tag

    Returns a dataframe with:
        uniprot_id, plddt_mean_final
    where plddt_mean_final is the mean of col 2 per UniProt.
    """
    print(f"[INFO] Loading BFVD metadata from: {path}")
    meta = pd.read_csv(path, sep="\t", header=None)

    # Sanity check: we expect at least 3 columns
    if meta.shape[1] < 3:
        raise ValueError(
            f"BFVD metadata file '{path}' has {meta.shape[1]} columns, "
            f"expected at least 3 (UniProt, model, pLDDT)."
        )

    # col 0: uniprot, col 2: mean pLDDT
    meta = meta[[0, 2]].copy()
    meta.columns = ["uniprot_id", "plddt_model_mean"]

    # Aggregate across models for same UniProt: mean pLDDT
    agg = (
        meta.groupby("uniprot_id", as_index=False)["plddt_model_mean"]
        .mean()
        .rename(columns={"plddt_model_mean": "plddt_mean_final"})
    )

    print(f"[INFO] BFVD metadata: {len(agg)} unique UniProt IDs with pLDDT")
    return agg


def enrich_with_plddt(
    df_conflicts: pd.DataFrame,
    bfvd_meta: pd.DataFrame,
) -> pd.DataFrame:
    """
    Map conflicting UniProt accessions (with chain suffixes) to base UniProt,
    join BFVD metadata, and attach plddt_mean_final.
    """
    df = df_conflicts.copy()
    df["base_uniprot"] = df["uniprot_accession"].astype(str).apply(strip_chain)

    enriched = df.merge(
        bfvd_meta,
        left_on="base_uniprot",
        right_on="uniprot_id",
        how="left",
    )

    # Drop helper column from BFVD
    enriched.drop(columns=["uniprot_id"], inplace=True)

    found = enriched["plddt_mean_final"].notna().sum()
    total = len(enriched)
    print(f"[INFO] Found pLDDT for {found}/{total} conflicting proteins")

    return enriched


# -------------------------------------------------------------------
# Plotting
# -------------------------------------------------------------------

def plot_plddt_vs_length(df: pd.DataFrame, prefix: str):
    data = df[["length", "plddt_mean_final"]].dropna()
    if data.empty:
        print("[WARN] No data for pLDDT vs length scatter plot")
        return

    plt.figure(figsize=(7, 6))
    plt.scatter(data["length"], data["plddt_mean_final"], alpha=0.7)
    plt.xlabel("Sequence length (aa)")
    plt.ylabel("pLDDT mean")
    plt.title("pLDDT vs sequence length (TMH+TMB conflicts)")
    plt.grid(True, alpha=0.3)

    out = f"{prefix}_plddt_vs_length.png"
    plt.tight_layout()
    plt.savefig(out, dpi=300)
    plt.close()
    print(f"[INFO] Saved scatter plot to: {out}")


def plot_plddt_ecdf(df: pd.DataFrame, prefix: str):
    vals = df["plddt_mean_final"].dropna().values
    if len(vals) == 0:
        print("[WARN] No data for pLDDT ECDF plot")
        return

    vals = np.sort(vals)
    y = np.arange(1, len(vals) + 1) / len(vals)

    plt.figure(figsize=(7, 6))
    plt.step(vals, y, where="post")
    plt.xlabel("pLDDT mean")
    plt.ylabel("ECDF")
    plt.title("pLDDT distribution (TMH+TMB conflicts)")
    plt.grid(True, alpha=0.3)

    out = f"{prefix}_plddt_ecdf.png"
    plt.tight_layout()
    plt.savefig(out, dpi=300)
    plt.close()
    print(f"[INFO] Saved ECDF plot to: {out}")


# -------------------------------------------------------------------
# CLI
# -------------------------------------------------------------------

def parse_args():
    p = argparse.ArgumentParser(
        description=(
            "Filter TMH+TMB conflicts, enrich with pLDDT from local BFVD "
            "metadata (no downloads), and plot summary."
        )
    )
    p.add_argument(
        "--input", "-i", required=True,
        help="Input CSV with TM annotations (e.g. shards_merged.csv)"
    )
    p.add_argument(
        "--bfvd-meta", required=True,
        help="Local BFVD metadata TSV (e.g. 'bfvd_metadata (1).tsv')"
    )
    p.add_argument(
        "--output-prefix", "-o", default=None,
        help="Prefix for output files (default: derived from input filename)"
    )
    p.add_argument(
        "--out-format", choices=["csv", "jsonl"], default="csv",
        help="Output format for enriched table (default: csv)"
    )
    return p.parse_args()


def main():
    args = parse_args()

    input_path = args.input
    if args.output_prefix is None:
        base = os.path.splitext(os.path.basename(input_path))[0]
        output_prefix = base + "_tm_conflicts_enriched"
    else:
        output_prefix = args.output_prefix

    print(f"[INFO] Loading main CSV from: {input_path}")
    df = load_main_table(input_path)

    print("[INFO] Filtering TMH+TMB conflicts...")
    conflicts = filter_conflicts(df)
    print(f"[INFO] Found {len(conflicts)} conflicting proteins")

    if conflicts.empty:
        print("[INFO] No conflicts found, nothing to do.")
        return

    print(f"[INFO] Loading BFVD metadata from: {args.bfvd_meta}")
    bfvd_meta = load_bfvd_metadata(args.bfvd_meta)

    print("[INFO] Enriching conflicts with pLDDT from BFVD metadata...")
    enriched = enrich_with_plddt(conflicts, bfvd_meta)

    # Save enriched table
    if args.out_format == "csv":
        out_table = f"{output_prefix}.csv"
        enriched.to_csv(out_table, index=False)
    else:
        out_table = f"{output_prefix}.jsonl"
        enriched.to_json(out_table, orient="records", lines=True)
    print(f"[INFO] Written enriched table to: {out_table}")

    # Plots
    plot_plddt_vs_length(enriched, output_prefix)
    plot_plddt_ecdf(enriched, output_prefix)

    print("[INFO] Done.")


if __name__ == "__main__":
    main()
```
