## Supplementary material for "*biocentral*: embedding-based protein predictions": pdb_to_tmbed_fasta.py

```
#!/usr/bin/env python3
"""
BFVD PDB → FASTA extraction script

This script scans a directory of PDB files and reconstructs amino-acid sequences
for every chain. It extracts sequences either from SEQRES records or, if
missing, from ATOM records (CA backbone atoms). All sequences are converted from
3-letter to 1-letter codes, cleaned to the canonical 20 amino acids, wrapped at
60 characters, and written out into FASTA shards of a user-defined size
(default: 50,000 sequences per shard).

IMPORTANT — Why the output does NOT match the total BFVD chain count:

1) Chains shorter than 30 amino acids (after conversion + cleaning) are skipped.
   Many BFVD entries are tiny fragments, antibody regions, tags, or incomplete
   constructs. These are deliberately filtered out to keep only meaningful
   protein sequences.

2) Non-standard or modified residues (e.g., UNK, MSE, XAA) are removed during
   cleanup. If a chain consists mostly of such residues, it becomes very short
   or empty and is excluded by the length filter.

3) PDB files lacking SEQRES + CA-ATOM information cannot be reconstructed and
   therefore produce no sequence.

4) Any PDB that cannot be parsed due to format issues is skipped (logged as
   [skip]). This prevents the script from crashing when encountering malformed
   files.

Because of these filters, the number of sequences produced (≈345k) is lower than
the raw number of BFVD chains (351,242). This is expected and represents a
quality-filtered subset of the dataset, not a 1:1 reconstruction of all entries.

Output:
  - FASTA shards named bfvd_shard_00000.fasta, bfvd_shard_00001.fasta, ...
  - Each shard contains ≤ SHARD_SIZE sequences.

Usage:
  python extract_bfvd_pdb_sequences.py <input_dir> <output_dir> [SHARD_SIZE]

Example:
  python extract_bfvd_pdb_sequences.py ./bfvd_pdb/ ./out_fasta/ 50000
"""
import sys, re, os
from pathlib import Path

IN_DIR  = Path(sys.argv[1])
OUT_DIR = Path(sys.argv[2])
SHARD_SIZE = int(sys.argv[3]) if len(sys.argv) > 3 else 50000

OUT_DIR.mkdir(parents=True, exist_ok=True)

AA3 = {
    "ALA":"A","ARG":"R","ASN":"N","ASP":"D","CYS":"C","GLN":"Q","GLU":"E","GLY":"G",
    "HIS":"H","ILE":"I","LEU":"L","LYS":"K","MET":"M","PHE":"F","PRO":"P","SER":"S",
    "THR":"T","TRP":"W","TYR":"Y","VAL":"V",
    "MSE":"M","SEC":"C","PYL":"K","ASX":"N","GLX":"Q","XLE":"L","XAA":"", "UNK":""
}
VALID = set("ACDEFGHIKLMNPQRSTVWY")

def clean(seq):
    seq = re.sub(r"[^A-Z]", "", seq.upper())
    return "".join(c for c in seq if c in VALID)

def write_wrap(fh, s, w=60):
    for i in range(0, len(s), w): fh.write(s[i:i+w]+"\n")

def shard_writer(i):
    fp = OUT_DIR / f"bfvd_shard_{i:05d}.fasta"
    return fp, open(fp, "w", encoding="utf-8", buffering=1024*1024)

shard_idx, in_shard, total, files = 0, 0, 0, 0
out_path, out_fh = shard_writer(shard_idx)

def roll():
    global shard_idx, in_shard, out_path, out_fh
    out_fh.close()
    shard_idx += 1
    in_shard = 0
    out_path, out_fh = shard_writer(shard_idx)

def yield_pdbs(root: Path):
    # Faster than glob on Windows for huge trees
    stack = [root]
    while stack:
        d = stack.pop()
        with os.scandir(d) as it:
            for e in it:
                if e.is_dir(follow_symlinks=False):
                    stack.append(Path(e.path))
                elif e.is_file() and e.name.lower().endswith(".pdb"):
                    yield Path(e.path)

for pdb in yield_pdbs(IN_DIR):
    files += 1
    if files % 2000 == 0:
        print(f"[info] scanned {files:,} files | sequences so far: {total:,}", flush=True)
    try:
        chains = {}
        has_seqres = False
        with open(pdb, "r", encoding="utf-8", errors="ignore") as fh:
            for line in fh:
                if line.startswith("SEQRES"):
                    has_seqres = True
                    chain = (line[11].strip() or "X")[:1]
                    resnames = line[19:].strip().split()
                    chains.setdefault(chain, []).extend(resnames)

        if not has_seqres:
            # Fallback: extract sequence from ATOM CA records
            seen = set()
            with open(pdb, "r", encoding="utf-8", errors="ignore") as fh:
                for line in fh:
                    if not line.startswith("ATOM"): continue
                    if line[12:16].strip() != "CA": continue
                    chain = (line[21].strip() or "X")[:1]
                    resi  = line[22:27].strip()
                    key = (chain, resi)
                    if key in seen: continue
                    seen.add(key)
                    res3 = line[17:20].strip().upper()
                    chains.setdefault(chain, []).append(res3)

        for chain_id, res3s in chains.items():
            if not res3s: continue
            seq = "".join(AA3.get(r, "") for r in res3s)
            seq = clean(seq)
            if len(seq) < 30:  # skip tiny fragments
                continue
            header = f"{pdb.stem}_{chain_id}"
            out_fh.write(f">{header}\n")
            write_wrap(out_fh, seq)
            in_shard += 1; total += 1
            if in_shard >= SHARD_SIZE:
                roll()
    except Exception as e:
        print(f"[skip] {pdb}: {e}")

out_fh.close()
print(f"[done] sequences: {total:,} | shards: {shard_idx+1} | out: {OUT_DIR}")
```
