## Supplementary material for "*biocentral*: embedding-based protein predictions": predict_tmbed.py

```
import random

from biocentral_api import BiocentralAPI

biocentral_api = BiocentralAPI()


def read_fasta_with_predictions(input_path: str):
    with open(input_path, "r") as f:
        lines = f.readlines()
        seqs = {}
        preds = {}
        for idx, line in enumerate(lines):
            if line.startswith(">"):
                seq_id = line.strip().split(" ")[0][1:]
                seqs[seq_id] = lines[idx + 1].strip()
                preds[seq_id] = lines[idx + 2].strip()
        return seqs, preds


def remap_tmbed_predictions(tmbed_preds):
    remapped_preds = {}
    for key, preds in tmbed_preds.items():
        pred = preds[0]
        assert pred.model_name == "TMbed"
        assert pred.prediction_name == 'trans_membrane'
        assert pred.protocol == 'residue_to_class'
        remapped_preds[key] = pred.value.replace("o", ".").replace("i", ".")
    return remapped_preds


def check_predictions(loaded_prediction_data, server_preds):
    differences = []
    for key, pred in server_preds.items():
        loaded_prediction = loaded_prediction_data[key]
        if pred != loaded_prediction:
            differences.append((key, pred, loaded_prediction))
    print(f"Found {len(differences)} differences!")


def batched(dictionary, n):
    """
    Split a dictionary into n batches.

    Args:
        dictionary: The dictionary to batch
        n: Number of batches to create

    Yields:
        dict: Dictionary batches
    """
    items = list(dictionary.items())
    total_items = len(items)

    # Calculate batch size (distribute items evenly)
    batch_size = (total_items + n - 1) // n  # Ceiling division

    for i in range(n):
        start_idx = i * batch_size
        end_idx = min(start_idx + batch_size, total_items)

        if start_idx >= total_items:
            break

        yield dict(items[start_idx:end_idx])


model_names = ["TMbed"]
input_path = "bfvd_shard_00006.preds"
subset = False
loaded_sequence_data, loaded_prediction_data = read_fasta_with_predictions(input_path)

if subset:
    random.seed(42)
    test_subset_n = 10000
    random_selected_keys = random.sample(list(loaded_sequence_data.keys()), k=test_subset_n)
    subset_sequence_data = {random_key: loaded_sequence_data[random_key] for random_key in random_selected_keys}
    assert len(subset_sequence_data) == test_subset_n
else:
    subset_sequence_data = loaded_sequence_data

#_ = biocentral_api.embed(embedder_name="Rostlab/prot_t5_xl_uniref50", reduce=False,
#                         sequence_data=subset_sequence_data, ).run_with_progress()

result = {}
batch_id = 0
for batch in batched(subset_sequence_data, n=10):
    batch_result = biocentral_api.predict(model_names=model_names, sequence_data=batch).run_with_progress()
    result.update(batch_result)
    print(f"Batch {batch_id} done.")
    batch_id += 1

server_preds = remap_tmbed_predictions(result)
# print(server_preds)

check_predictions(loaded_prediction_data, server_preds)

with open(input_path + ".server", "w") as f:
    for key, pred in server_preds.items():
        f.write(f">{key}\n{loaded_sequence_data[key]}\n{pred}\n")
```
