## Supplementary material for "*biocentral*: embedding-based protein predictions": tm_envelope_significance_cli.py

```
# tm_envelope_significance_cli.py
"""
Significance & effect-size analysis for TM vs Envelope status.

What it does (without changing your labeling logic):
1) Builds 2x2 table (Enveloped vs Non-enveloped) x (TM vs no-TM).
2) Two-proportion z-test, 95% CI for difference, Relative Risk (RR), Odds Ratio (OR).
3) Optional chi-square test (if SciPy is installed).
4) Cohen's h (standardized difference in proportions).
5) Per-cluster (default: taxid) TM fractions + Wilcoxon rank-sum (if SciPy is installed).
6) Logistic regression (has_TM_segment ~ envelope + covariates) with cluster-robust SEs by cluster
   (if statsmodels is installed); covariates included if present: length, has_signal_peptide.

Outputs:
- Console logs
- If --report is given, writes:
  - a one-row CSV with all key stats (exactly like before, plus new fields),
  - a short Markdown report next to it (same stem, "_report.md").

Usage:
  py tm_envelope_significance_cli.py --input BFVD_with_envelope.csv --report out/tm_signif_summary.csv
Optional:
  --cluster-by taxid         (default "taxid")
  --alpha 0.05               (display only; CIs are 95%%)
  --log-level INFO|DEBUG|...
"""

from __future__ import annotations
from pathlib import Path
import argparse, logging, math, statistics
import pandas as pd

# ---------- helpers (no external deps) ----------
def norm(x):
    if pd.isna(x): return None
    return " ".join(str(x).strip().lower().split())

def read_csv_any(path: Path) -> pd.DataFrame:
    try:
        return pd.read_csv(path, dtype=str, sep=None, engine="python", low_memory=False)
    except Exception:
        return pd.read_csv(path, dtype=str, low_memory=False)

def ensure_has_tm(df: pd.DataFrame) -> pd.DataFrame:
    """Create has_TM_segment (0/1) from num_TMH/num_TMB if available; else from tm_class."""
    if "has_TM_segment" in df.columns:
        df["has_TM_segment"] = pd.to_numeric(df["has_TM_segment"], errors="coerce").fillna(0).astype(int)
        return df
    if "num_TMH" in df.columns and "num_TMB" in df.columns:
        df["num_TMH"] = pd.to_numeric(df["num_TMH"], errors="coerce").fillna(0).astype(int)
        df["num_TMB"] = pd.to_numeric(df["num_TMB"], errors="coerce").fillna(0).astype(int)
        df["has_TM_segment"] = ((df["num_TMH"] + df["num_TMB"]) > 0).astype(int)
        return df
    if "tm_class" in df.columns:
        t = df["tm_class"].astype(str).str.upper()
        df["has_TM_segment"] = t.isin(["TMH","TMB","TMH+TMB","TM"]).astype(int)
        return df
    raise SystemExit("[ERR] No TM columns found (need has_TM_segment OR (num_TMH & num_TMB) OR tm_class).")

def normalize_env(series: pd.Series) -> pd.Series:
    def map_env(v):
        v = norm(v)
        if v in (None, "", "nan"): return "Unknown"
        if v in {"enveloped","yes"}: return "Enveloped"
        if v in {"non-enveloped","non enveloped","no","nonenveloped"}: return "Non-enveloped"
        if v in {"variable","yes or no","predicted yes"}: return "Variable"
        # carry through custom categories
        return v.capitalize()
    return series.map(map_env)

def two_prop_z_test(a, b, c, d):
    """Compare p1=a/(a+b) vs p2=c/(c+d). Returns z, p(two-sided), diff, se_diff, (ci_lo, ci_hi), (p1, p2)."""
    n1 = a + b; n2 = c + d
    p1 = a / n1; p2 = c / n2
    p_pool = (a + c) / (n1 + n2)
    se = math.sqrt(p_pool * (1 - p_pool) * (1/n1 + 1/n2))
    z = (p1 - p2) / se if se > 0 else float("inf")
    p_two = math.erfc(abs(z) / math.sqrt(2))
    se_diff = math.sqrt(p1*(1-p1)/n1 + p2*(1-p2)/n2)
    zcrit = 1.959963984540054
    ci_lo = (p1 - p2) - zcrit * se_diff
    ci_hi = (p1 - p2) + zcrit * se_diff
    return z, p_two, (p1 - p2), se_diff, (ci_lo, ci_hi), (p1, p2)

def effect_sizes(a, b, c, d):
    """RR and OR with 95% CI; Haldane-Anscombe correction if zeros."""
    if 0 in (a,b,c,d): a+=0.5; b+=0.5; c+=0.5; d+=0.5
    n1 = a + b; n2 = c + d
    p1 = a / n1; p2 = c / n2
    rr = p1 / p2
    se_log_rr = math.sqrt(1/a - 1/n1 + 1/c - 1/n2)
    zcrit = 1.959963984540054
    rr_lo = math.exp(math.log(rr) - zcrit*se_log_rr)
    rr_hi = math.exp(math.log(rr) + zcrit*se_log_rr)
    or_ = (a*d) / (b*c)
    se_log_or = math.sqrt(1/a + 1/b + 1/c + 1/d)
    or_lo = math.exp(math.log(or_) - zcrit*se_log_or)
    or_hi = math.exp(math.log(or_) + zcrit*se_log_or)
    return (rr, rr_lo, rr_hi), (or_, or_lo, or_hi)

def cohens_h(p1: float, p2: float) -> float:
    return 2*math.asin(math.sqrt(p1)) - 2*math.asin(math.sqrt(p2))

# ---------- optional deps ----------
def try_scipy():
    try:
        from scipy.stats import chi2_contingency, mannwhitneyu
        return chi2_contingency, mannwhitneyu
    except Exception:
        return None, None

def try_statsmodels():
    try:
        import statsmodels.api as sm
        import statsmodels.formula.api as smf
        return sm, smf
    except Exception:
        return None, None

# ---------- per-cluster TM fractions & Wilcoxon ----------
def per_cluster_tm(df: pd.DataFrame, cluster_col: str, env_col: str) -> tuple[pd.DataFrame, pd.DataFrame]:
    """Return (per-cluster summary, filtered summary with unambiguous envelope)."""
    g = (
        df.groupby([cluster_col], dropna=False)
          .agg(
              n=('has_TM_segment','size'),
              tm=('has_TM_segment','sum'),
          )
          .reset_index()
    )
    # determine cluster envelope by majority label within cluster
    env_mode = (
        df.groupby([cluster_col, env_col], dropna=False)
          .size()
          .reset_index(name='count')
    )
    env_mode = env_mode.sort_values(['count'], ascending=False)
    env_first = env_mode.groupby(cluster_col).first().reset_index()[[cluster_col, env_col]]
    summary = g.merge(env_first, on=cluster_col, how='left')
    summary['tm_frac'] = summary['tm'] / summary['n']
    # mark ambiguous clusters (mixed labels)
    mix = (
        df.groupby(cluster_col)[env_col].nunique().reset_index(name='k')
    )
    summary = summary.merge(mix, on=cluster_col, how='left')
    unamb = summary[summary['k'] == 1].copy()
    ambig = summary[summary['k'] > 1].copy()
    return summary, unamb  # we use unambiguous for Wilcoxon

# ---------- logistic with cluster-robust SEs ----------
def logistic_cluster(df: pd.DataFrame, cluster_col: str) -> dict | None:
    sm, smf = try_statsmodels()
    if sm is None:
        logging.info("[logit] statsmodels not installed; skipping logistic regression.")
        return None
    work = df[df["__env__"].isin(["Enveloped","Non-enveloped"])].copy()
    work["env_bin"] = (work["__env__"] == "Enveloped").astype(int)
    # optional covariates
    covars = []
    if "length" in work.columns:
        work["length_num"] = pd.to_numeric(work["length"], errors="coerce")
        covars.append("length_num")
    if "has_signal_peptide" in work.columns:
        # attempt to coerce to binary
        hsp = work["has_signal_peptide"].astype(str).str.lower().isin(["1","true","yes","y","t"])
        work["hsp_bin"] = hsp.astype(int)
        covars.append("hsp_bin")
    formula = "has_TM_segment ~ env_bin" + ((" + " + " + ".join(covars)) if covars else "")
    try:
        model = smf.logit(formula, data=work).fit(disp=0)
        rob = model.get_robustcov_results(cov_type='cluster', groups=work[cluster_col])
        # pull env_bin coefficient
        coef = rob.params.get("env_bin", float("nan"))
        se = rob.bse.get("env_bin", float("nan"))
        z = coef/se if se and se==se else float("nan")
        p = 2* (1 - 0.5*math.erfc(-abs(z)/math.sqrt(2))) if z==z else float("nan")
        or_ = math.exp(coef) if coef==coef else float("nan")
        zcrit = 1.959963984540054
        lo = math.exp(coef - zcrit*se) if coef==coef and se==se else float("nan")
        hi = math.exp(coef + zcrit*se) if coef==coef and se==se else float("nan")
        return {
            "n": int(len(work)),
            "clusters": int(work[cluster_col].nunique()),
            "coef_env": coef, "se_env": se, "z_env": z, "p_env": p,
            "OR_env": or_, "OR_env_lo": lo, "OR_env_hi": hi,
            "formula": formula, "covariates": covars,
        }
    except Exception as e:
        logging.warning(f"[logit] model failed: {e}")
        return None

# ---------- main ----------
def main():
    ap = argparse.ArgumentParser(description="Significance tests for TM fraction: Enveloped vs Non-enveloped.")
    ap.add_argument("--input", required=True, help="Enriched CSV from labeling pipeline.")
    ap.add_argument("--alpha", type=float, default=0.05, help="Display alpha (CIs are 95%%).")
    ap.add_argument("--report", default=None, help="Optional CSV to write summary; a Markdown report with same stem is also written.")
    ap.add_argument("--cluster-by", default="taxid", help="Column to treat as cluster (default: taxid).")
    ap.add_argument("--log-level", default="INFO", choices=["DEBUG","INFO","WARNING","ERROR","CRITICAL"])
    args = ap.parse_args()

    logging.basicConfig(level=getattr(logging, args.log_level), format="%(asctime)s %(levelname)s %(message)s")

    df = read_csv_any(Path(args.input))
    if "envelope_status" in df.columns:
        env_col = "envelope_status"
    elif "envelope_status_ictv" in df.columns:
        env_col = "envelope_status_ictv"
    elif "envelope" in df.columns:
        env_col = "envelope"
    else:
        raise SystemExit("[ERR] Could not find envelope label column (envelope_status[_ictv] or envelope).")

    df = ensure_has_tm(df)
    df["__env__"] = normalize_env(df[env_col])

    # binary subset
    sub = df[df["__env__"].isin(["Enveloped","Non-enveloped"])].copy()
    if sub.empty:
        raise SystemExit("[ERR] No rows with Enveloped/Non-enveloped labels.")

    # 2x2 counts
    a = int(((sub["__env__"]=="Enveloped")     & (sub["has_TM_segment"]==1)).sum())
    b = int(((sub["__env__"]=="Enveloped")     & (sub["has_TM_segment"]==0)).sum())
    c = int(((sub["__env__"]=="Non-enveloped") & (sub["has_TM_segment"]==1)).sum())
    d = int(((sub["__env__"]=="Non-enveloped") & (sub["has_TM_segment"]==0)).sum())
    n1 = a + b; n2 = c + d

    logging.info("2x2 table (rows=group, cols=TM):")
    logging.info("                 TM=1     TM=0      total")
    logging.info(f"Enveloped      {a:8d} {b:8d} {n1:10d}")
    logging.info(f"Non-enveloped  {c:8d} {d:8d} {n2:10d}")

    # primary tests
    z, p_two, diff, se_diff, (ci_lo, ci_hi), (p1, p2) = two_prop_z_test(a, b, c, d)
    (rr, rr_lo, rr_hi), (or_, or_lo, or_hi) = effect_sizes(a, b, c, d)
    h = cohens_h(p1, p2)

    logging.info(f"\nTwo-proportion z-test (two-sided): z = {z:.3f}, p = {p_two:.3e}")
    logging.info(f"Proportion with TM | Enveloped = {p1:.4f}  Non-enveloped = {p2:.4f}")
    logging.info(f"Difference (E - NE) = {diff:.4f}  95% CI [{ci_lo:.4f}, {ci_hi:.4f}]")
    logging.info(f"Relative Risk (RR): {rr:.3f}  95% CI [{rr_lo:.3f}, {rr_hi:.3f}]")
    logging.info(f"Odds Ratio (OR):    {or_:,.3f}  95% CI [{or_lo:.3f}, {or_hi:.3f}]")
    logging.info(f"Cohen's h:          {h:.3f} (small–medium)")

    # optional chi-square
    chi2_fn, mw_fn = try_scipy()
    chi2 = chi2_p = chi2_df = None
    if chi2_fn:
        chi2, chi2_p, chi2_df, _ = chi2_fn([[a, b],[c, d]], correction=False)
        logging.info(f"\nChi-square test: X^2 = {chi2:.3f} (df={chi2_df}), p = {chi2_p:.3e}")
    else:
        logging.info("[chi2] SciPy not installed; skipping chi-square test.")

    # per-cluster TM fractions + Wilcoxon
    cluster_col = args.cluster_by
    wilc_n_env = wilc_n_non = None
    wilc_med_env = wilc_med_non = None
    wilc_u = wilc_p = None
    if cluster_col in df.columns:
        clust_all, clust_unamb = per_cluster_tm(df, cluster_col, "__env__")
        kept = clust_unamb[clust_unamb["__env__"].isin(["Enveloped","Non-enveloped"])].copy()
        env_tm = kept.loc[kept["__env__"]=="Enveloped", "tm_frac"].tolist()
        non_tm = kept.loc[kept["__env__"]=="Non-enveloped","tm_frac"].tolist()
        wilc_n_env, wilc_n_non = len(env_tm), len(non_tm)
        wilc_med_env = statistics.median(env_tm) if env_tm else float("nan")
        wilc_med_non = statistics.median(non_tm) if non_tm else float("nan")
        if mw_fn and wilc_n_env > 0 and wilc_n_non > 0:
            u, p = mw_fn(env_tm, non_tm, alternative="two-sided")
            wilc_u, wilc_p = float(u), float(p)
            logging.info(f"\nWilcoxon/Mann–Whitney on per-{cluster_col} TM fractions:")
            logging.info(f"n_env={wilc_n_env}, n_nonenv={wilc_n_non}, median_env={wilc_med_env:.4f}, median_nonenv={wilc_med_non:.4f}, U={u:.1f}, p={p:.3e}")
        else:
            logging.info(f"[wilcoxon] SciPy not installed or not enough clusters; skipping rank-sum. "
                         f"n_env={wilc_n_env}, n_nonenv={wilc_n_non}")
    else:
        logging.info(f"[wilcoxon] cluster column '{cluster_col}' not found; skipping per-cluster test.")

    # logistic regression with cluster-robust SEs
    logit_res = logistic_cluster(df, cluster_col) if cluster_col in df.columns else None
    if logit_res:
        logging.info("\nLogistic regression with cluster-robust SEs:")
        logging.info(f"formula: {logit_res['formula']}  | clusters={logit_res['clusters']}, n={logit_res['n']}")
        logging.info(f"env_bin OR = {logit_res['OR_env']:.3f}  95% CI [{logit_res['OR_env_lo']:.3f}, {logit_res['OR_env_hi']:.3f}]  "
                     f"(z={logit_res['z_env']:.3f}, p={logit_res['p_env']:.3e})")
        if logit_res["covariates"]:
            logging.info(f"covariates included: {', '.join(logit_res['covariates'])}")
    else:
        logging.info("[logit] not run (statsmodels missing or model failed).")

    # write report(s)
    if args.report:
        out_path = Path(args.report)
        out_path.parent.mkdir(parents=True, exist_ok=True)

        row = {
            "enveloped_tm": a, "enveloped_no_tm": b,
            "nonenv_tm": c, "nonenv_no_tm": d,
            "n_enveloped": n1, "n_non_enveloped": n2,
            "p_enveloped": p1, "p_non_enveloped": p2,
            "diff": diff, "diff_ci_lo": ci_lo, "diff_ci_hi": ci_hi,
            "z": z, "p_two_sided": p_two,
            "RR": rr, "RR_ci_lo": rr_lo, "RR_ci_hi": rr_hi,
            "OR": or_, "OR_ci_lo": or_lo, "OR_ci_hi": or_hi,
            "cohens_h": h,
            "chi2": chi2, "chi2_p": chi2_p, "chi2_df": chi2_df,
            "wilcoxon_U": wilc_u, "wilcoxon_p": wilc_p,
            "clusters_env": wilc_n_env, "clusters_nonenv": wilc_n_non,
            "median_tm_frac_env": wilc_med_env, "median_tm_frac_nonenv": wilc_med_non,
        }
        # add logistic if available
        if logit_res:
            row.update({
                "logit_n": logit_res["n"], "logit_clusters": logit_res["clusters"],
                "logit_OR_env": logit_res["OR_env"], "logit_OR_env_lo": logit_res["OR_env_lo"], "logit_OR_env_hi": logit_res["OR_env_hi"],
                "logit_z_env": logit_res["z_env"], "logit_p_env": logit_res["p_env"],
                "logit_formula": logit_res["formula"], "logit_covariates": ";".join(logit_res["covariates"]),
            })

        pd.DataFrame([row]).to_csv(out_path, index=False)
        logging.info(f"[write] summary CSV → {out_path}")

        # Markdown report
        md_path = out_path.with_name(out_path.stem + "_report.md")
        with md_path.open("w", encoding="utf-8") as f:
            f.write("# TM vs Envelope — Significance Summary\n\n")
            f.write(f"**Input:** `{Path(args.input).name}`  \n")
            f.write(f"**Binary groups:** Enveloped (n={n1}), Non-enveloped (n={n2})\n\n")
            f.write("## Two-proportion z-test\n")
            f.write(f"- p(TM|Env) = {p1:.4f}  vs  p(TM|NonEnv) = {p2:.4f}\n")
            f.write(f"- Difference = {diff:.4f}  (95% CI [{ci_lo:.4f}, {ci_hi:.4f}])\n")
            f.write(f"- z = {z:.3f}, p = {p_two:.3e}\n")
            f.write(f"- RR = {rr:.3f}  (95% CI [{rr_lo:.3f}, {rr_hi:.3f}])\n")
            f.write(f"- OR = {or_:.3f} (95% CI [{or_lo:.3f}, {or_hi:.3f}])\n")
            f.write(f"- Cohen's h = {h:.3f}\n\n")
            if chi2 is not None:
                f.write("## Chi-square test\n")
                f.write(f"- X^2 = {chi2:.3f} (df={chi2_df}), p = {chi2_p:.3e}\n\n")
            if cluster_col in df.columns:
                f.write(f"## Per-{cluster_col} TM fractions (unambiguous clusters)\n")
                if wilc_u is not None:
                    f.write(f"- n_env = {wilc_n_env}, n_nonenv = {wilc_n_non}  \n")
                    f.write(f"- median(TM frac) env = {wilc_med_env:.4f}, nonenv = {wilc_med_non:.4f}  \n")
                    f.write(f"- Wilcoxon/Mann–Whitney: U = {wilc_u:.1f}, p = {wilc_p:.3e}\n\n")
                else:
                    f.write("- Wilcoxon not run (SciPy missing or not enough clusters).\n\n")
            if logit_res:
                f.write("## Logistic regression (cluster-robust SEs)\n")
                f.write(f"- formula: `{logit_res['formula']}`  (clusters = {logit_res['clusters']}, n = {logit_res['n']})  \n")
                if logit_res["covariates"]:
                    f.write(f"- covariates: {', '.join(logit_res['covariates'])}  \n")
                f.write(f"- env_bin OR = {logit_res['OR_env']:.3f}  "
                        f"(95% CI [{logit_res['OR_env_lo']:.3f}, {logit_res['OR_env_hi']:.3f}]; "
                        f"z = {logit_res['z_env']:.3f}, p = {logit_res['p_env']:.3e})\n\n")
            f.write("---\n")
            f.write("_Notes: (i) Binary tests exclude Unknown/Variable groups. "
                    "(ii) Cluster tests treat each {cluster} as one unit to account for within-virus correlation. "
                    "(iii) All computations keep your original labeling logic unchanged._\n".format(cluster=cluster_col))
        logging.info(f"[write] Markdown report → {md_path}")

if __name__ == "__main__":
    main()
```
