## Supplementary material for "*biocentral*: embedding-based protein predictions": tmbed_convert.py

```
#!/usr/bin/env python
"""Convert TMbed prediction shards to JSONL or CSV format.
Usage:
    python tmbed_convert.py -i input_dir_or_files -o output.jsonl -f jsonl
  python tmbed_convert.py -i input_dir_or_files -o output.csv -f csv
"""
import re
import json
import csv
import argparse
from pathlib import Path

# TMbed label → (segment type, orientation)
LABEL_MAP = {
    "H": ("TMH", "in->out"),
    "h": ("TMH", "out->in"),
    "B": ("TMB", "in->out"),
    "b": ("TMB", "out->in"),
}


def extract_ids_from_header(header: str):
    """Extract accession and UniProt ID from FASTA header."""
    h = header[1:].strip()
    acc, uid = None, ""

    m = re.search(r"\|([A-NR-Z0-9]{6,10})\|([A-Z0-9_]+)", h)
    if m:
        acc = m.group(1)
        uid = m.group(2)
    else:
        toks = h.split()
        if toks:
            acc = toks[0]
        m2 = re.search(r"\b([A-Z0-9]{2,}_[A-Z0-9]{2,})\b", h)
        if m2:
            uid = m2.group(1)

    return acc, uid


def contiguous_runs(labels: str, allowed: set[str]):
    """Return list of (label_char, start_1based, end_exclusive)."""
    runs = []
    j, L = 0, len(labels)
    while j < L:
        c = labels[j]
        if c in allowed:
            k = j + 1
            while k < L and labels[k] == c:
                k += 1
            runs.append((c, j + 1, k))
            j = k
        else:
            j += 1
    return runs


def parse_pred_file(fp: Path):
    """Parse TMbed 3-line FASTA-like prediction file."""
    lines = fp.read_text(encoding="utf-8", errors="ignore").splitlines()
    for i in range(0, len(lines), 3):
        if i + 2 >= len(lines):
            break
        h, seq, lab = lines[i:i + 3]
        if not h.startswith(">"):
            continue
        acc, uid = extract_ids_from_header(h)
        if not acc:
            continue
        yield acc, uid, seq.strip(), lab.strip()


def collect_input_files(inputs):
    """Collect TMbed files from user input (files or directories)."""
    files = []
    for p in inputs:
        p = Path(p)
        if p.is_dir():
            files.extend(
                list(p.glob("*.pred")) +
                list(p.glob("*.preds")) +
                list(p.glob("*.preds.server"))
            )
        elif p.is_file():
            files.append(p)
        else:
            print(f"[WARN] Ignored invalid path: {p}")

    return sorted(set(files))


def process_all(input_paths, out_path: Path, out_format: str):
    """Process all TMbed files and write JSONL or CSV."""
    files = collect_input_files(input_paths)

    if not files:
        print("[ERROR] No TMbed input files found.")
        return

    print("[INFO] Processing files:")
    for f in files:
        print(f"       {f}")

    out_path.parent.mkdir(parents=True, exist_ok=True)

    seen_acc = set()
    rows = []  # store for CSV if needed

    if out_format == "jsonl":
        out_f = out_path.open("w", encoding="utf-8")

    for fp in files:
        for acc, uid, seq, lab in parse_pred_file(fp):
            if acc in seen_acc:
                continue
            seen_acc.add(acc)

            L = len(seq)

            # parse segments
            runs = contiguous_runs(lab, set("HhBbS"))
            tmh_count = 0
            tmb_count = 0
            sp_count = 0
            segments = []

            for c, start_1based, end_exclusive in runs:
                if c in "HhBb":
                    seg_type, orientation = LABEL_MAP[c]
                    segments.append({
                        "type": seg_type,
                        "start": start_1based,
                        "end": end_exclusive,
                        "orientation": orientation
                    })
                    if seg_type == "TMH":
                        tmh_count += 1
                    else:
                        tmb_count += 1
                elif c == "S":
                    sp_count += 1

            # determine tm_class
            if tmh_count > 0 and tmb_count == 0:
                tm_class = "TMH"
            elif tmb_count > 0 and tmh_count == 0:
                tm_class = "TMB"
            elif tmh_count > 0 and tmb_count > 0:
                tm_class = "Mixed"
            else:
                tm_class = "None"

            has_signal_peptide = sp_count > 0

            record = {
                "uniprot_accession": acc,
                "length": L,
                "tm_class": tm_class,
                "has_signal_peptide": has_signal_peptide,
                "num_TMH": tmh_count,
                "num_TMB": tmb_count,
                "segments": segments,
                "sequence": seq,
                "labels": lab,
            }

            if out_format == "jsonl":
                json.dump(record, out_f)
                out_f.write("\n")

            elif out_format == "csv":
                rows.append({
                    "uniprot_accession": acc,
                    "length": L,
                    "tm_class": tm_class,
                    "has_signal_peptide": has_signal_peptide,
                    "num_TMH": tmh_count,
                    "num_TMB": tmb_count,
                    "segments": json.dumps(segments),
                    "sequence": seq,
                    "labels": lab,
                })

    # Write CSV if needed
    if out_format == "csv":
        with out_path.open("w", newline="", encoding="utf-8") as f:
            writer = csv.DictWriter(
                f,
                fieldnames=[
                    "uniprot_accession",
                    "length",
                    "tm_class",
                    "has_signal_peptide",
                    "num_TMH",
                    "num_TMB",
                    "segments",
                    "sequence",
                    "labels",
                ],
            )
            writer.writeheader()
            writer.writerows(rows)

    if out_format == "jsonl":
        out_f.close()

    print(f"[INFO] Done. Output written to {out_path}")


def main():
    parser = argparse.ArgumentParser(
        description="Convert TMbed prediction shards to JSONL or CSV."
    )
    parser.add_argument(
        "-i", "--input",
        nargs="+",
        required=True,
        help="Input TMbed files or directories."
    )
    parser.add_argument(
        "-o", "--output",
        required=True,
        help="Output file path."
    )
    parser.add_argument(
        "-f", "--format",
        choices=["jsonl", "csv"],
        required=True,
        help="Output format."
    )

    args = parser.parse_args()

    process_all(args.input, Path(args.output), args.format)


if __name__ == "__main__":
    main()
```
