## Supplementary material for "*biocentral*: embedding-based protein predictions": biocentral_som.docx

ALGORITHM 1: Model integration into the biocentral service

(1) convert the model to ONNX; (2) upload the ONNX artifact; (3) implement the Python class with pre/postprocessing and the ONNX model call. If functions from the original model repository are necessary: create a new deployment branch in the model repository and delete all unnecessary files and dependencies – this allows to re-use existing code while keeping the dependencies minimal; (4) add triton-specific files for optimized inference; (5) submit a pull request in the <https://github.com/biocentral/biocentral_server> repository with the code and access to the ONNX model.

This approach avoids introducing additional ML framework or Python version dependencies, thereby reducing deprecation and compatibility issues.

ALGORITHM 2: Reproducing the ExoTox predictor

(1) load the training and test sequences and labels from the ExoTox supplementary material; (2) create a validation subset from the provided training set (10% of the training set); (3) convert the data to biocentral-compatible classes (SequenceTrainingData); (4) create the following training configuration:

{

"embedder_name": “Rostlab/prot_t5_xl_uniref50”,

"protocol": "sequence_to_class",

"model_choice": "FNN",

"dimension_reduction_method": "pca",

"n_reduced_components": 20,

}; (5) create an instance of the biocentral api client; (6) use the biocentral-api’s train method with the sequence training data and the training configuration; (7) compare and visualize results.

We provide these steps, including training of baselines, as a ready-to-use Jupyter notebook at: <https://github.com/biocentral/biocentral_api/blob/main/python/examples/exotox_reproduction/exotox_reproduction.ipynb>.

Figure 1: TMbed/BFVD predictions

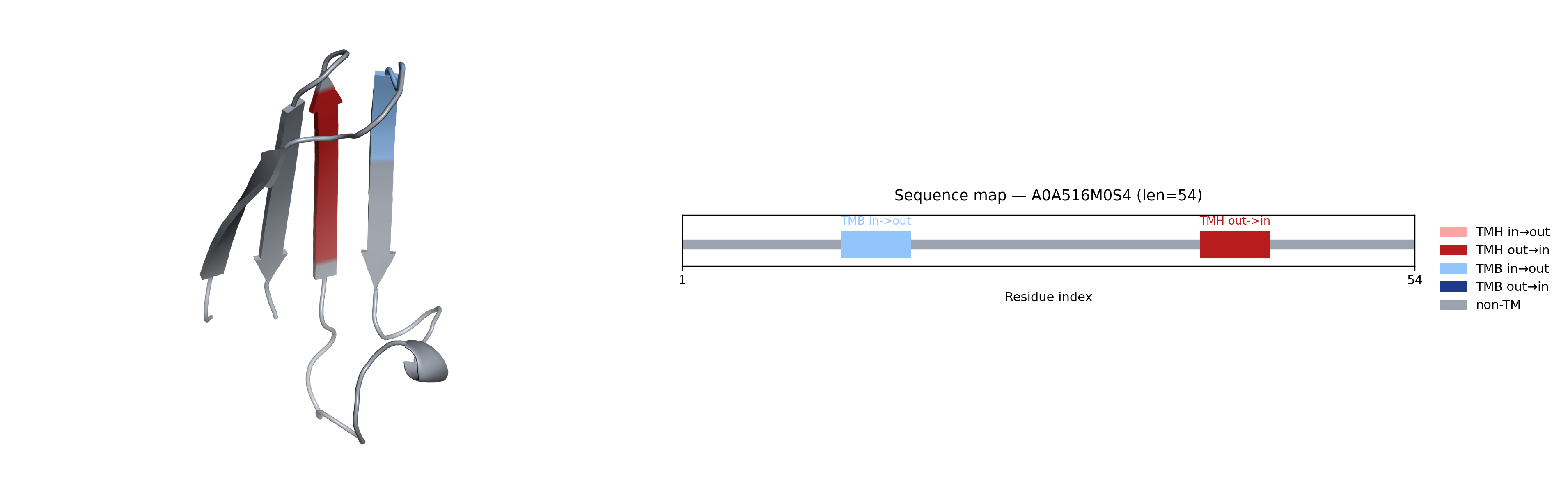

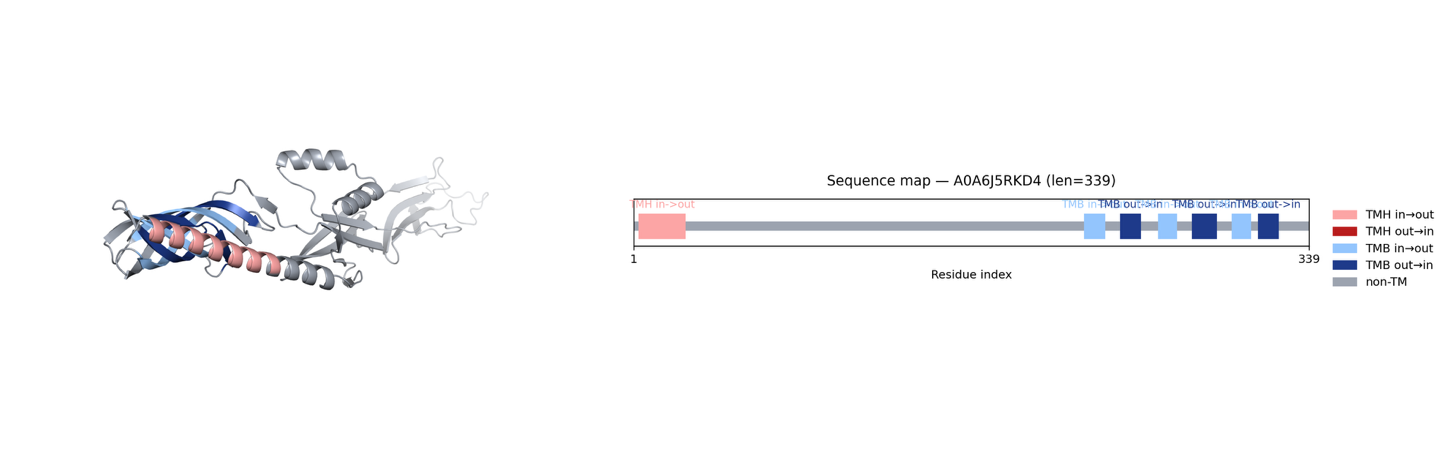

**Figure 1** **Visual comparison of TMH+TMB predictions for high-confidence BFVD proteins.**
Shown are the shortest (A0A516M0S4; 54 residues; average pLDDT ≈ 88; upper panel) and longest (A0A6J5RKD4; 339 residues; average pLDDT ≈ 85; lower panel) proteins within the BFVD subset that simultaneously (i) received both TMH and TMB predictions from TMbed and (ii) reached an average pLDDT above 70. In A0A516M0S4, TMbed predicts a transmembrane segment at an implausible position, indicating substantial confusion. In A0A6J5RKD4, the β-barrel prediction appears more coherent, whereas the predicted helix is likely a misclassified signal peptide. Together, these examples highlight atypical and readily identifiable prediction artefacts despite otherwise high structural confidence.
