## Supplementary material for "*biocentral*: embedding-based protein predictions": exotox_conversion.ipynb

```
# %% [code]
# Import dependencies
import pickle
import numpy as np
import onnxruntime as rt

from skl2onnx import to_onnx
from collections import Counter
from exotox_utils import read_fasta, read_labels
from biocentral_api import BiocentralAPI, CommonEmbedder

# %% [code]
# Load the predictor
predictor_file = "exotox_files/sklearn_svcPC20_SST30_CV10_embeddingsProtT5"
with open(predictor_file, 'rb') as f:
    predictor = pickle.load(f)

# %% [code]
# Get test set embeddings
test_set_file = "exotox_files/X_test_SST30.fasta"
test_seqs = read_fasta(test_set_file)
test_labels = read_labels("exotox_files/y_test_SST30.csv")

embedder_name = CommonEmbedder.ProtT5
biocentral_api = BiocentralAPI().wait_until_healthy()

embeddings = biocentral_api.embed(embedder_name=embedder_name, sequence_data=test_seqs, reduce=True).run_with_progress()

# %% [code]
# Run predictor on the test set for verification of results
X = list(embeddings.values())
Y = [int(test_labels[seq_id]) for seq_id in embeddings.keys()]
y_pred = predictor.predict(X)
y_probas = predictor.predict_proba(X)[:, 1]
count = Counter(y_pred)
accuracy = sum([1 for idx, y in enumerate(y_pred) if int(y) == int(Y[idx])]) / len(Y)

print(count)
print(f"Accuracy: {accuracy}")

# %% [code]
# Convert to ONNX
X_array = np.array(X, dtype=np.float32)

# Specify options to get tensor outputs instead of sequences
options = {id(predictor): {'zipmap': False}}

# Convert with proper output specification
onx = to_onnx(
    predictor,
    X_array[:1],
    options=options,
    target_opset=18
)

with open("exotox.onnx", "wb") as f:
    f.write(onx.SerializeToString())

# %% [code]
# Run ONNX model on test set
sess = rt.InferenceSession("exotox.onnx", providers=["CPUExecutionProvider"])
input_name = sess.get_inputs()[0].name
label_name = sess.get_outputs()[0].name
print(label_name)
pred_onnx = sess.run([label_name], {input_name: X_array.astype(np.float32)})[0]

count_onnx = Counter(pred_onnx)
accuracy_onnx = sum([1 for idx, y in enumerate(pred_onnx) if int(y) == int(Y[idx])]) / len(Y)

print(count_onnx)
print(f"Accuracy: {accuracy_onnx}")

assert accuracy == accuracy_onnx

# %% [code]
```
