## Supplementary material for "*biocentral*: embedding-based protein predictions": classifying_unknown_proteins.py

```
#!/usr/bin/env python3
# coding: utf-8


########################################################################################################################
# Author: Tanja Krüger
# Aim: This file reuses a finished trained predictor for new predictions
# Input1: a trained predictor.
#   Where:You can find those under the Predictor folder of this project.
#   Recommended model: sklearn_svcPC20_SST30_CV10_embeddingsProtT5.
#   Alternatives: The same folder has other alternatives. Everything that starts with sklearn in the beginning of the
#       name is compatible with this script. Careful what model you choose, some were trained on scrambled sequences.
#       Predictions will be significantly less good - these models have "allscr" in their name.
# Input2: X: embeddings from the sequences you want to predict.
#   The embeddings should be in h5 format. Here are the base instructions how to get the embeddings. For more detail go
#   to the makefile. #Create embeddings from these fasta files:Therefore go to the following link:
#   https://colab.research.google.com/drive/1TUj-ayG3WO52n5N50S7KH9vtt6zRkdmj?usp=sharing#scrollTo=tRe7CfuqFFmY and
#   upload the file. Run the first two cells inside the notebook and then move the uploaded file to the protT5 folder.
#   Turn off the per_residue and the sec_struct options and keep the per_protein turned on. Change the seq_path to
#   include the filename of interest. Run all the cells.
# Output:The predictions

########################################################################################################################

import numpy as np

import pickle
import h5py
import pandas as pd
import re, argparse, csv, collections,random
from collections import Counter
from datetime import datetime
# #################################################################################################
# Option depending where the user wants the run the code form, default running the code with make from the project folder.
cl=""
# If one wants to execute this file from the Code/python folder uncomment the next line.
cl="../../"

########################################################################################################################
# Get the arguments from the command line.
parser = argparse.ArgumentParser(prog="classifying_unknown_proteins",
                                 description=" predicted toxicity to proteins in embeddings format")
parser.add_argument("pred",
                    type=str,
                    help="trained predictor")
parser.add_argument("X",
                    type=str,
                    help="secreted proteins of unknown toxicity in embeddings format")

# animal_tox_animal_non_tox_embeddings.h5
# example_phage_proteins_handpicked_embeddings.h5
# animal_controls_per_protein_embeddings.h5
# animal_toxins_per_protein_embeddings.h5
# bacterial_controls_testset_per_protein_embeddings.h5
# bacterial_non_toxins_nonsecreted_per_protein_embeddings.h5
# bacterial_toxins_testset_per_protein_embeddings.h5
# eukaryot_infecting_viruses_per_protein_embeddings.h5
# example_phage_proteins_handpicked_embeddings.h5
# example_phage_proteins_spike_in_baseplate.pdb


args = parser.parse_args()

# Extract information from the provided parameters
# Number of cross validations in optimizing the predictor
cv=re.search("CV(\d+)_", args.pred).group(1)
# folder where the results are saved
where=re.search("Predictor/(.*)_(.*)_SST(\d+)_", args.pred).group(1)
# mmseqs2 reduction level
sst_level=re.search("SST(\d+)", args.pred).group(1)
# the type of embedding during training
embedding_type_train = re.search("Predictor/(.*)_(.*)_SST(\d+)_CV(\d+)_(.*)",  args.pred).group(5)
# the model architecture
architecture=re.search("Predictor/(.*)_(.*)_SST(\d+)_", args.pred).group(2)


# Make the results reproducible
random_seed=7
random_state=7


########################################################################################################################
# Step 1: Open data
# Step 1.1: Open the predictor
with open(args.pred, 'rb') as f:
    predictor = pickle.load(f)

# Step 1.2: Open the embeddings
with h5py.File(args.X, "r") as f:
    embs = dict((k, list(f[k])) for k in f)
    X=pd.DataFrame(embs).T
    X.index=[i.split(' ',1)[0] for i in f]


# Step 2: run the test file on the imported predictor
y_pred = predictor.predict(X)
y_probas =predictor.predict_proba(X)[:, 1]
count = Counter(y_pred)

print(count)
print(y_pred)
print(y_probas)
```
