## Supplementary material for "*biocentral*: embedding-based protein predictions": exotox_reproduction.ipynb

```
# %% [code]
# Import dependencies
import numpy as np
import matplotlib.pyplot as plt

from copy import deepcopy
from collections import Counter
from biocentral_api import BiocentralAPI, CommonEmbedder, BiocentralPredictionModel
from exotox_utils import do_bootstrapping, read_fasta, read_labels, create_training_data, split_train_val

# %% [code]
# Input Data
train_fasta = "exotox_files/X_train_SST30.fasta"
test_fasta = "exotox_files/X_test_SST30.fasta"
train_csv = "exotox_files/y_train_SST30.csv"
test_csv = "exotox_files/y_test_SST30.csv"


# %% [code]
# Create input data
val_size = 0.1  # Original paper used k-fold cross validation, we use hold-out cross validation with validation dataset

train_seqs = read_fasta(train_fasta)
train_labels = read_labels(train_csv)
test_seqs = read_fasta(test_fasta)
test_labels = read_labels(test_csv)

train_seqs, val_seqs, train_labels, val_labels  = split_train_val(train_seqs, train_labels, val_size)

training_data = create_training_data(train_seqs, val_seqs, test_seqs, train_labels, val_labels, test_labels)
sequence_data = {data_point.seq_id: data_point.sequence for data_point in training_data}

# %% [code]
# Show statistics
print(f"Total sequences: {len(training_data)}")
print(f"Training sequences: {len(train_seqs)}")
print(f"Validation sequences: {len(val_seqs)}")
print(f"Test sequences: {len(test_seqs)}")

assert len(test_seqs) == 324
assert len(train_seqs) < 2020
assert len(val_seqs) > 200

print(f"Percentage of toxins (1) in training_data: {sum([int(label) for label in train_labels.values()]) / len(train_labels)} ")
print(f"Percentage of toxins (1) in validation_data: {sum([int(label) for label in val_labels.values()]) / len(val_labels)}")
print(f"Percentage of toxins (1) in test_data: {sum([int(label) for label in test_labels.values()]) / len(test_labels)}")

# %% [code]
# Create client
client = BiocentralAPI().wait_until_healthy()

# %% [code]
# Embed OHE
reduce = True  # per-protein embeddings for per-protein task

embedder_name_ohe = CommonEmbedder.ONE_HOT_ENCODING
embeddings_ohe = client.embed(embedder_name=embedder_name_ohe, reduce=True, sequence_data=sequence_data).run_with_progress()

# %% [code]
# Embed ProtT5
embedder_name_prott5 = CommonEmbedder.ProtT5
embeddings_prott5 = client.embed(embedder_name=embedder_name_prott5, reduce=True, sequence_data=sequence_data).run_with_progress()

# %% [code]
# Train OHE Model
config_baseline_ohe = {
    "embedder_name": embedder_name_ohe,
    "protocol": "sequence_to_class",
    "model_choice": "FNN"
}

result_ohe = client.train(config=config_baseline_ohe, training_data=training_data).run_with_progress()

# %% [code]
# Train Random Embedder Model
config_baseline_random_embeddings = deepcopy(config_baseline_ohe)
config_baseline_random_embeddings["embedder_name"] = CommonEmbedder.RANDOM_EMBEDDER

result_random_embeddings = client.train(config=config_baseline_random_embeddings, training_data=training_data).run_with_progress()

# %% [code]
# Train AAOntology Model
config_baseline_aaontology = deepcopy(config_baseline_ohe)
config_baseline_aaontology["embedder_name"] = CommonEmbedder.AAOntology

result_aaontology = client.train(config=config_baseline_aaontology, training_data=training_data).run_with_progress()

# %% [code]
# Train BLOSUM62 Model
config_baseline_blosum62 = deepcopy(config_baseline_ohe)
config_baseline_blosum62["embedder_name"] = CommonEmbedder.BLOSUM62

result_blosum62 = client.train(config=config_baseline_blosum62, training_data=training_data).run_with_progress()

# %% [code]
# Train ProtT5 Model
config = {
    "embedder_name": embedder_name_prott5,
    "protocol": "sequence_to_class",
    "model_choice": "FNN",
    "dimension_reduction_method": "pca",
    "n_reduced_components": 20,
}

result_prott5 = client.train(config=config, training_data=training_data).run_with_progress()
print(result_prott5)

# %% [code]
# Run biocentral ExoTox predictor on server for comparison
predictions_exotox_server = client.predict(model_names=[BiocentralPredictionModel.EXOTOX], sequence_data=test_seqs).run_with_progress()
print(predictions_exotox_server)

# %% [code]
reamapped_server_predictions = {seq_id: 0 if "NOT" in pred[0].value else 1 for seq_id, pred in predictions_exotox_server.items()}
count_server = Counter(reamapped_server_predictions.values())

exotox_server_preds = np.array(list(reamapped_server_predictions.values()))
server_test_labels_sorted = np.array([int(test_labels[seq_id]) for seq_id in reamapped_server_predictions.keys()])

result_exotox_server = do_bootstrapping(y_true=server_test_labels_sorted, y_pred=exotox_server_preds, iterations=30, sample_size=-1, confidence_level=0.05)
print(count_server)
print(result_exotox_server)

# %% [code]
# Visualize model results

# Extract MCC values from the dictionaries
exotox_onnx_mcc = result_exotox_server['mcc']['mean']
exotox_onnx_mcc_lower = result_exotox_server['mcc']['lower']
exotox_onnx_mcc_upper = result_exotox_server['mcc']['upper']

# ProtT5 results
prott5_mcc = result_prott5['test_results']['test']['bootstrapping']['results']['matthews-corr-coeff']['mean']
prott5_mcc_lower = result_prott5['test_results']['test']['bootstrapping']['results']['matthews-corr-coeff']['lower']
prott5_mcc_upper = result_prott5['test_results']['test']['bootstrapping']['results']['matthews-corr-coeff']['upper']

prott5_random_mcc = result_prott5['test_results']['test']['test_baselines']['random_model']['results']['matthews-corr-coeff']['mean']
prott5_random_lower = result_prott5['test_results']['test']['test_baselines']['random_model']['results']['matthews-corr-coeff']['lower']
prott5_random_upper = result_prott5['test_results']['test']['test_baselines']['random_model']['results']['matthews-corr-coeff']['upper']

# OHE results
ohe_mcc = result_ohe['test_results']['test']['bootstrapping']['results']['matthews-corr-coeff']['mean']
ohe_mcc_lower = result_ohe['test_results']['test']['bootstrapping']['results']['matthews-corr-coeff']['lower']
ohe_mcc_upper = result_ohe['test_results']['test']['bootstrapping']['results']['matthews-corr-coeff']['upper']

## AAOntology results
aaontology_mcc = result_aaontology['test_results']['test']['bootstrapping']['results']['matthews-corr-coeff']['mean']
aaontology_mcc_lower = result_aaontology['test_results']['test']['bootstrapping']['results']['matthews-corr-coeff']['lower']
aaontology_mcc_upper = result_aaontology['test_results']['test']['bootstrapping']['results']['matthews-corr-coeff']['upper']

## BLOSUM62 results
blosum62_mcc = result_blosum62['test_results']['test']['bootstrapping']['results']['matthews-corr-coeff']['mean']
blosum62_mcc_lower = result_blosum62['test_results']['test']['bootstrapping']['results']['matthews-corr-coeff']['lower']
blosum62_mcc_upper = result_blosum62['test_results']['test']['bootstrapping']['results']['matthews-corr-coeff']['upper']

# Random Embedder results
random_embedder_mcc = result_random_embeddings['test_results']['test']['bootstrapping']['results']['matthews-corr-coeff']['mean']
random_embedder_mcc_lower = result_random_embeddings['test_results']['test']['bootstrapping']['results']['matthews-corr-coeff']['lower']
random_embedder_mcc_upper = result_random_embeddings['test_results']['test']['bootstrapping']['results']['matthews-corr-coeff']['upper']

ohe_random_mcc = result_ohe['test_results']['test']['test_baselines']['random_model']['results']['matthews-corr-coeff']['mean']
ohe_random_lower = result_ohe['test_results']['test']['test_baselines']['random_model']['results']['matthews-corr-coeff']['lower']
ohe_random_upper = result_ohe['test_results']['test']['test_baselines']['random_model']['results']['matthews-corr-coeff']['upper']

# Shared baselines (one_only and zero_only are the same for both)
one_only_mcc = result_prott5['test_results']['test']['test_baselines']['one_only']['results']['matthews-corr-coeff']['mean']
one_only_lower = result_prott5['test_results']['test']['test_baselines']['one_only']['results']['matthews-corr-coeff']['lower']
one_only_upper = result_prott5['test_results']['test']['test_baselines']['one_only']['results']['matthews-corr-coeff']['upper']

zero_only_mcc = result_prott5['test_results']['test']['test_baselines']['zero_only']['results']['matthews-corr-coeff']['mean']
zero_only_lower = result_prott5['test_results']['test']['test_baselines']['zero_only']['results']['matthews-corr-coeff']['lower']
zero_only_upper = result_prott5['test_results']['test']['test_baselines']['zero_only']['results']['matthews-corr-coeff']['upper']

# Prepare data for plotting
models = ['ExoTox', 'Reproduced\n(ProtT5)', 'One Hot Encodings', 'BLOSUM62', 'AAOntology','Random\nEmbeddings', 'Untrained Model\n(ProtT5)'] #,'Random\n(OHE)', 'One Only', 'Zero Only']
mcc_means = [exotox_onnx_mcc, prott5_mcc, ohe_mcc, blosum62_mcc, aaontology_mcc, random_embedder_mcc, prott5_random_mcc] #ohe_random_mcc, one_only_mcc, zero_only_mcc]

# Calculate error bars (distance from mean to lower/upper bounds)
error_lower = [
    exotox_onnx_mcc - exotox_onnx_mcc_lower,
    prott5_mcc - prott5_mcc_lower,
    ohe_mcc - ohe_mcc_lower,
    blosum62_mcc - blosum62_mcc_lower,
    aaontology_mcc - aaontology_mcc_lower,
    random_embedder_mcc - random_embedder_mcc_lower,
    prott5_random_mcc - prott5_random_lower,
    #ohe_random_mcc - ohe_random_lower,
    #one_only_mcc - one_only_lower,
    #zero_only_mcc - zero_only_lower
]

error_upper = [
    exotox_onnx_mcc_upper - exotox_onnx_mcc,
    prott5_mcc_upper - prott5_mcc,
    ohe_mcc_upper - ohe_mcc,
    blosum62_mcc_upper - blosum62_mcc,
    aaontology_mcc_upper - aaontology_mcc,
    random_embedder_mcc_upper - random_embedder_mcc,
    prott5_random_upper - prott5_random_mcc,
    #ohe_random_upper - ohe_random_mcc,
    #one_only_upper - one_only_mcc,
    #zero_only_upper - zero_only_mcc
]

mcc_means = list(map(lambda x: round(x, 3), mcc_means))
error_lower = list(map(lambda x: round(x, 3), error_lower))
error_upper = list(map(lambda x: round(x, 3), error_upper))


errors = [error_lower, error_upper]

# Create the plot
fig, ax = plt.subplots(figsize=(12, 7))

# Create bars
x_pos = np.arange(len(models))
bars = ax.bar(x_pos, mcc_means, yerr=errors, capsize=8, color='darkgrey',
                alpha=0.8, edgecolor='black', linewidth=1.5,
               error_kw={'linewidth': 2, 'elinewidth': 2})

# Customize the plot
ax.set_ylabel('Matthews Correlation Coefficient (MCC)', fontsize=14, fontweight='bold')
ax.set_xlabel('Models', fontsize=14, fontweight='bold')
#ax.set_title('MCC Comparison: ExoTox vs. Reproduced ProtT5 vs Baselines', fontsize=16, fontweight='bold', pad=20)
ax.set_xticks(x_pos)
ax.set_xticklabels(models, fontsize=12)
ax.tick_params(axis='y', labelsize=11)

# Add grid for better readability
ax.grid(axis='y', alpha=0.3, linestyle='--', linewidth=0.7)
ax.set_axisbelow(True)

# Add a horizontal line at y=0 for reference
ax.axhline(y=0, color='black', linestyle='-', linewidth=0.8, alpha=0.5)

# Add value labels on top of bars
for i, (bar, mean_val) in enumerate(zip(bars, mcc_means)):
    height = bar.get_height()
    y_pos = height + errors[1][i] + 0.02 if height >= 0 else height - errors[0][i] - 0.02
    ax.text(bar.get_x() + bar.get_width()/2., y_pos,
            f'{mean_val:.3f}',
            ha='center', va='bottom' if height >= 0 else 'top',
            fontsize=10, fontweight='bold')

# Adjust layout and display
plt.tight_layout()
plt.savefig('exotox_model_barplot.png', dpi=400, bbox_inches='tight', facecolor='white')
plt.show()

# Print summary statistics
print("MCC Summary Statistics:")
print("=" * 60)
print(f"Exotox Model:        {exotox_onnx_mcc:.4f} [{exotox_onnx_mcc_lower:.4f}, {exotox_onnx_mcc_upper:.4f}]")
print(f"ProtT5 Model:        {prott5_mcc:.4f} [{prott5_mcc_lower:.4f}, {prott5_mcc_upper:.4f}]")
print(f"OHE Model:           {ohe_mcc:.4f} [{ohe_mcc_lower:.4f}, {ohe_mcc_upper:.4f}]")
print(f"AAOntology Model:           {aaontology_mcc:.4f} [{aaontology_mcc_lower:.4f}, {aaontology_mcc_upper:.4f}]")
print(f"Random Embedder Model:           {random_embedder_mcc:.4f} [{random_embedder_mcc_lower:.4f}, {random_embedder_mcc_upper:.4f}]")
print(f"Random Initialized Model (ProtT5):     {prott5_random_mcc:.4f} [{prott5_random_lower:.4f}, {prott5_random_upper:.4f}]")
print(f"Random Initialized Model (OHE):        {ohe_random_mcc:.4f} [{ohe_random_lower:.4f}, {ohe_random_upper:.4f}]")
print(f"One Only Baseline:   {one_only_mcc:.4f} [{one_only_lower:.4f}, {one_only_upper:.4f}]")
print(f"Zero Only Baseline:  {zero_only_mcc:.4f} [{zero_only_lower:.4f}, {zero_only_upper:.4f}]")

# %% [code]
```
