## Supplementary material for "*biocentral*: embedding-based protein predictions": exotox_utils.py

```
# Utility functions for conversion and reproduction
import numpy as np
import pandas as pd

from typing import List, Dict, Tuple, Callable
from biocentral_api import SequenceTrainingData
from sklearn.model_selection import train_test_split
from sklearn.metrics import accuracy_score, matthews_corrcoef


def read_fasta(input_path: str):
    with open(input_path, "r") as f:
        lines = f.readlines()
        seqs = {}
        for line in lines:
            if line.startswith(">"):
                seq_id = line.strip().split(" ")[0][1:]
                seqs[seq_id] = ""
            else:
                seqs[seq_id] += line.strip()
        return seqs


def read_labels(input_path: str):
    df = pd.read_csv(input_path)
    labels = {}
    for _, row in df.iterrows():
        labels[row["id"]] = str(row["label"])
    return labels


def split_train_val(train_seqs, train_labels, val_size: float):
    """
    Split training sequences and labels into train and validation sets.

    Args:
        train_seqs: Dictionary mapping sequence IDs to sequences
        train_labels: Dictionary mapping sequence IDs to labels
        val_size: Fraction of data to use for validation (0.0 to 1.0)

    Returns:
        Tuple of (train_seqs_dict, val_seqs_dict, train_labels_dict, val_labels_dict)
    """
    # Extract aligned lists of IDs, sequences, and labels
    seq_ids = list(train_seqs.keys())
    labels = [train_labels[seq_id] for seq_id in seq_ids]

    # Split the data
    train_ids, val_ids, train_labels_list, val_labels_list = train_test_split(
        seq_ids, labels, test_size=val_size,
        random_state=42, stratify=labels
    )

    # Reconstruct dictionaries
    train_seqs_dict = {seq_id: train_seqs[seq_id] for seq_id in train_ids}
    val_seqs_dict = {seq_id: train_seqs[seq_id] for seq_id in val_ids}
    train_labels_dict = {seq_id: train_labels[seq_id] for seq_id in train_ids}
    val_labels_dict = {seq_id: train_labels[seq_id] for seq_id in val_ids}

    return train_seqs_dict, val_seqs_dict, train_labels_dict, val_labels_dict


def create_training_data(train_seqs, val_seqs, test_seqs, train_labels, val_labels, test_labels) -> List[
    SequenceTrainingData]:
    seqs_labels_sets = [(train_seqs, train_labels, "train"), (val_seqs, val_labels, "val"),
                        (test_seqs, test_labels, "test")]
    training_data = []
    for seqs, labels, set_name in seqs_labels_sets:
        for seq_id, seq in seqs.items():
            label = labels[seq_id]
            data_point = SequenceTrainingData(seq_id=seq_id, sequence=seq, label=label, set=set_name, mask=None)
            training_data.append(data_point)
    return training_data


def _calculate_metrics(y_true, y_pred):
    accuracy = accuracy_score(y_true, y_pred)
    mcc = matthews_corrcoef(y_true, y_pred)
    return accuracy, mcc


def _get_mean_and_confidence_bounds(values: np.ndarray, confidence_level: float) -> Tuple[float, float, float]:
    """
    Calculates the mean and confidence range for the given values. Used for bootstrapping error reporting.

    :param values: Array of metric values from bootstrap iterations
    :param confidence_level: Confidence level for result confidence intervals (0.05 => 95% percentile)
    :return: Tuple: mean, lower_bound, upper_bound
    """
    if not 0 < confidence_level < 1:
        raise ValueError(f"Confidence level must be between 0 and 1, given: {confidence_level}!")

    mean = np.mean(values)

    # Calculate percentiles from actual distribution
    lower_percentile = (confidence_level / 2) * 100
    upper_percentile = (1 - confidence_level / 2) * 100

    lower_bound = np.percentile(values, lower_percentile)
    upper_bound = np.percentile(values, upper_percentile)

    return mean, lower_bound, upper_bound


def do_bootstrapping(
        y_true: np.ndarray,
        y_pred: np.ndarray,
        iterations: int = 1000,
        sample_size: int = -1,
        confidence_level: float = 0.05,
        random_seed: int = None
) -> Dict[str, Dict[str, float]]:
    """
    Perform bootstrapping to estimate confidence intervals for metrics.

    :param y_true: True labels (numpy array)
    :param y_pred: Predicted labels (numpy array)
    :param iterations: Number of bootstrap iterations to perform
    :param sample_size: Sample size to use for bootstrapping. -1 defaults to all samples which is recommended.
    :param confidence_level: Confidence level for result error intervals (0.05 => 95% percentile)
    :param random_seed: Random seed for reproducibility. If None, uses a random seed.
    :return: Dictionary with metrics as keys, each containing 'mean', 'lower', and 'upper' bounds
    """
    if len(y_true) != len(y_pred):
        raise ValueError(f"y_true and y_pred must have same length. Got {len(y_true)} and {len(y_pred)}")

    if sample_size == -1:
        sample_size = len(y_true)

    def default_metrics(y_t, y_p):
        acc, mcc = _calculate_metrics(y_t, y_p)
        return {"accuracy": acc, "mcc": mcc}

    metrics_func = default_metrics

    # Set random seed
    if random_seed is None:
        random_seed = np.random.get_state()[1][0] if np.random.get_state()[1] is not None else 42
    rng = np.random.RandomState(random_seed)

    # Generate all random indices at once
    all_indices = rng.choice(len(y_true), size=(iterations, sample_size), replace=True)

    iteration_results = []
    for indices in all_indices:
        # Sample with replacement
        sampled_y_true = y_true[indices]
        sampled_y_pred = y_pred[indices]

        # Compute metrics for this bootstrap sample
        iteration_result = metrics_func(sampled_y_true, sampled_y_pred)
        iteration_results.append(iteration_result)

    # Process results
    metrics = list(iteration_results[0].keys())
    result_dict = {}
    for metric in metrics:
        all_metric_values = np.array([res[metric] for res in iteration_results], dtype=np.float32)
        mean, lower_bound, upper_bound = _get_mean_and_confidence_bounds(
            values=all_metric_values,
            confidence_level=confidence_level
        )
        result_dict[metric] = {"mean": mean, "lower": lower_bound, "upper": upper_bound}

    return result_dict
```
