## Supplementary material for "*biocentral*: embedding-based protein predictions": pyproject.toml

```
[project]
name = "exotox-reproduction"
version = "1.0.0"
description = "Biocentral API example reproducing the ExoTox predictor"
requires-python = "==3.11.11"
dependencies = [
    "notebook>=7.4.7",
    "numpy==1.26.0",
    "pandas>=2.1.0",
    "scikit-learn==1.2.2",
    "skl2onnx>=1.19.1",
    "ml_dtypes>=0.5.0",
    "onnx>=1.16.0",
    "onnxruntime>=1.23.2",
    "matplotlib>=3.10.7",
    "pandas-stubs==2.3.2.250926",
    "biocentral-api==1.0.0"
]
```
