## Supplementary material for "*biocentral*: embedding-based protein predictions": README.md

# ExoTox Reproduction Example

This example shows how to apply the biocentral_api to reproduce the
[ExoTox predictor](https://doi.org/10.1186/s13040-025-00469-2) and convert it to *ONNX*.

## Required files

All files can be downloaded from the supplementary material of the Exotox paper
[here](https://data.ub.uni-muenchen.de/576/).

The following files are required for the reproduction and conversion:

* [Training sequences](exotox_files/X_train_SST30.fasta)
* [Test sequences](exotox_files/X_test_SST30.fasta)
* [Training labels](exotox_files/y_train_SST30.csv)
* [Test labels](exotox_files/y_test_SST30.csv)
* [ExoTox predictor model checkpoint](exotox_files/sklearn_svcPC20_SST30_CV10_embeddingsProtT5)
* [ExoTox predictor script - showing usage of the predictor](exotox_files/classifying_unknown_proteins.py)

## Installation

To use the notebooks for these examples, make sure you
have [uv](https://docs.astral.sh/uv/getting-started/installation/) installed.

Then run:
```shell
uv venv
source .venv/bin/activate
uv sync
```

## Conversion to ONNX

The [conversion notebook](exotox_conversion.ipynb) shows how to convert the ExoTox predictor to ONNX.
The [onnx file](exotox.onnx) is the result of this conversion. This is the model that is also available
via the `biocentral_api` predict module (`model_name=ExoTox`).

## Reproduction

As a showcase of the `biocentral_api`, the [reproduction notebook](exotox_reproduction.ipynb) shows how to
reproduce the ExoTox predictor with [ProtT5](https://github.com/agemagician/ProtTrans) embeddings.
It compares the reproduced results to the original model and various baselines. The comparison results
are shown in this [barplot](exotox_model_barplot.png).

## Citations

```bibtex
@Article{Krueger2025,
  author    = {Krueger, Tanja and Durmaz, Damla A. and Jimenez-Soto, Luisa F.},
  journal   = {BioData Mining},
  title     = {Exo-Tox: Identifying Exotoxins from secreted bacterial proteins},
  year      = {2025},
  issn      = {1756-0381},
  month     = aug,
  number    = {1},
  volume    = {18},
  doi       = {10.1186/s13040-025-00469-2},
  groups    = {Biocentral_server},
  publisher = {Springer Science and Business Media LLC},
}
```
