## Supplementary figures and images for "*biocentral*: embedding-based protein predictions"

### A0A1B0WLV4.png

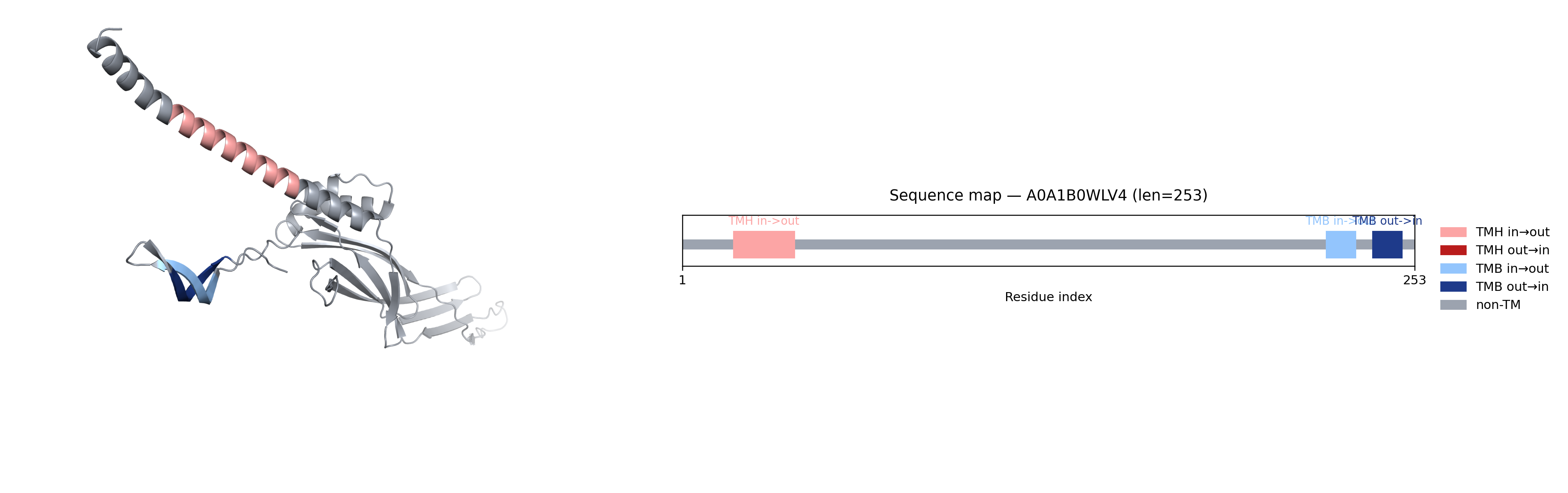

### A0A6B9LG87.png

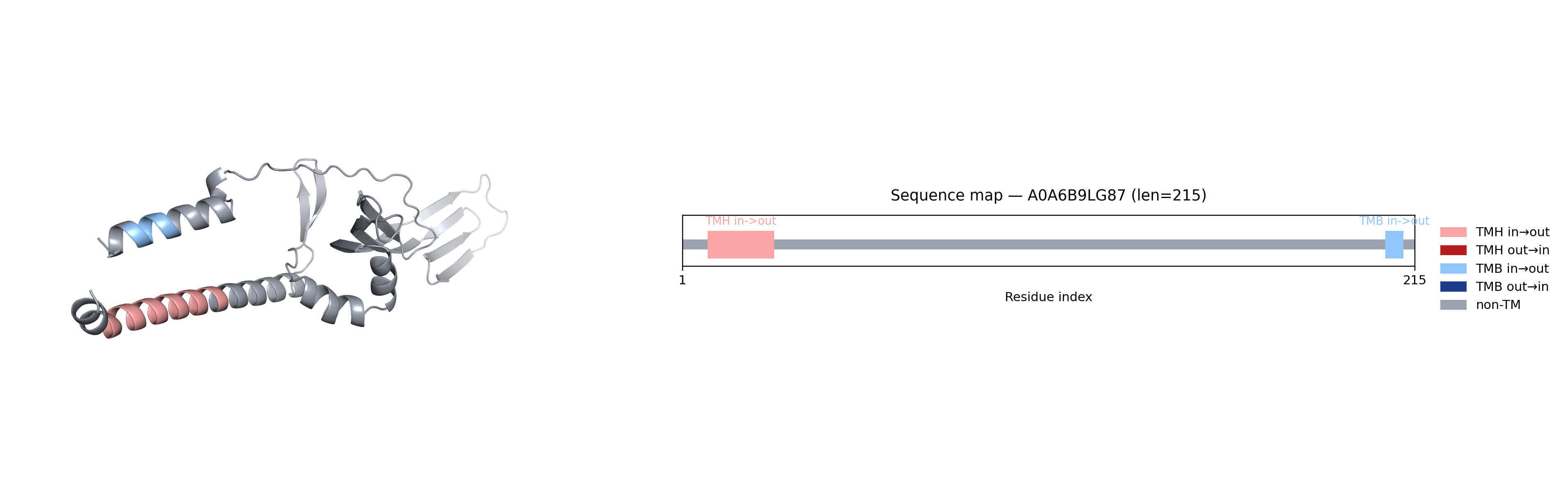

### A0A6J5KMN9.png

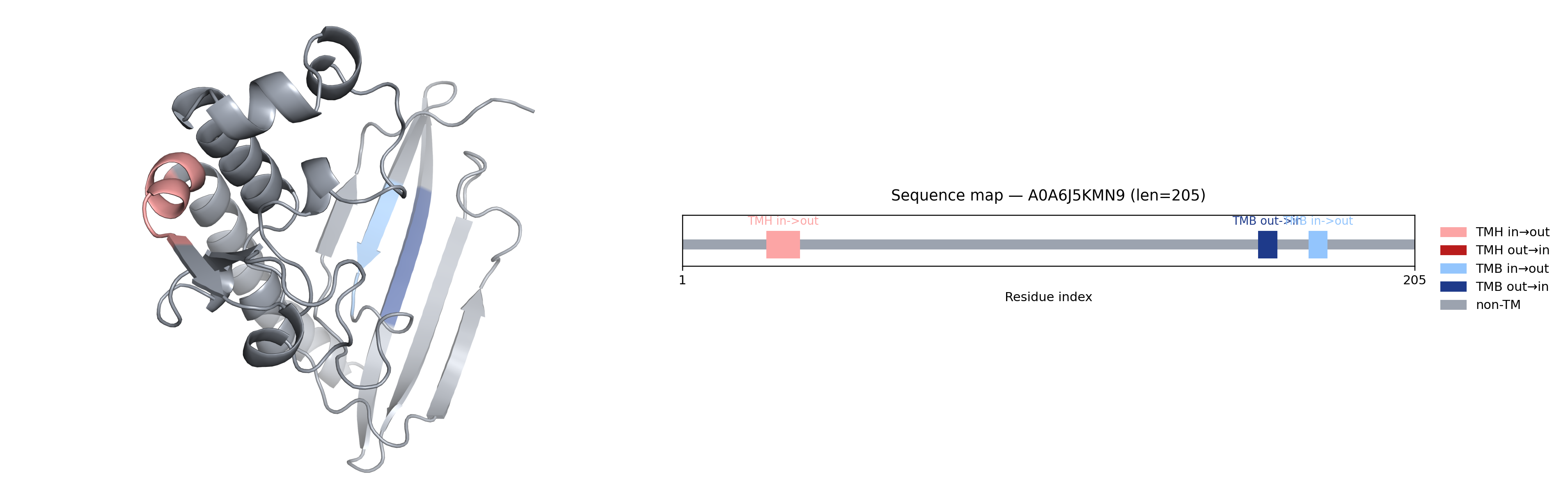

### A0A6J5PUB4.png

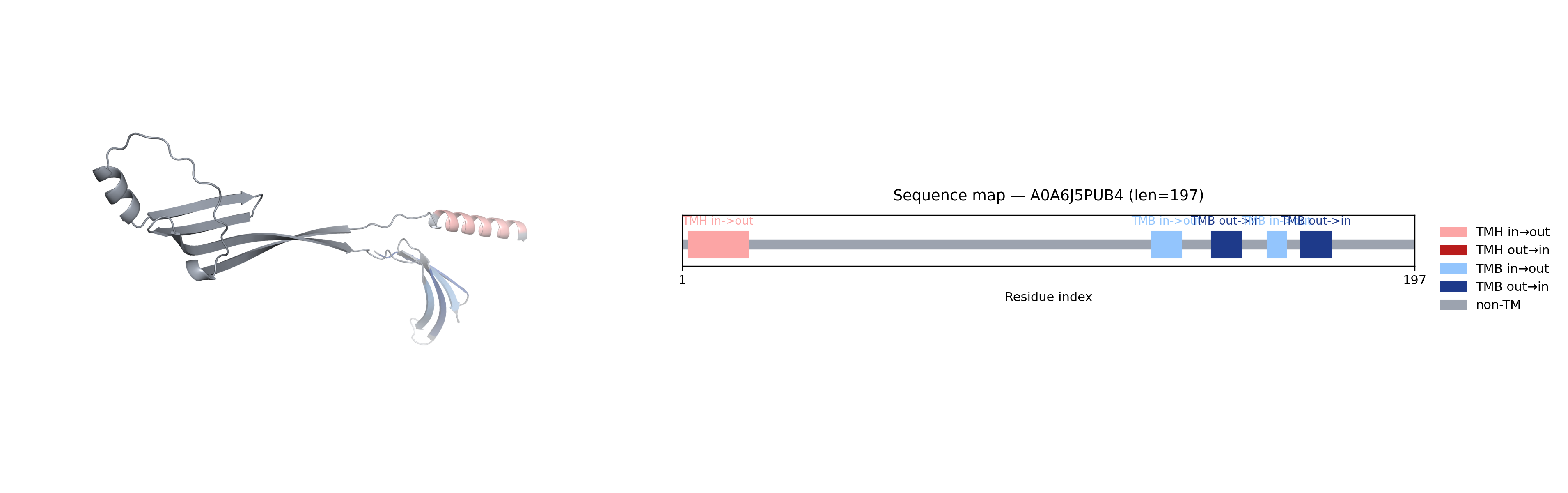

### A0A6J5PY22.png

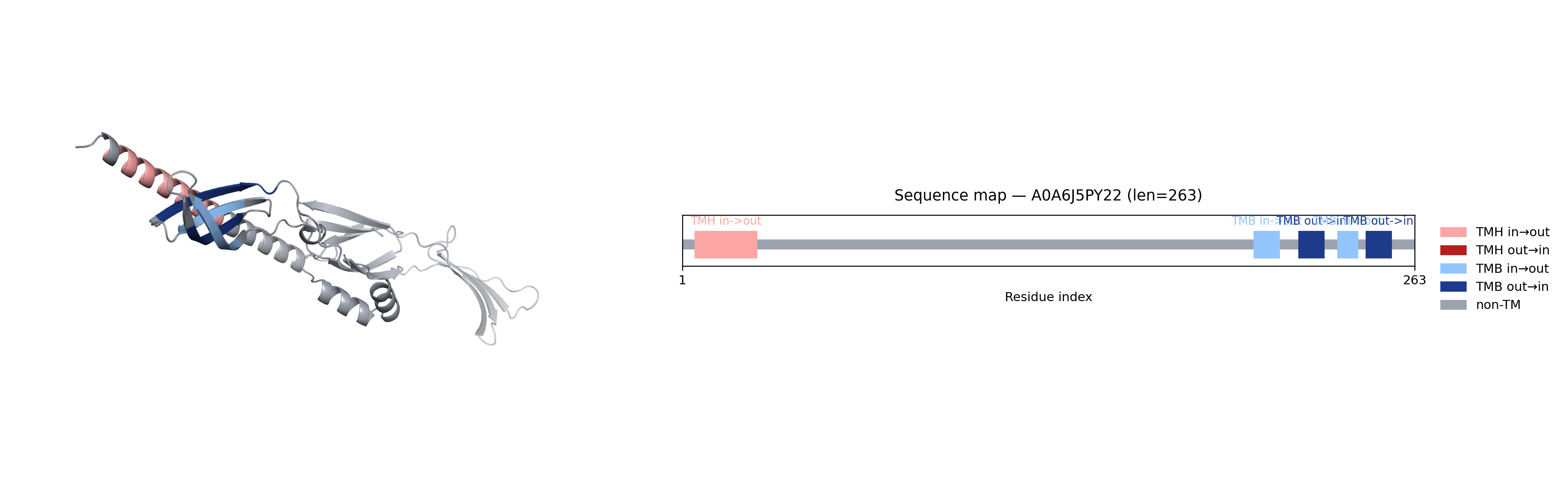

### A0A6J5RKD4.png

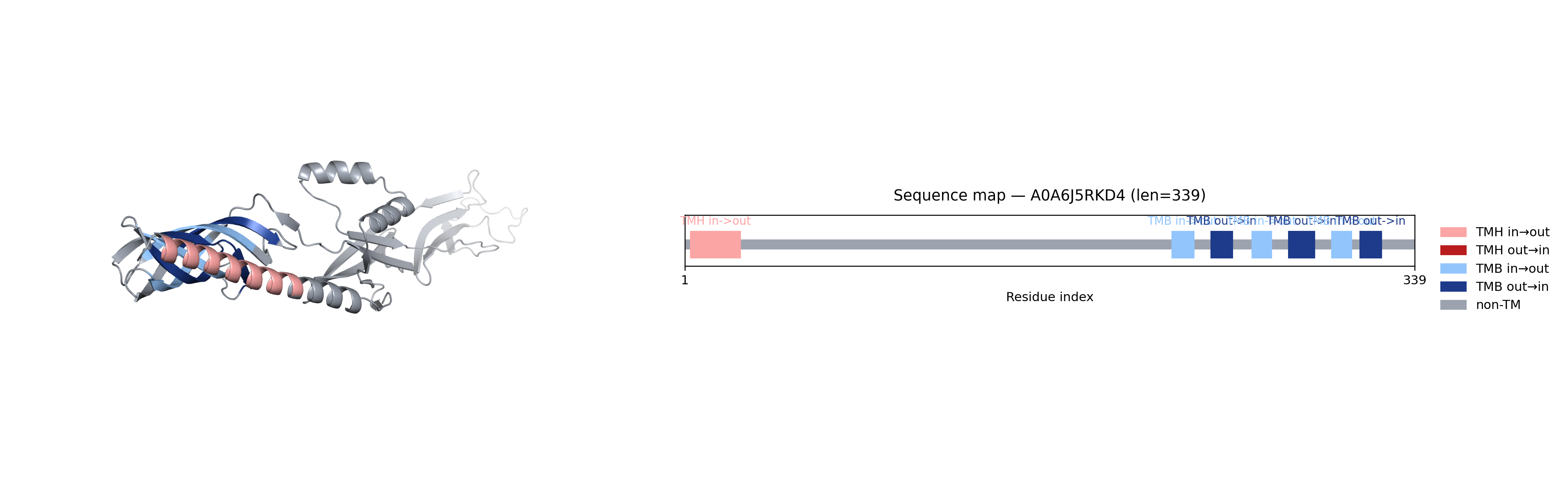

### A0A6J5SVA5.png

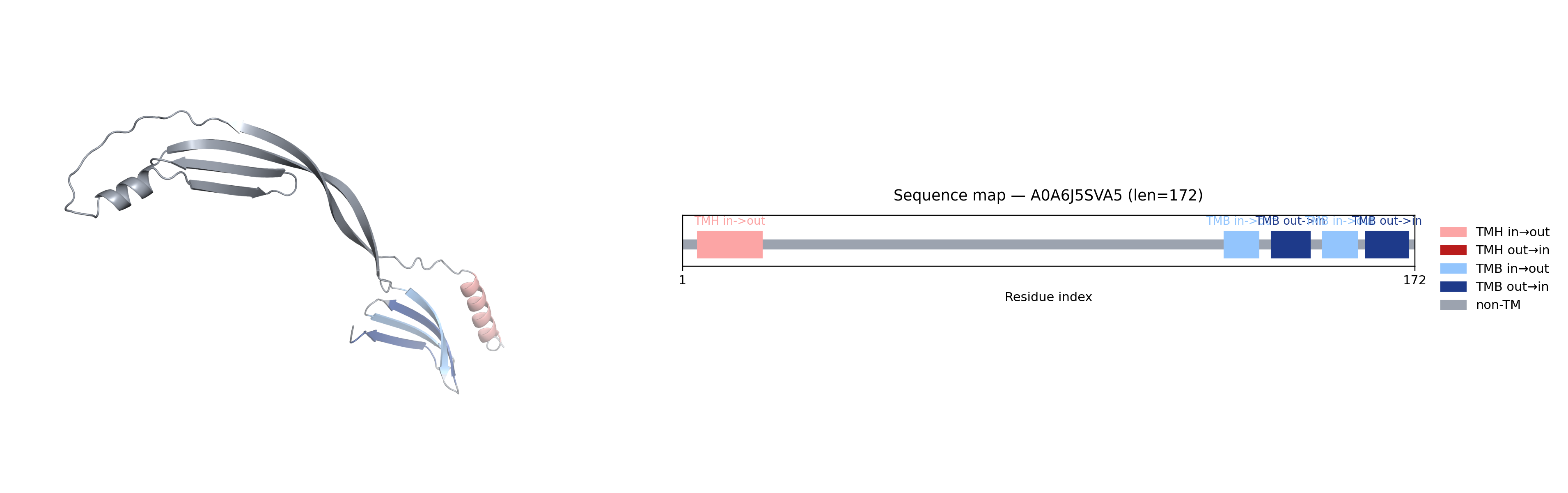

### A0A7D7F863.png

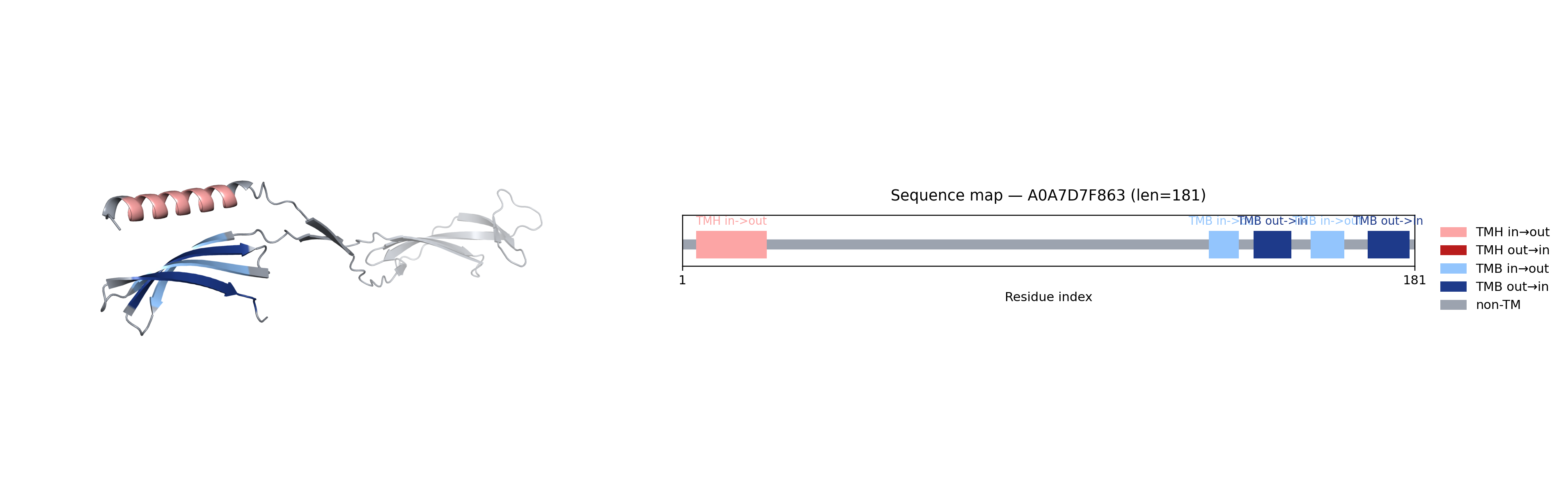

### A0A8S5N8Q4.png

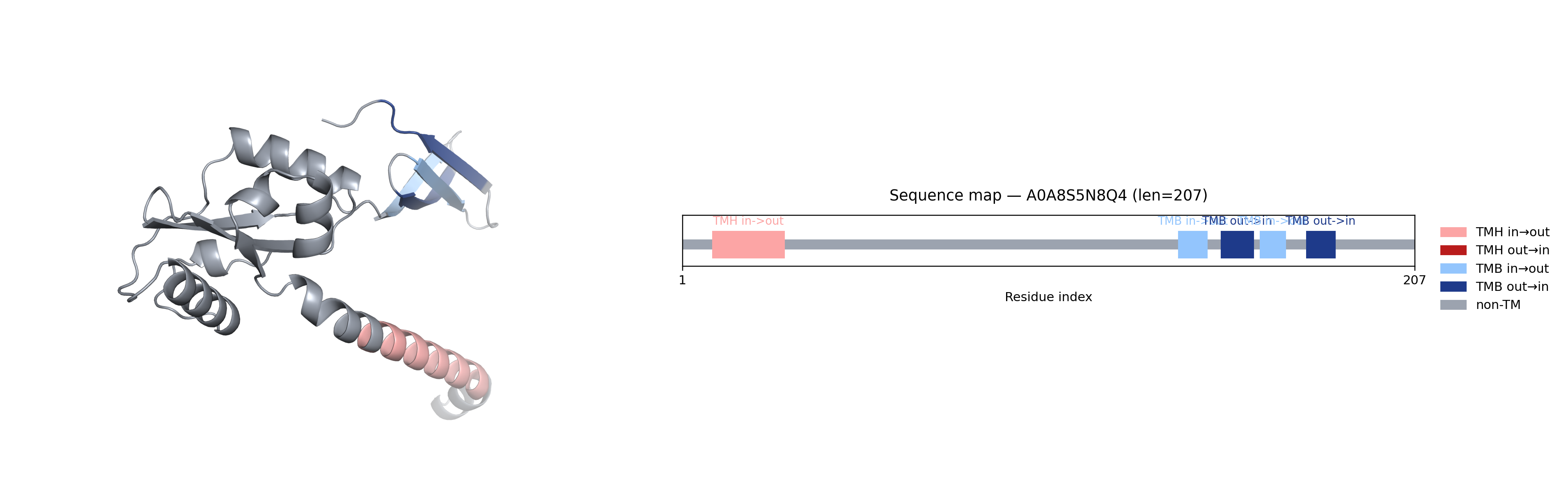

### A0A8S5PJU3.png

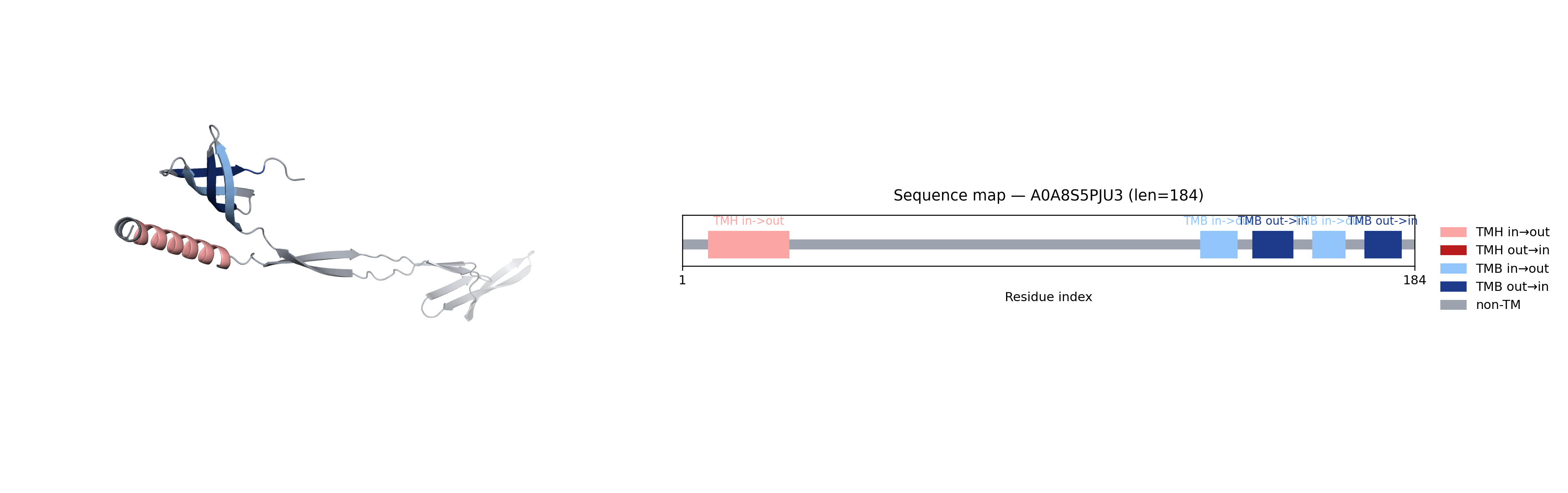

### A0A8S5PWQ1.png

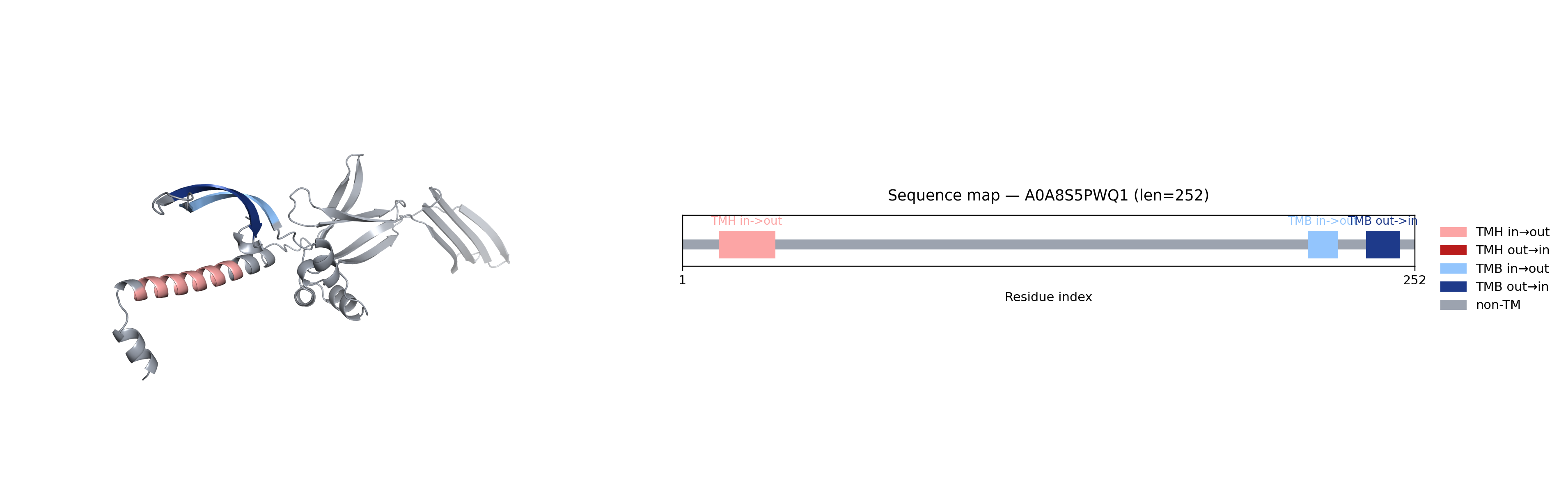

### A0A8S5R786.png

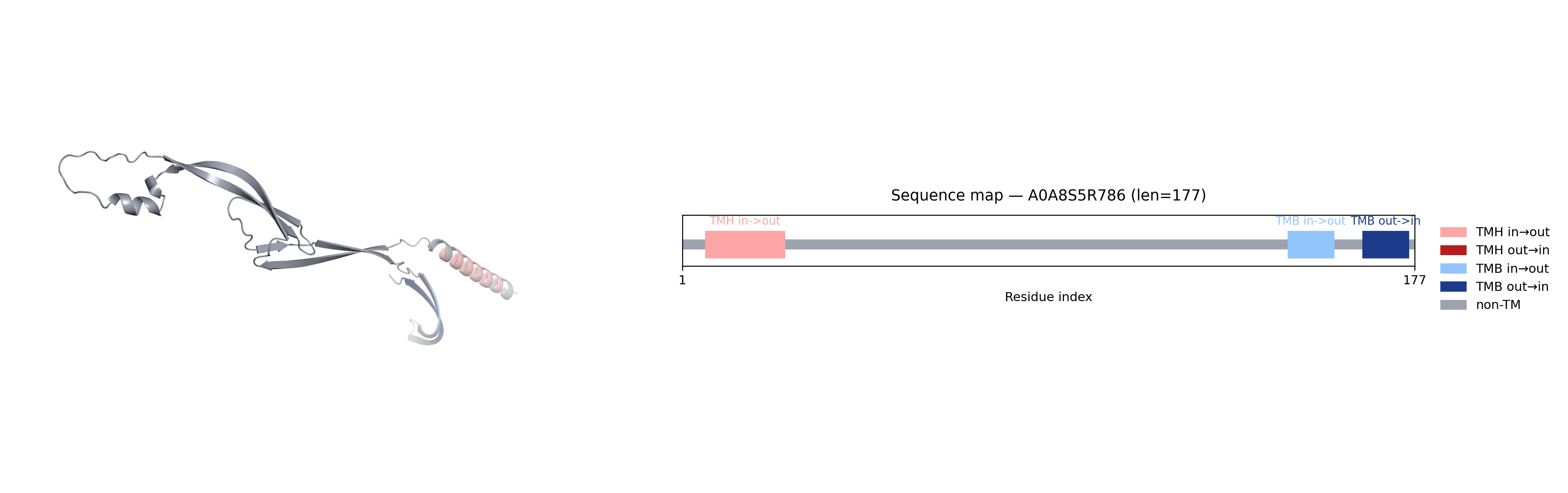

### A0A8S5V0W9.png

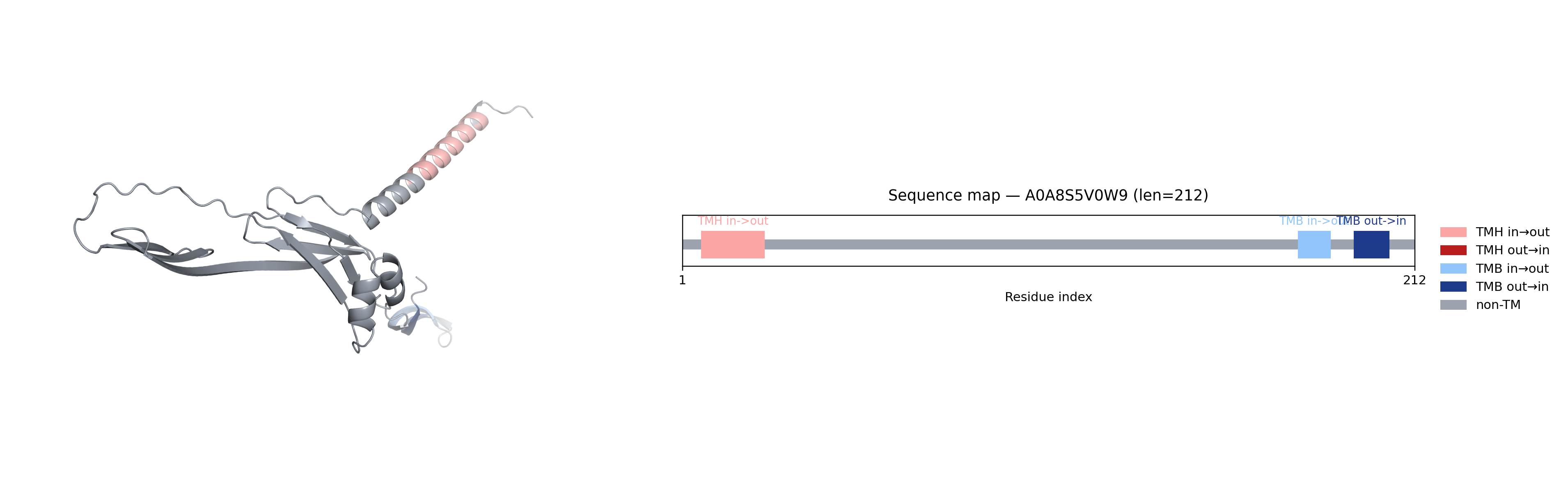

### A0A8S5VRB3.png

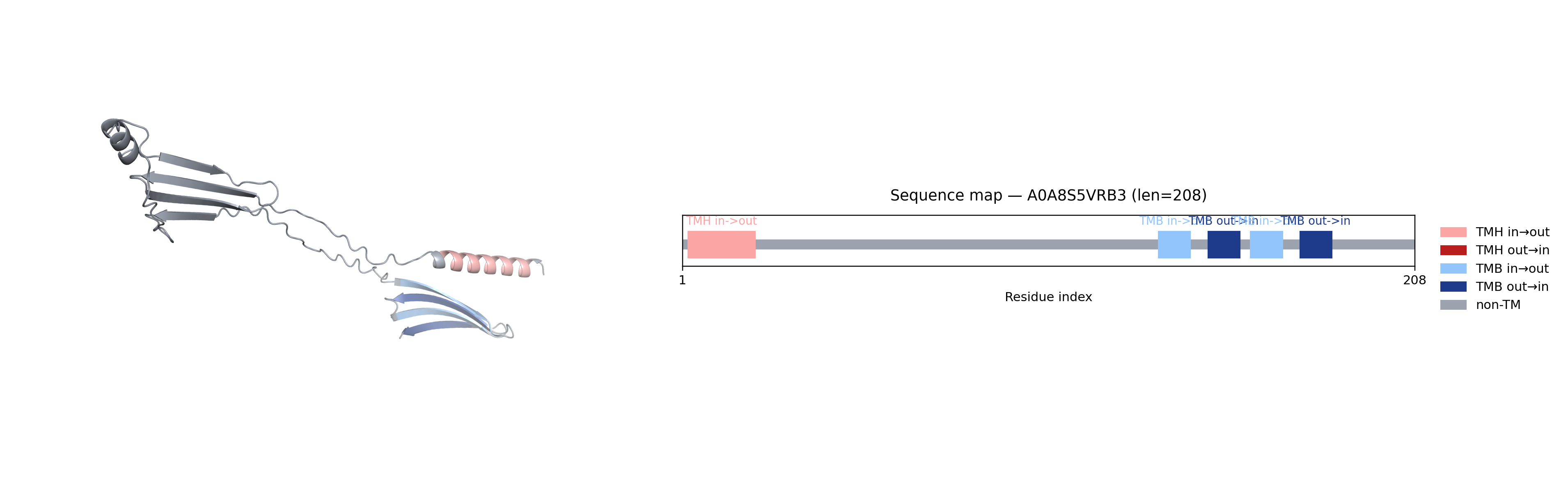

### A0A345BPM7.png

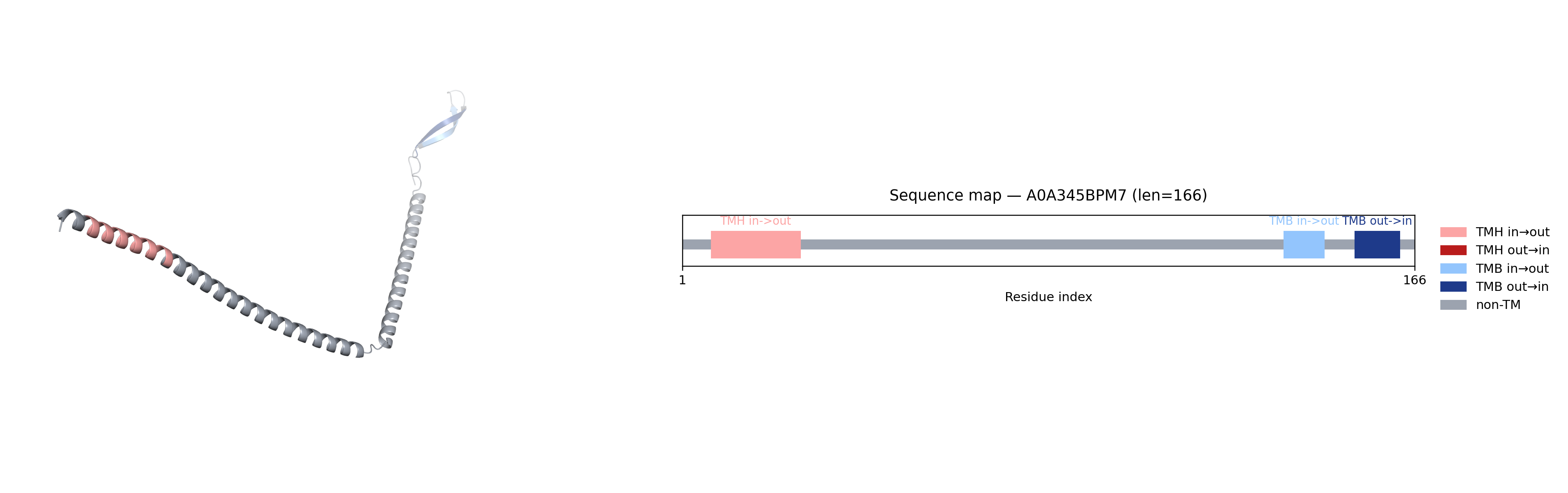

### A0A516M0S4.png

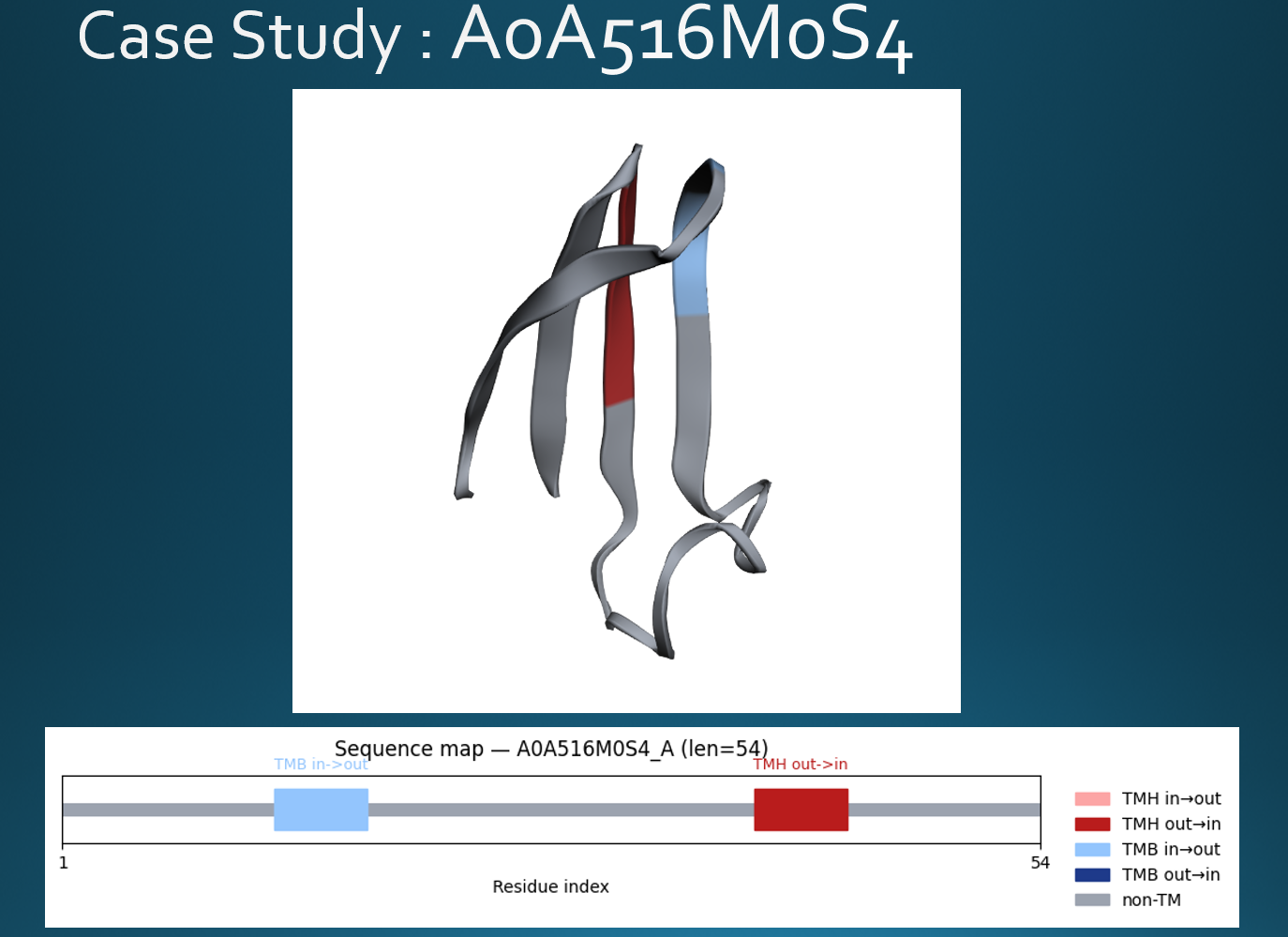

### A0A516M0S4_combined.png

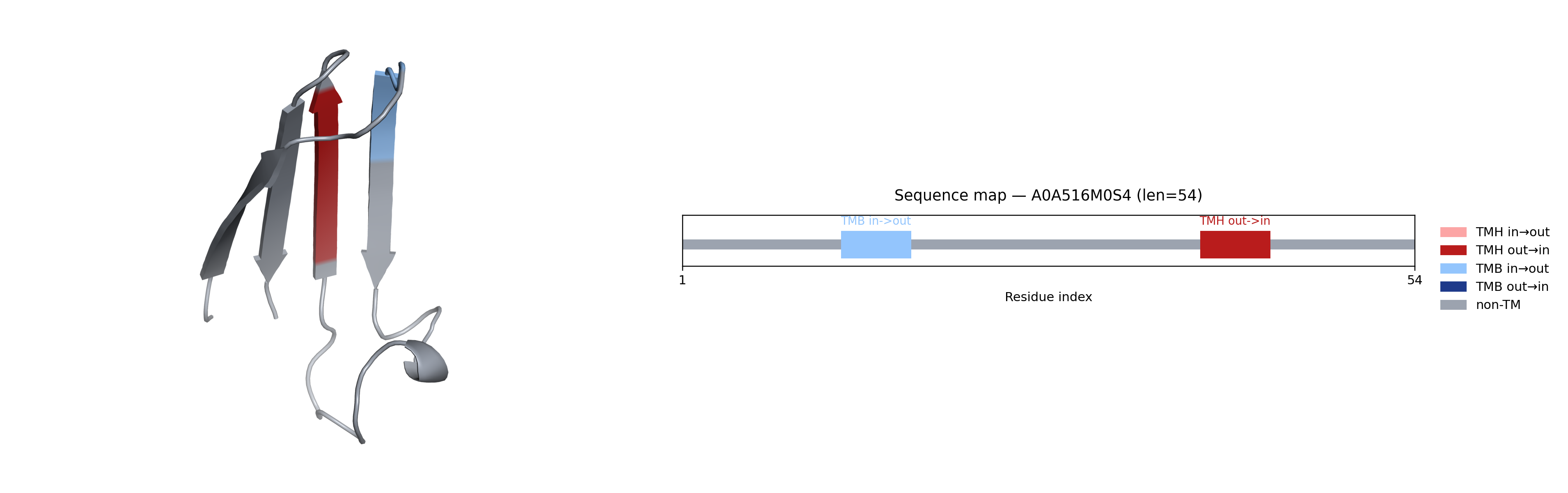

### A0A516M3K9.png

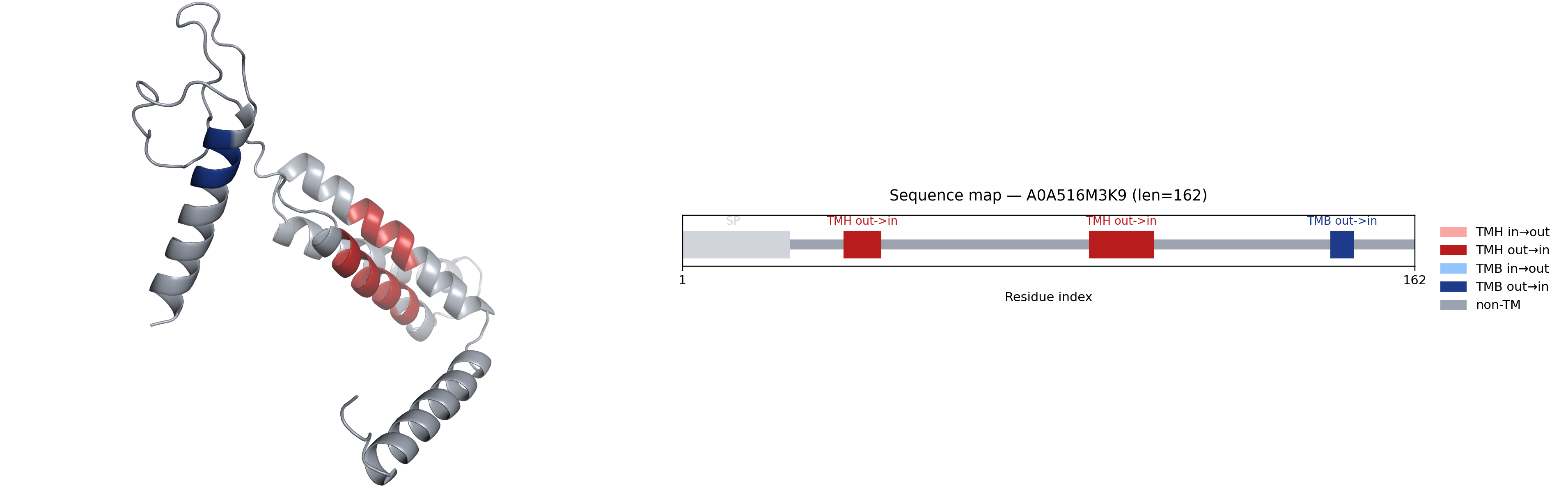

### bfvd_assigned_tm_fraction_by_envelope.png

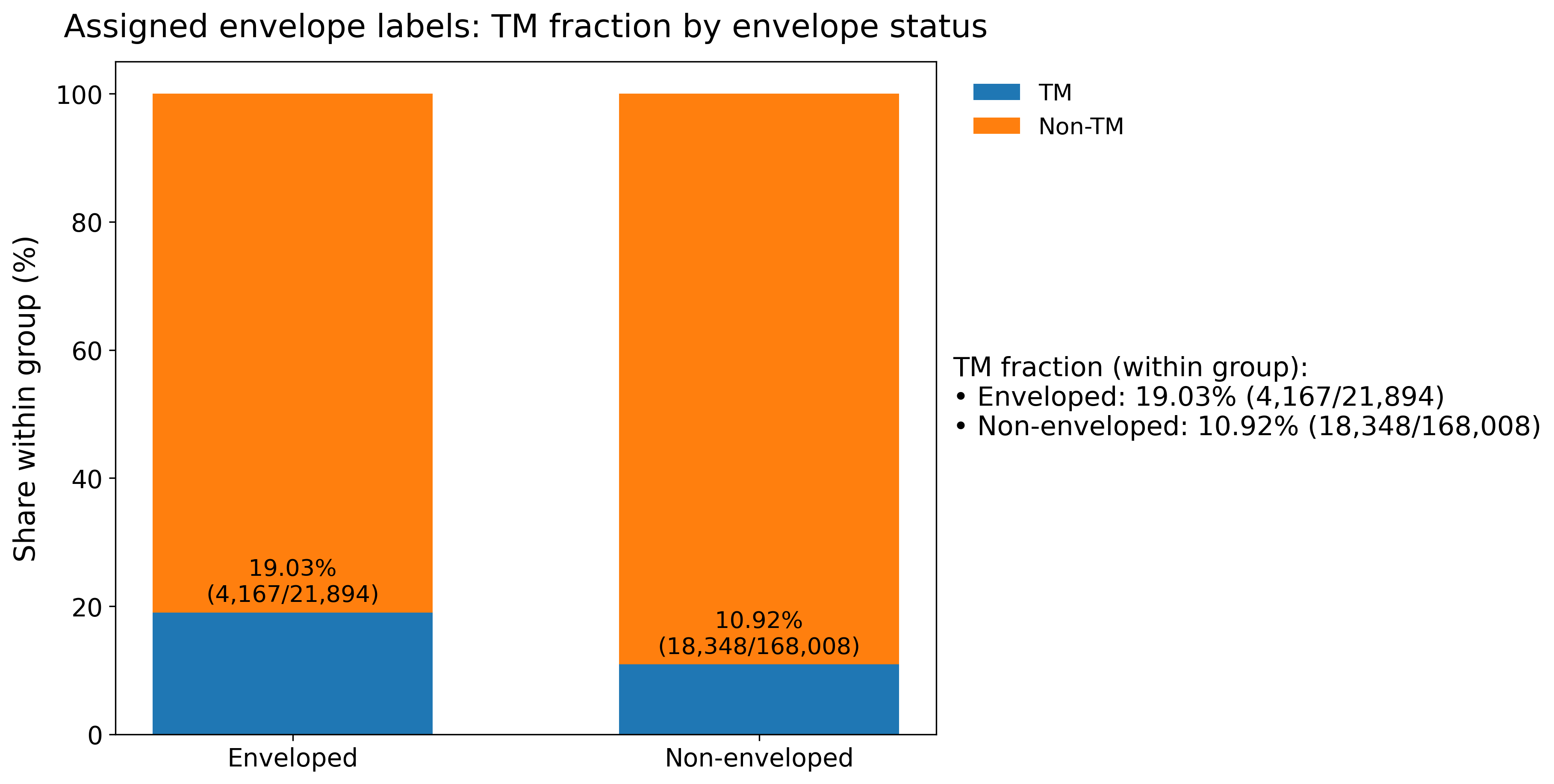

### bfvd_envelope_label_coverage_pie.png

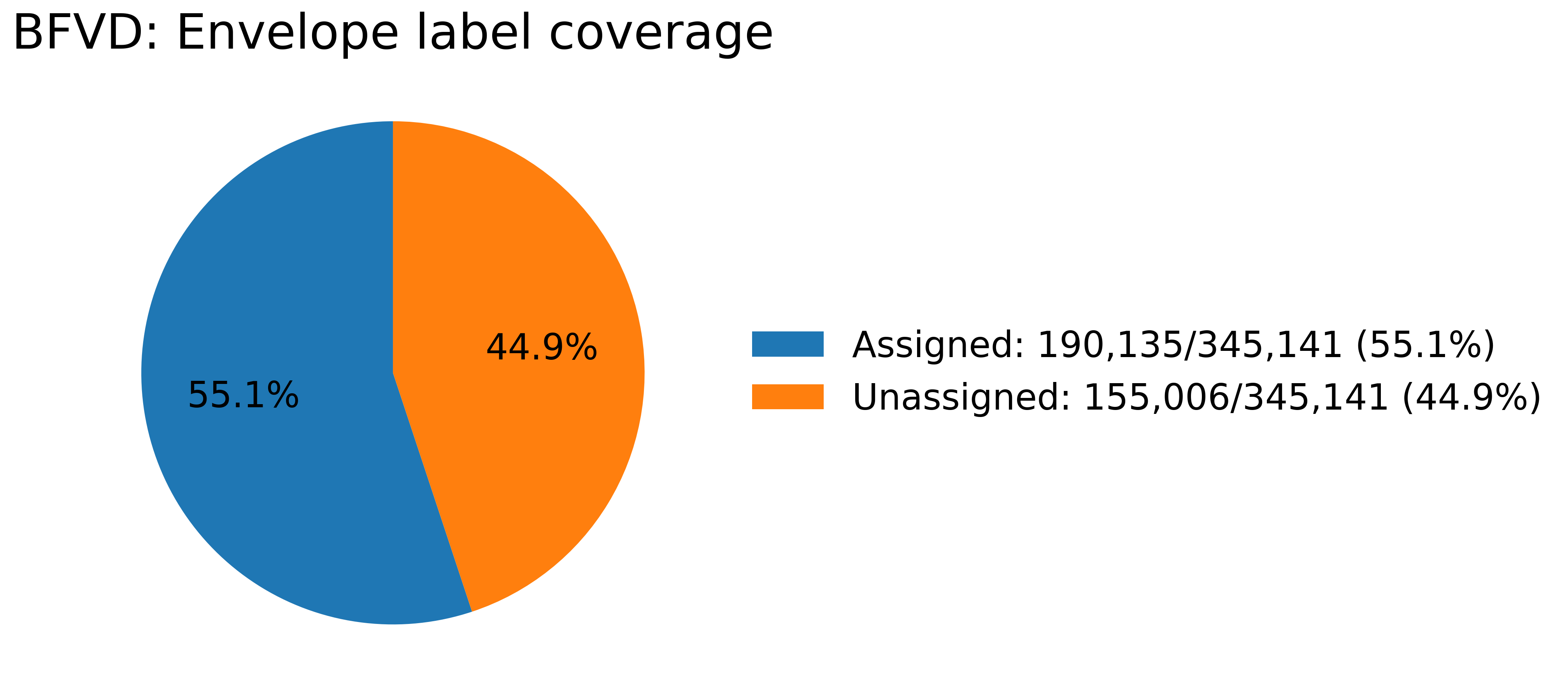

### exotox_model_barplot.png

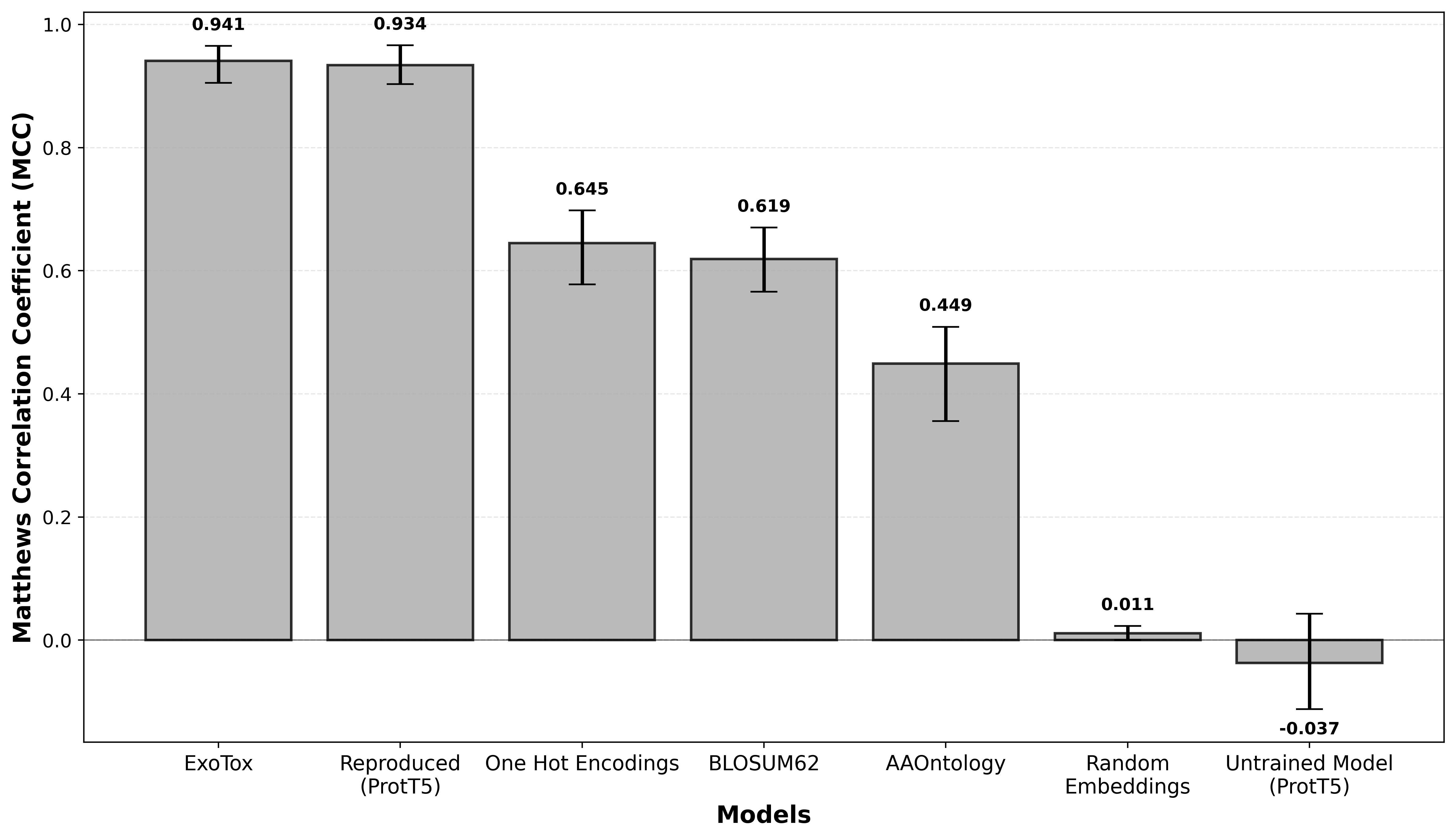

### mermaid-BFVD_Pipeline.png

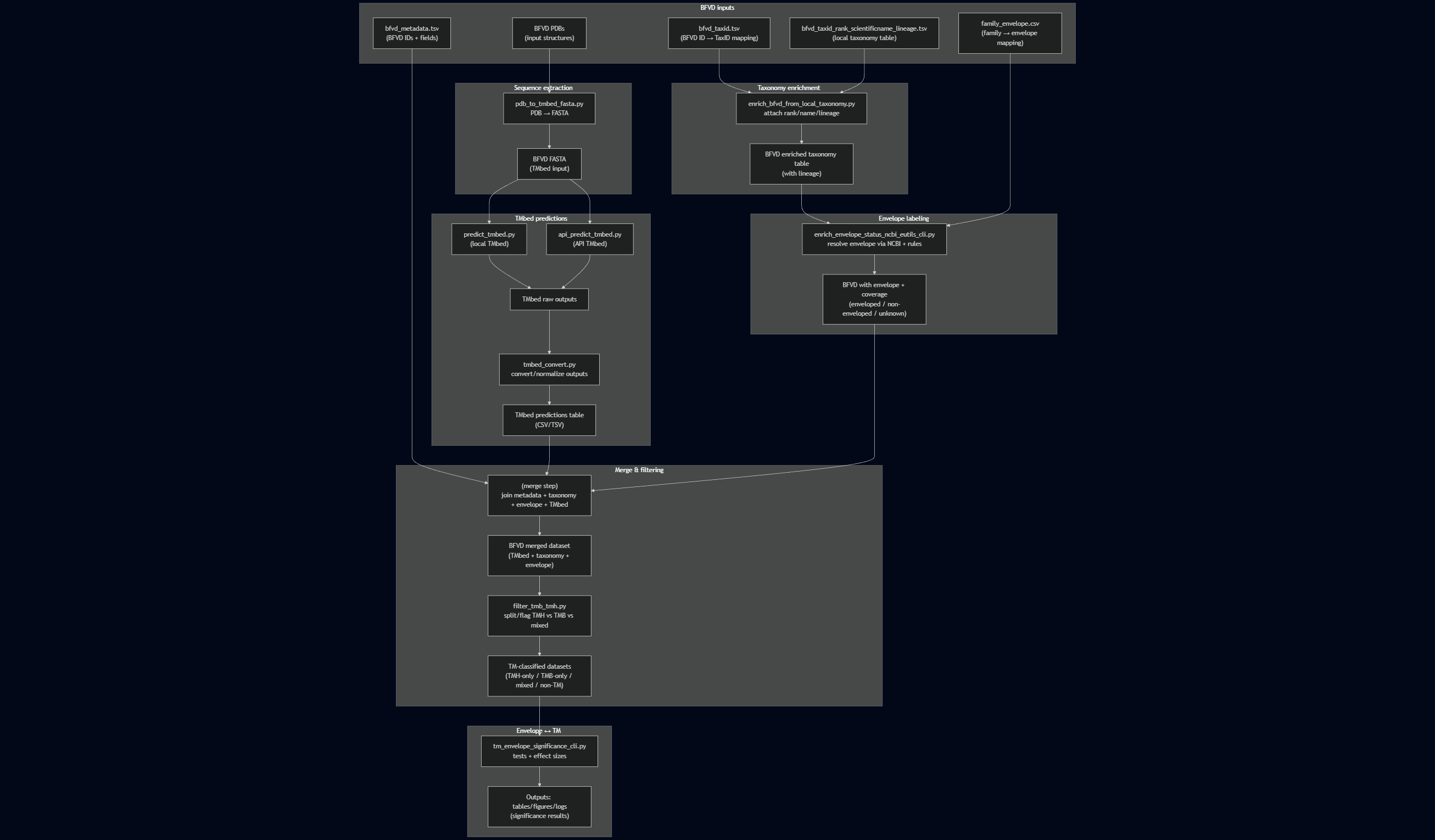

### R9ZZB8.png

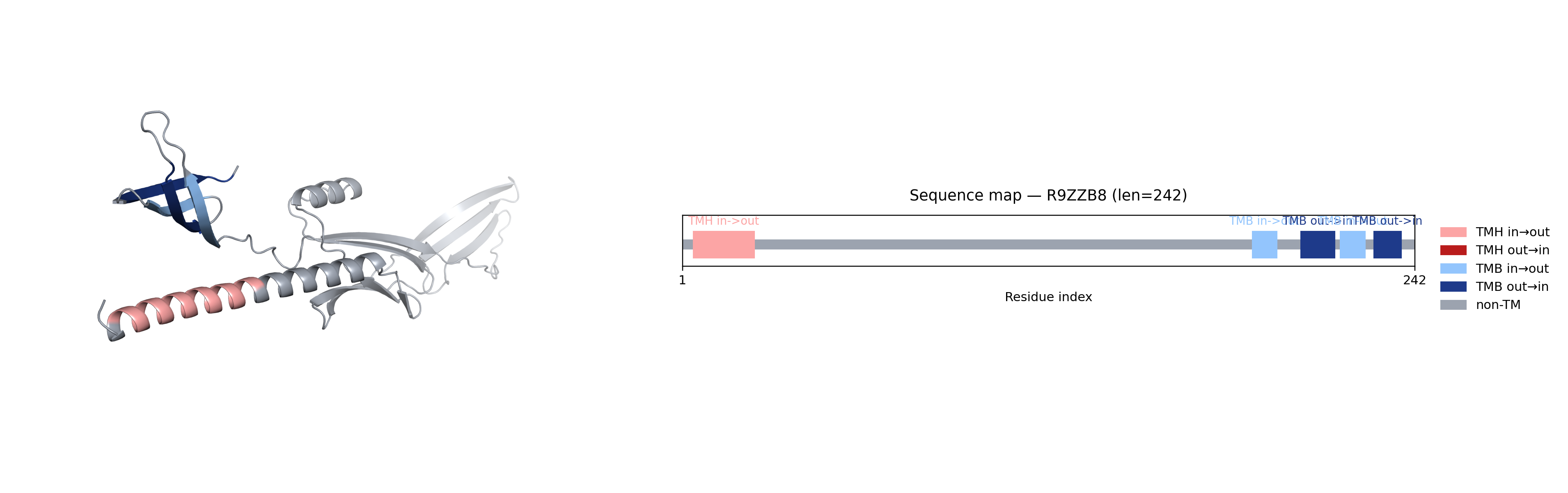

### W8CQR3.png

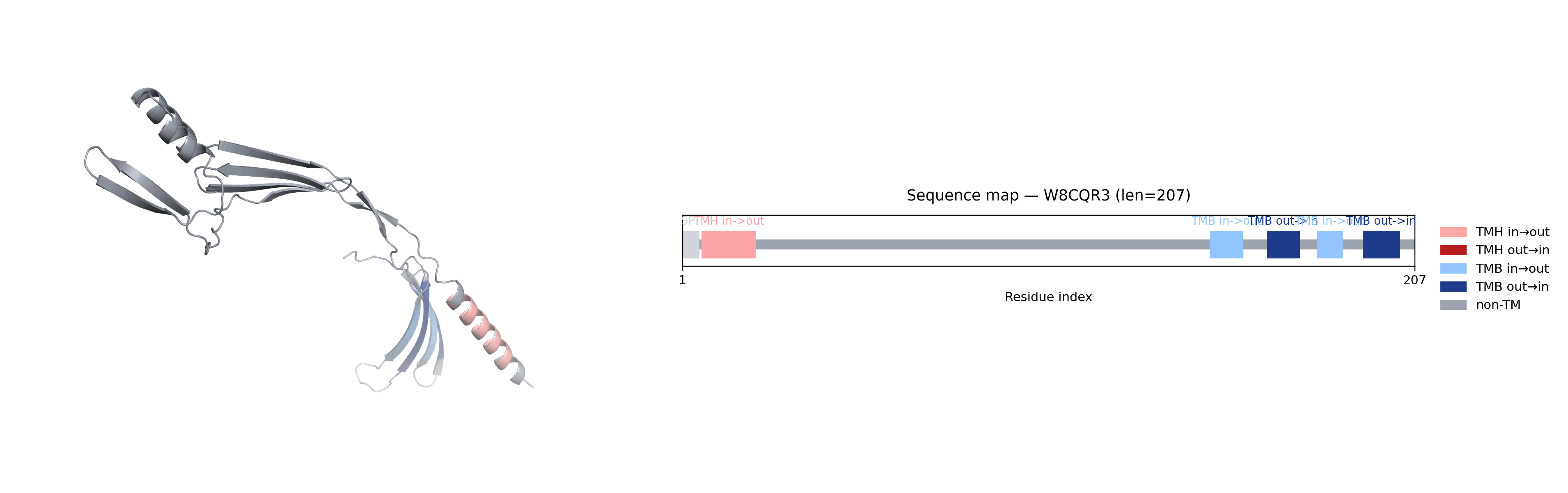
